# PefA, an ALG-2-like Ca^2+^ sensor, regulates ESCRT and autophagy responses to mycobacterial vacuole damage

**DOI:** 10.64898/2026.01.30.702722

**Authors:** Sandra Guallar-Garrido, Angélique Peret, Amado Carreras-Sureda, Céline Michard, Thierry Soldati

## Abstract

Calcium influx is a universal early signal triggering membrane repair pathways, yet how Ca^2+^ sensors coordinate the balance between ESCRT-mediated sealing and autophagy-based responses at damaged endolysosomes and pathogen-containing vacuoles remains unclear. Here we use the *Dictyostelium discoideum*–*Mycobacterium marinum* infection model, a surrogate to study intracellular pathogenesis of *Mycobacterium tuberculosis*, combined with genetic, imaging, and proteomic analyses to identify the penta-EF-hand protein PefA, an ALG-2-like Ca^2+^ sensor, as a Ca^2+^-responsive regulator that orchestrates recruitment of the E3 ubiquitin ligase TrafE, ESCRT components, and the autophagy machinery to damaged membranes. PefA is transcriptionally upregulated and accumulates at the mycobacterial vacuole, promoting timely repair that preserves vacuolar integrity and supports bacterial replication. Loss of PefA impairs ESCRT and autophagy engagement, leading to premature bacterial escape into the cytosol and altered infection outcomes. These findings uncover a conserved Ca^2+^-dependent mechanism linking membrane damage sensing to coordinated repair pathways and shapes host–pathogen interactions, with direct relevance to tuberculosis pathogenesis and host resilience to infection.

**Teaser:** The PefA calcium sensor times membrane repair and autophagy to control vacuole integrity during mycobacterial infection.

## Introduction

Prokaryotic and eukaryotic cells are frequently subjected to membrane injury from mechanical stress, metabolic by-products, pore-forming toxins, and pathogen effectors. In eukaryotic cells, disruption of the plasma membrane or endosomes disturbs ionic homeostasis and can lead to lysosomal membrane permeabilization, protease leakage, and ultimately cell death unless rapid repair mechanisms are activated(1). To counteract such damage, cells initiate a staged response: first containing ion fluxes, then resealing membrane discontinuities, and finally remodeling and clearing damaged membrane remnants (2). Recent research highlights the importance of timing in these processes, with events unfolding over milliseconds, seconds, minutes, and longer periods. Rapid Ca^2+^ influx acts as the earliest and most universal alarm, triggering the accumulation of Ca^2+^-binding proteins at the wound sites, which is crucial for cytoskeleton reorganization, membrane resealing, and the orchestration of downstream effectors. Within this temporal framework, the endo-lysosomal damage response (ELDR) in animal cells provides a unifying perspective that encompasses the plasma membrane, endosomes, lysosomes and bacteria-containing vacuoles. ELDR integrates endosomal sorting complexes required for transport (ESCRT)-dependent scission, autophagy-based repair and sequestration, and lysosomal exocytosis to restore ionic homeostasis and proteostasis, while preventing inflammasome activation and cell death (3).

Early repair phases are dominated by the assembly of the general ESCRT membrane-sealing machinery, particularly ESCRT-III. The ESCRT machinery is recruited to seal nanometer-scale disruptions at the plasma membrane and damaged endolysosomes, often in response to Ca^2+^ sensing or membrane curvature alteration (4–6). At the plasma membrane, Ca^2+^-sensing proteins such as annexins create curvature-stabilizing coats, promote budding, and coordinate Ca^2+^-triggered lysosome exocytosis, which supplies lipids that promote membrane fusion to the wound edge (7, 8). Spatiotemporal phosphoinositide patterns also orchestrate ESCRT assembly and turnover at lesions, translating Ca^2+^ spikes into remodeling kinetics (9). On damaged endosomes, ESCRT rapidly “cap” small holes to prevent larger damage. However, when damage exceeds the threshold for ESCRT repair, autophagy is engaged to repair larger regions or, if repair is not possible, to shift the response toward irreversible degradation of damaged organelles by lysophagy. This process also applies to the capture and restriction of pathogen vacuole remnants and foreign cytosolic bacterial intruders by xenophagy (3, 10). Notably, membrane repair influences cell-fate decisions by dampening pro-death signaling, but persistent or overwhelming damage can drive regulated necrosis or pyroptosis with ESCRT modulating these outcomes by resealing pores or shedding damaged membranes (6, 11). These temporally regulated, Ca^2+^-dependent events form a framework for understanding host-pathogen interactions, where toxins and effectors can perforate or destabilize host membranes (3, 12). However, it remains unclear exactly how Ca^2+^ regulates the balance between damage and repair, and which are the primary Ca^2+^ sensors. Studies in mammalian systems have highlighted the role of the Ca^2+^-binding penta-EF-hand (PEF) protein ALG-2 in linking cytosolic Ca^2+^ elevations to ESCRT recruitment via the adaptor protein ALIX (13–18). Despite this, a conserved pathway that integrates Ca^2+^ sensors acting on endolysosomes and pathogen-modified vacuoles to arbitrate the ESCRT–autophagy balance has yet to be identified. This gap is significant because the decision to repair or remove a damaged compartment determines both host outcome and microbial fate (19, 20). Rapid resealing stabilizes cellular physiology and preserves the pathogen vacuolar niche, allowing for persistence (21). In contrast, failed maintenance or targeted removal exposes bacteria to the cytosol, triggers galectin binding and ubiquitination (22–24), engages xenophagy, and often restricts or eliminates the microbe (19).

Tuberculosis (TB) underscores the stakes of this host–pathogen interplay. *Mycobacterium tuberculosis* (Mtb) remains a leading cause of mortality by a single infectious agent, and its close relative *Mycobacterium marinum* (Mmar) recapitulates key virulence strategies in genetically tractable host model organisms, such as the optically transparent zebrafish vertebrate system (19). After uptake into phagocytes, both Mtb and Mmar remodel the phagosome to avoid acidification and fusion with lysosomes (25). This remodeling involves enzymatic interference with endosomes, most notably through the secreted PI3-phosphatase SapM, which depletes phagosomal PI3P and prevents EEA1-dependent maturation (26), and the tyrosine phosphatase PtpA, which blocks the recruitment of the V-

ATPase proton pump (27). Together, these activities maintain a weakly acidified, maturation-arrested compartment that supports bacterial persistence. Beyond blocking phagosome maturation, mycobacteria actively damage the mycobacteria-containing vacuole (MCV) via secretion of ESX-1 effectors to enable bacterial access to the host cytosol(28). Host cells respond by mobilizing ESCRT components to the damaged MCV, stabilizing the vacuole and preventing catastrophic rupture, thereby limiting uncontrolled bacterial release (29). However, MCV damage simultaneously exposes luminal glycans and membrane-associated danger signals to the cytosol, triggering galectin-binding and ubiquitination dependent pathways that target bacteria toward xenophagic clearance (21, 24, 25, 30). Thus, MCV membrane disruption emerges as a critical decision point, balancing bacterial cytosolic escape against detection and elimination by the host.

Ca^2+^ leakage from damaged Mtb MCV has been shown to recruit the V-ATPase–ATG16L1 complex, independently of the canonical ULK autophagy initiation machinery, including FIP200 and ATG13, thereby driving localized ATG8/LC3 lipidation that limits membrane injury and restricts bacterial replication (31). In parallel, ESCRT is mobilized to cap lesions on Mmar MCVs (29). However, it remains unknown whether Ca^2+^-responsive sensors coordinate the interplay between ESCRT-mediated repair and autophagy at the MCV, or whether similar Ca^2+^-driven mechanisms operate more broadly at damaged intracellular compartments. Clarifying how cells sense and respond to Ca^2+^ dynamics at the damaged MCV is directly relevant to TB pathogenesis and to the development of host-directed therapies aimed at enhancing cellular resilience without stabilizing the pathogen niche. Environmental amoebae such as *Dictyostelium discoideum* (Dd) are professional phagocytes and have emerged as a versatile model for dissecting cell-autonomous defense mechanisms (25). Dd deploys a repertoire of antibacterial mechanisms that closely mirrors those of human phagocytes (32), making it a powerful system to study diverse human pathogens, including mycobacteria (25, 33). Recently, the Dd-Mmar model, a genetically tractable phagocyte-pathogen system in which bacteria replicate within MCVs that can rupture to grant cytosolic access (34, 35), has enabled the dissection of the role of TrafE. TrafE is an evolutionarily conserved TRAF-family E3 ubiquitin ligase that is rapidly recruited to pathogen- or physically damaged endolysosomal membranes and functions at the ESCRT-autophagy interface to promote effective membrane repair and, ultimately, bacteria restriction through xenophagy (36).

Here, we use the Dd – Mmar model to unravel how Ca^2+^, acts through the penta-EF-hand protein (PefA), coordinates ESCRT and autophagy at damaged lysosomes and at the MCV, thereby determining how Ca^2+^ dictates the decision between repair and removal. To address this, we combined assays of Ca^2+^ dynamics, lysosomal pH, proteomic mapping of damage-dependent interactomes for TrafE, ESCRT-III (Vps32), and Atg8a; genetic and pharmacological perturbations of Ca^2+^ (Δ*pefA* and BAPTA-AM); and infection-stage imaging to track mCherry-tagged PefA (mCh-PefA) and quantify recruitment and accumulation of TrafE, Vps32, and Atg8a at the MCV, alongside bacterial compartmentalization and fitness. We find that elevating extracellular Ca^2+^ strengthens cytosolic Ca^2+^ transients and shortens the ESCRT/autophagy response window after lysosomal injury. We identify PefA as a damage-dependent, Ca^2+^-responsive interactor shared among TrafE, Vps32, and Atg8 upon sterile damage induction. During infection, the Ca^2+^ sensor PefA accumulates at the MCV and is required for the recruitment and accumulation of TrafE, Vps32, and Atg8a. In the absence of PefA, this recruitment is impaired, leading to increased cytosolic Mmar exposure.

## Results

### Extracellular Ca^2+^ regulates host fitness and mycobacterial microcolonies

Ca^2+^ is considered an early chemical signal released upon membrane damage, important for the recruitment of host repair machineries (8, 37) . To test whether elevating extracellular Ca^2+^ (i.e. adding an extra 5 mM) modifies Dd cell response to stress, we compared the impact of Ca^2+^ addition in Dd culture medium (HL5c versus HL5c + Ca^2+^). Ca^2+^addition noticeably improved Dd growth (Fig. 1A and fig. S1A). Automated quantification confirmed faster replication in Ca^2+^ supplementation (Fig. 1B and fig. S1B), resulting in a significantly higher integrated growth (area under the curve, a.u.c.) (Fig. 1C). Interestingly, Ca^2+^ supplementation in mycobacteria cultures, either in 7H9 (fig. S1C) or HL5c (fig. S1D), resulted in enhanced Mmar growth at later time points.

**Fig. 1.**
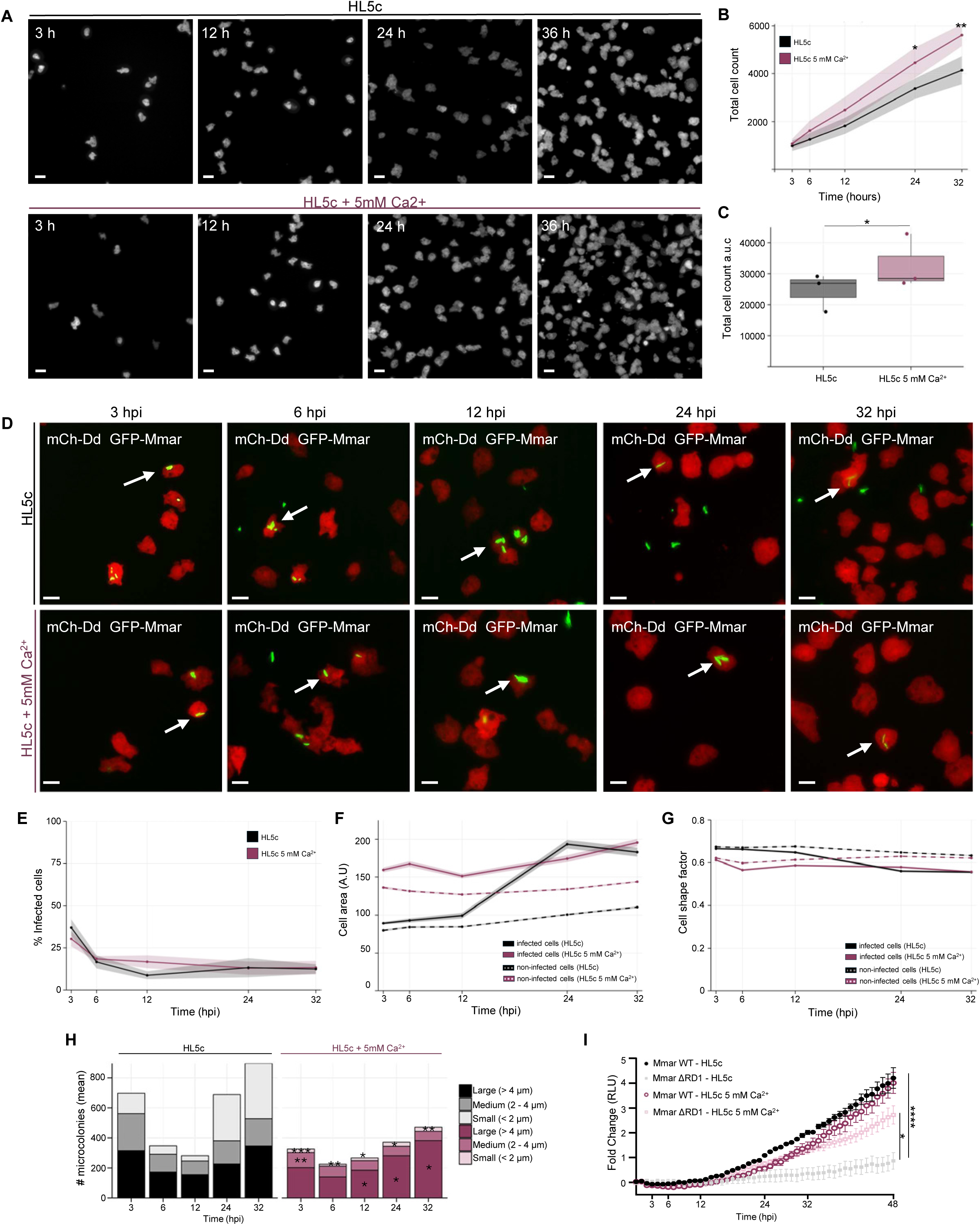
Extracellular Ca^2+^ reshapes the dynamics of Mmar infection in Dd. (**A**) Representative confocal images of uninfected Dd cultures grown in HL5c (top row) or HL5c supplemented with 5 mM Ca^2+^ (bottom row) at 3 h, 12 h, 24 h and 36 h after seeding. Scale bar, 10 µm (**B**) Total Dd cell counts over time in HL5c (black) and HL5c + 5 mM Ca^2+^ (magenta); shaded areas indicate SEM. Quantification was performed per cell ( >900 cells per condition: mean ± SEM, n=3; N=3).Statistical comparisons at each time point were performed using a two-sided Welch’s t-test (*p≤0.05, **p≤0.01). (**C**) Area under the curve (a.u.c.) of time course shown in (B), two-sided Wilcoxon rank-sum test (Mann–Whitney U) test. *p < 0.05, **p < 0.01 (**D**) Representative images of cytosolic mCherry-expressing Dd (red) infected with GFP-expressing Mmar (green) and cultured in HL5c (top row) or HL5c + 5 mM Ca^2+^ (bottom row) at the indicated times post-infection. White arrows indicate infected cells. Scale bar, 10 µm (**E-G)** Quantification of D by automated image analysis of Dd infection kinetics in HL5c (black) and HL5c + 5 mM Ca^2+^ (magenta) at the indicated times post-infection (>2000 cells per condition; mean ± SEM, n=3; N=3), showing (**E)** proportion of infected cells; (**F**) Dd area, and (**G**) Dd shape factor of infected cells (solid lines) and non-infected cells (dashed lines). (**H-J**) Automated image analysis of mycobacterial microcolonies over time in HL5c (black) and HL5c + 5 mM Ca^2+^ (magenta) ( >675microcolonies per condition (mean ± SEM, n=3; N=3). (**H**) Number and size distribution of mycobacterial microcolonies, classified as small (<2 µm), medium (2–4 µm), or large (>4 µm). Statistical comparisons between conditions were performed on per-well microcolony counts for each time point and size using the Wilcoxon rank-sum test, with Benjamini–Hochberg correction for multiple testing. *p < 0.05, **p < 0.01 (**I**) Dd WT cells were infected with bioluminescent Mmar WT cultured in HL5c (black) or HL5c + 5 mM Ca^2+^ (grey), or with Mmar ΔRD1 cultured in HL5c (magenta) or HL5c + 5 mM Ca^2+^ (pink). Bioluminescence was monitored for 48 hours (mean fold change ± SEM, n=3, N=3). Statistical analysis was performed using one-way ANOVA followed by Turkey’s post-hoc test, *p≤0.05, **p≤0.01, ***p≤0.005, ****p≤0.0001). hpi, hours post-infection; mCh, mCherry; A.U, arbitrary unit; RLU, relative luminescence unit.

To connect this growth effect to infection, we infected Dd with Mmar. Virulent strains use the ESX-1 system (encoded in the genomic Region of Difference 1, RD1) to perforate the MCV and access the cytosol, while the ΔRD1 strain lacks this activity, remains contained inside its MCV, and is thereby attenuated. Because Ca^2+^ is an early injury signal that coordinates host repair machineries, we asked whether raising extracellular Ca^2+^ modifies how cells respond to ESX-1-dependent damage. We therefore infected cells in HL5c or HL5c plus Ca^2+^ and monitored infection burden, microcolony dynamics, and cell fitness over time (Fig. 1D to I). Overall, Ca^2+^ did not alter the fraction of infected cells over the course of infection (Fig. 1D and E) nor the percentage of phagocytosed bacteria (Fig. S1E, S1F and S1G). In addition, we computed a shape factor (where 0 = non-rounded/irregular and 1 = perfect circle) as an indirect index of amoeba cell fitness. Infected cells cultured in Ca^2+^-supplemented medium displayed a larger cell area (Fig. 1F) and lower shape factor (Fig. 1G), consistent with fitter, more polarized motile cells. We next quantified both the number and size of intracellular mycobacterial microcolonies in infected cells to assess whether Ca^2+^ impacts bacterial fate. Time-resolved analyses revealed that Dd cells infected with Mmar and cultured in HL5c harbored a higher number of intracellular microcolonies than cells maintained in Ca^2+^-supplemented medium (Fig. 1H). Size-resolved analyses further showed that Ca^2+^ supplementation was associated with a significantly lower number of small microcolonies at all time points analyzed. Moreover, from approximately 12 hpi onward, Ca^2+^ supplementation led to a significant increase in the number of large microcolonies compared with HL5c alone (Fig. 1H and Fig. S1G), suggesting a distinct infection trajectory. These observations raised the possibility that Ca^2+^ selectively enhances intracellular replication depending on mycobacterial pathogenicity and vacuole integrity. To assess the impact of Ca^2+^ on pathogenic versus non-pathogenic mycobacteria, we monitored intracellular growth of lux-expressing Mmar WT and ΔRD1 strains using plate-reader assays (Fig. 1I). While Mmar WT growth in Dd WT cells was unaffected by Ca^2+^ supplementation up to 48 hpi, intracellular growth of the ΔRD1 mutant was significantly enhanced in the presence of Ca^2+^ (Fig. 1I). These results indicate that Mmar WT, which induces MCV damage and later escapes to the cytosol, is initially less affected by Ca^2+^ supplementation than Mmar ΔRD1, which remains confined in its MCV, highlighting a compartment- and damage-dependent impact of Ca^2+^ on intracellular mycobacterial fate. Overall, these data indicate that extracellular Ca^2+^ enhances both Dd and mycobacteria fitness without altering the initial phagocytic uptake and infection efficiency. However, the observed shifts in microcolony morphology and the Ca^2+^-dependent phenotype of the Mmar ΔRD1 mutant prompted further mechanistic studies of Ca^2+^-dependent damage-repair pathways, to determine whether mycobacteria are more effectively contained within the vacuole upon Ca^2+^ supplementation, thereby limiting their growth, or are cleared more rapidly by xenophagy upon escape to the cytosol.

### Ca^2+^ signaling promotes lysosomal repair and cell survival

Because Ca^2+^ is an early signal following membrane damage and a key trigger for recruitment of host repair machineries (8, we used L-leucyl-L-leucine methyl ester (LLOMe)–induced endolysosomal injury as a paradigm to examine Ca^2+^-dependent membrane damage–repair pathways. LLOMe has been shown to reproduces key features of EsxA-mediated damage to the MCV {Perret, 2026 #2007, 37), but it is simpler to use, and induces synchronous, homogeneous, and reversible endolysosomal injury. Because it elicits conserved host responses, including Ca^2+^ fluxes, it enables direct testing of how extracellular Ca^2+^ modulates cellular tolerance to membrane stress. We therefore assessed the impact of extracellular Ca^2+^ on lysosomal function using LysoSensor, a pH-dependent probe that accumulates in acidic compartments, and quantified LysoSensor response following LLOMe-induced damage (Fig. 2A–D). Representative images show that Ca^2+^-supplemented cells showed higher resistance to LysoSensor leakage and faster recovery of LysoSensor fluorescence after LLOMe addition, consistent with preserved acidity or faster re-acidification (Fig. 2A). Quantitative analyses revealed that the fraction of LysoSensor-positive cells remained stable over time in the presence of Ca^2+^ with only minor fluctuations (Fig 2B) and that the area of LysoSensor-positive structures was better maintained in the presence of Ca^2+^ (Fig. 2C). In parallel, the shape factor re-normalized more rapidly with Ca^2+^, indicating improved cellular fitness (Fig. 2D). Because LysoSensor reports lumenal pH and the persistence of puncta reflects compartment integrity, these data support a model in which Ca^2+^-dependent repair and re-acidification enhance endolysosomal resilience upon damage.

**Fig. 2.**
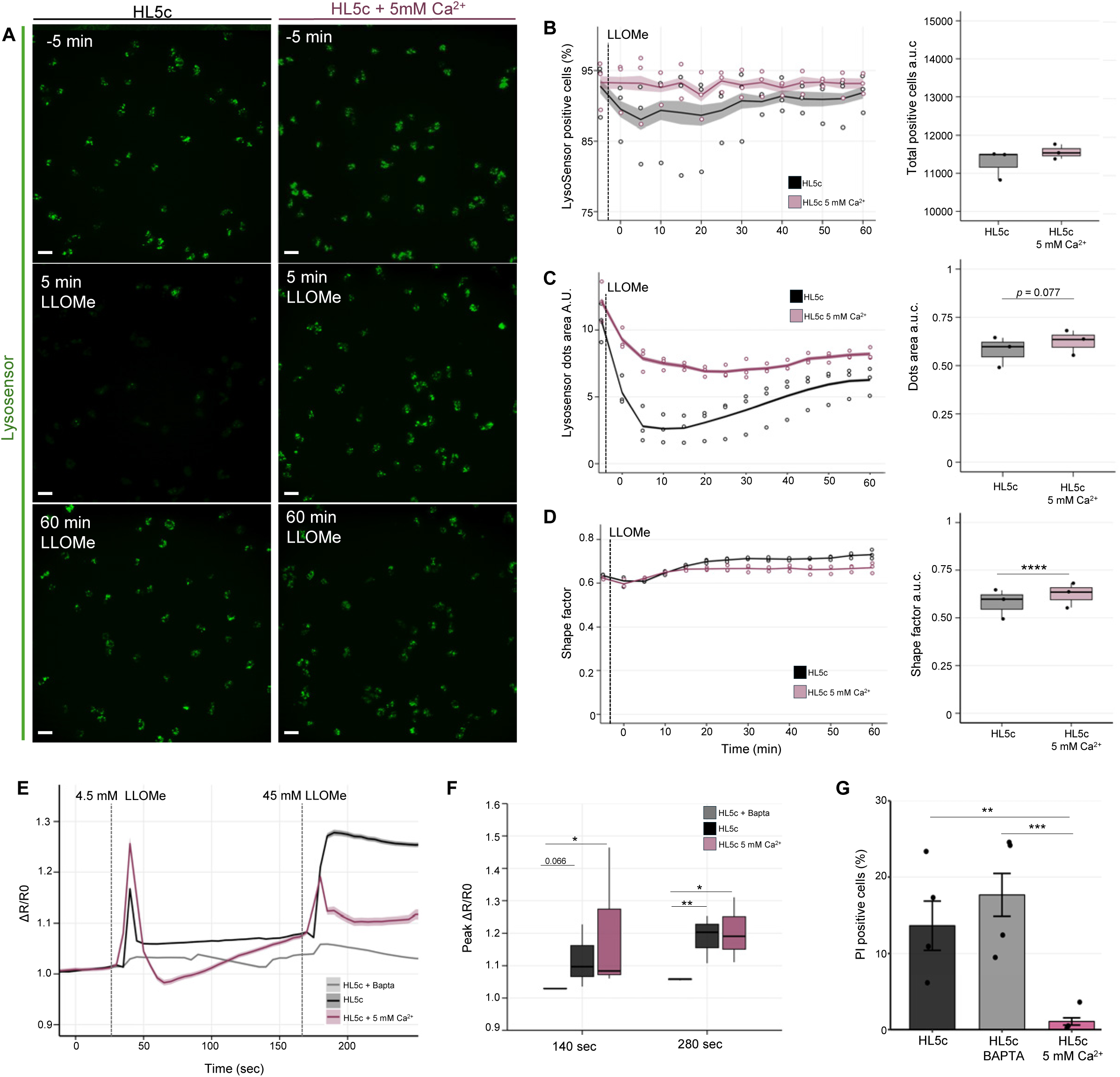
Extracellular Ca^2+^ enhances cell recovery upon sterile damage. (**A**) Time-lapse images of Dd cells cultured in HL5c or HL5c + 5 mM Ca^2+^ , stained with LysoSensor and treated with 4.5 mM of LLOMe for 60 min. Scale bar, 10 μm. (**B–D**) Quantification of the LysoSensor signal shown in (A), (>1300 cells per condition, mean ± SEM, n=3, N=3) (**B**) Image-based analyses of A showing (**B**) the proportion of LysoSensor-positive cells over 60 min and the corresponding area under the curve (a.u.c.), (**C**) the LysoSensor dot area over time and a.u.c, (**D**) the cell shape factor over time and a.u.c. Statistical comparisons were performed using two-sided Welch’s t-test, *p < 0.05, ****p < 0.0001. (**E**) Normalized FRET ratio of Dd cells expressing the YC3.60 Ca^2+^ FRET sensor cultured in HL5c, HL5c + 5 mM Ca^2+^ or HL5c + BAPTA, imaged every 5 s. Cells were sequentially exposed to 4.5 mM and 45 mM LLOMe at the times indicated by dashed vertical lines. (**F**) Boxplots from E summarizing the Ca^2+^ response amplitudes following the first (4.5 mM) and second (45 mM) LLOMe additions (mean ± SEM, n ≥350 cells n≥2 coverslips, N=3 per condition). Statistical comparisons were performed using Kruskal–Wallis tests with Benjamini–Hochberg–corrected pairwise Wilcoxon tests (*p < 0.05, **p < 0.01). (**G**) Percentage of dead (PI-positive) cells at 180 min after addition of 10 mM LLOMe (>2000 cells per condition, mean ± SEM; n = 3; N = 3). Statistical significance was assessed by one-way ANOVA(**p < 0.01, ***p≤0.005).

To directly monitor Ca^2+^ signaling during membrane injury, we monitored cytosolic Ca^2+^ dynamics using a Cameleon reporter (38, 39) upon sequential LLOMe challenges. Because LLOMe acutely permeabilizes endolysosomes, luminal Ca^2+^ should leak to the cytosol and be detected by such a reporter. Control experiments confirmed that the Ca^2+^ chelator BAPTA-AM abolished the Cameleon signal, confirming that the responses arise from Ca^2+^ leakage or cytosolic influx. Likewise, digitonin induced a pronounced Ca^2+^ increase consistent with plasma membrane permeabilization, and subsequent EGTA rapidly quenched the signal, further validating that the observed changes reflect Ca^2+^ fluxes (Fig. S2A).

A first mild LLOMe pulse (4.5 mM) triggered a rapid, transient ΔR/R₀ increase in both HL5c and HL5c supplemented with Ca^2+^ (Fig. 2E and F). This brief spike coincided with short-lived Vps32 and Vps4 recruitment, which is consistent with efficient membrane repair, as previously observed using this dose of LLOMe (36, 40). A second, stronger pulse (45 mM) caused a sustained ΔR/R₀ elevation in HL5c, indicating a persistent Ca^2+^ leak indicative of slow or inefficient repair. In contrast, in Ca^2+^-supplemented cells the response was reversible and decayed toward baseline, indicative of efficient repair (Fig. 2E). Quantitatively, extracellular Ca^2+^ tended to increase the first-peak amplitude and markedly reduced the prolonged Ca^2+^ signal following the second pulse compared to HL5c (Fig. 2F).

Together, these data indicate that LLOMe-induced damage triggers cytosolic Ca^2+^ transients drawing on both extracellular Ca^2+^ influx and intracellular stores, potentially including ER via ER-endolysosome contact sites (41). Elevating extracellular Ca^2+^ enhances cytosolic Ca^2+^ release from endosomes upon damage, promotes sustained repair-competent signaling, accelerates re-acidification and limits cumulative endolysosomal injury. Consequently, the observed lysosomal resilience should translate into better cell survival. We have shown that excessive endolysosomal damage, either because the insult is too strong (high LLOMe) or because repair is compromised, as previously observed in *atg1*-KO or *trafE*-KO Dd cells (36), progresses to cumulative injury that the cell cannot resolve and ultimately results in death. We therefore tested whether the improved repair and re-acidification observed upon Ca^2+^ addition also enhance survival. Cell death was quantified by propidium iodide (PI), a membrane-impermeant DNA dye that enters cells upon loss of plasma membrane integrity. The fraction of PI-positive cells was significantly reduced in HL5c supplemented with Ca^2+^ compared with HL5c alone, whereas Ca^2+^ chelation with BAPTA-AM markedly increased PI uptake (Fig. 2G).

Together, these data indicate that extracellular Ca^2+^ promotes efficient repair and re-acidification of LLOMe-damaged endolysosomes, reduces cell death, and establishes extracellular Ca^2+^ as a key determinant of endolysosomal resilience and a practical entry point to identify the Ca^2+^-responsive sensors that mediate this protection.

### Ca^2+^ modulates the balance between ESCRT-mediated repair and autophagy

ESCRT components, autophagy, and the E3 ubiquitin ligase TrafE, which has been shown to coordinate ESCRT- and autophagy-mediated repair, play central roles in the repair of sterile endolysosomal damage as well as mycobacteria-induced MCV injury (36). Because extracellular Ca^2+^ supplementation reduced LLOMe-induced cell death, consistent with more efficient membrane repair, we reasoned that Ca^2+^ availability may influence how these repair pathways are engaged and coordinated. To test this, we used stable Dd lines expressing GFP-TrafE, GFP-Vps32, or GFP-Atg8a from a safe haven chromosomal locus (36, 40). Vps32 reports ESCRT-mediated membrane repair, Atg8a reports autophagy, and TrafE, coordinates these pathways (25). We then induced repairable endolysosomal injury with LLOMe and compared responses in HL5c versus HL5c + 5 mM Ca^2+^ (Fig. 3).

**Fig. 3.**
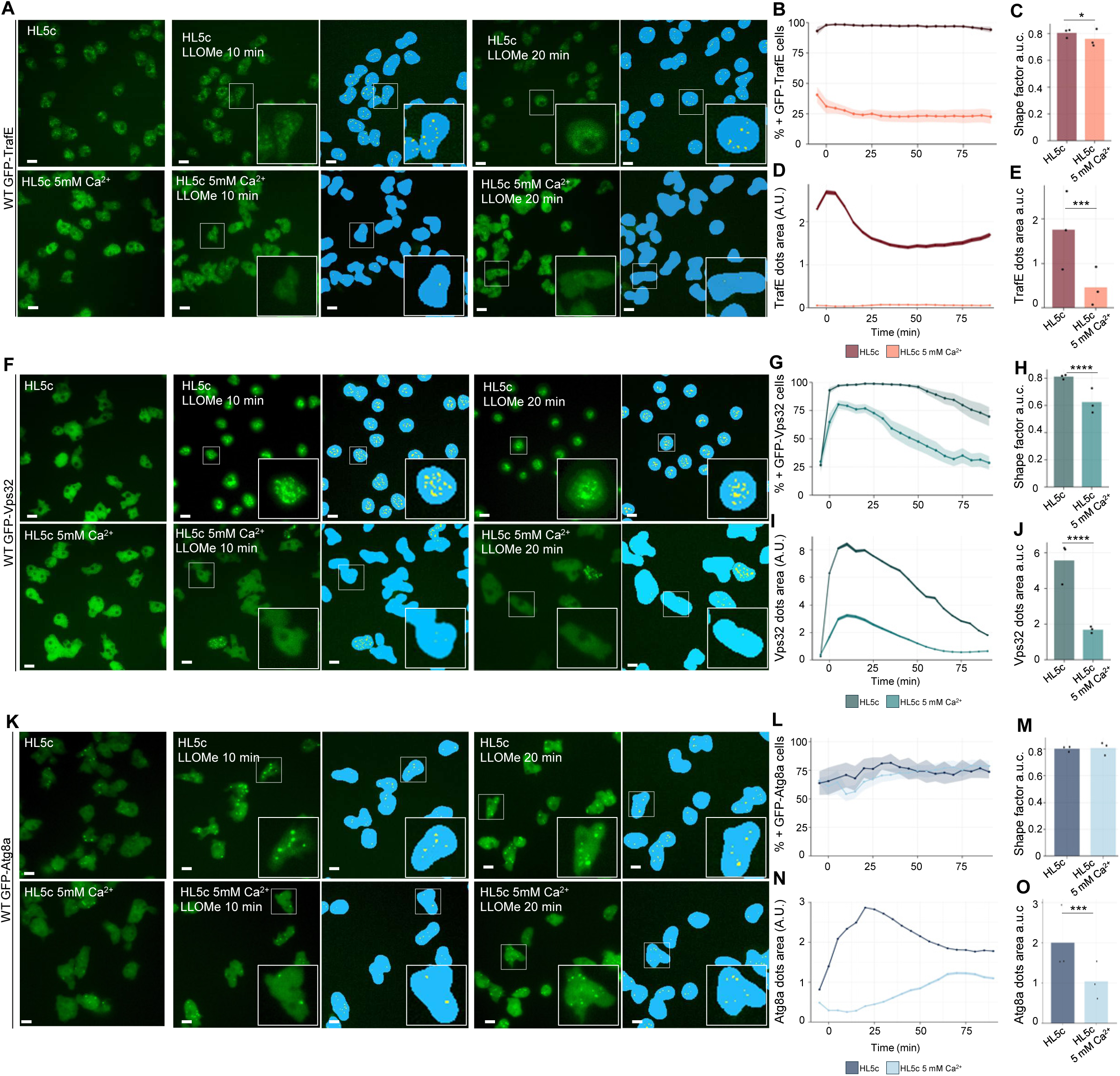
Extracellular Ca^2+^ favors cell repair. Dd WT expressing (**A-E**) GFP-TrafE, (**F-J**) GFP-Vps32 or (**K-O**) GFP-Atg8a were cultured in HL5c (top row) or HL5c + 5 mM Ca^2+^ (bottom row), and treated with 4.5 mM of LLOMe. (**A**, **F** and **K**) Representative live-cell images before treatment (0 min), and at 10 and 20 min after LLOMe addition. Insets shown 5× magnifications of selected cells. Cyan masks indicate the segmented cell area, and yellow masks indicate the segmented dot area used for quantification. Scale bar, 10 µm. Quantitative analyses for each marker show in (**B**, **G** and **L**) the percentage of dot-positive cells over time; in (**C**, **H** and **M**) the area under the curve (a.u.c.) of cell shape factor; in (**D**, **I** and **N**) the dots are over time; and in (**E**, **J** and **O**) the corresponding a.u.c. of dots area shown in (D, I and N)- (left bottom panel) and (right top panel) and dots area (right bottom panel). HL5c conditions are shown in darker colors, and HL5c + 5 mM Ca^2+^ in lighter tones. Analyses were performed on >300 cells for WT GFP-TrafE (HL5c), >630 cells for WT GFP-TrafE (HL5c + 5 mM Ca^2+^), >400 cells for WT GFP-Vps32 (HL5c), >500 cells for WT GFP-Vps32 (HL5c + 5 mM Ca^2+^), >500 cells for WT GFP-Atg8a (HL5c), and >400 cells for WT GFP-Atg8a (HL5c + 5 mM Ca^2+^) (n = 2–3; N ≥ 3). Statistical significance was assessed using the Kruskal–Wallis test, followed by pairwise Wilcoxon rank-sum tests with Benjamini–Hochberg correction for multiple comparisons (p < 0.05, **p ≤ 0.005, ***p < 0.0001). A.U, arbitrary unit.

LLOMe rapidly induced TrafE puncta in HL5c (Fig. 3A), resulting in a high and sustained fraction of GFP-TrafE-positive cells (Fig. 3B) accompanied by persistent cell rounding, reflected by an elevated shape factor (Fig. 3C). Because the shape factor a.u.c. integrates both the extent and duration of cell rounding, the higher a.u.c. observed in HL5c indicates prolonged damage-associated morphological stress. In contrast, Ca^2+^ supplementation strongly blunted TrafE recruitment, with few cells displaying detectable TrafE puncta (Fig. 3A and B). This attenuated and short-lived response was paralleled by a more rapid normalization of cell shape, yielding a significantly lower shape factor a.u.c. and indicating prompter recovery of cellular fitness (Fig. 3C). Analysis of TrafE puncta size further distinguished the two conditions. In HL5c, TrafE-positive dot area increased markedly and remained elevated for up to 20 min, consistent with sustained TrafE engagement at damaged membranes (Fig. 3D). Accordingly, the dot area a.u.c. was high, reflecting prolonged and extensive TrafE accumulation (Fig. 3E). By contrast, in Ca^2+^-supplemented cells, dot area formation was near-background and transient, resulting in a significantly reduced dot area a.u.c.. As ESCRT and autophagy act downstream of TrafE, this TrafE dynamics is expected to propagate to both repair pathways.

ESCRT-III (Vps32) recruitment mirrored TrafE (Fig. 3F to J). LLOMe induced robust Vps32 recruitment in HL5c cells, with a large fraction of GFP-Vps32-positive cells persisting over time (Fig. 3F and G) and sustained cell rounding reflected by an elevated shape-factor a.u.c. (Fig. 3H). Vps32-positive dot area increased markedly and remained elevated, yielding a high dot area a.u.c. consistent with prolonged ESCRT engagement (Fig. 3I and J). By contrast, Ca^2+^ supplementation reduced the fraction of Vps32-responding cells and was accompanied by a significantly lower shape factor than in HL5c cells. In addition, Vps32 puncta were less persistent in the presence of Ca^2+^, resulting in significantly lower dot area and shape factor a.u.c. These features are compatible with more efficient repair kinetics, in which ESCRT components are functionally recruited and resolved in a pre-programmed and efficient manner, thereby leaving a smaller temporal footprint.

LLOMe induced recruitment of Atg8a in both HL5c and Ca^2+^-supplemented cells, resulting in a comparable fraction of GFP–Atg8a–positive cells across conditions (Fig. 3K and L). Consistently, no significant differences in shape factor were observed between conditions, indicating similar overall cell morphology during autophagy engagement. However, the extent and timing of Atg8a accumulation differed markedly. In HL5c, Atg8a-positive dot area increased rapidly and reached an early maximum, yielding a higher dot area a.u.c. consistent with sustained autophagy engagement following damage (Fig. 3N and O). By contrast, Ca^2+^ supplementation reduced overall Atg8a accumulation and delayed the peak response, resulting in a significantly lower dot area a.u.c. This delayed and attenuated autophagy response suggests that efficient early repair by TrafE and ESCRT limits the requirement for autophagy, which is engaged primarily under conditions of unresolved membrane damage.

Notably, TrafE functions upstream of ESCRT and autophagy, and its limited accumulation in Ca^2+^-supplemented cells is consistent with a more transient recruitment that may fall below the temporal resolution of our imaging. In this scenario, TrafE dynamics would be sufficient to initiate downstream ESCRT and Atg8a responses while avoiding prolonged TrafE accumulation. This distinction is critical, as a smaller accumulation of repair factors can reflect two scenarios: (i) defective recruitment/repair, or (ii) more efficient, transient engagement that resolves damage without prolonged build-up. We discriminated between these alternatives using independent readouts of damage reversibility. Under Ca^2+^, cells rounded less and normalized shape more rapidly, LysoSensor fluorescence recovered faster, and PI uptake decreased, indicating preserved compartment function and improved survival (see Fig. 2). These outcomes are incompatible with impaired repair and instead support a model in which extracellular Ca^2+^ promotes rapid, pulse-like ESCRT engagement and limits the need for secondary intervention of autophagy, thereby shortening the repair phase and accelerating recovery. Together, these data indicate that elevating extracellular Ca^2+^ enhances the efficiency of ESCRT-dependent repair, yielding smaller, short-lived lesions and minimizing the requirement for extensive ESCRT and autophagy accumulation.

### PefA emerges as a Ca^2+^-responsive regulator of repair pathways

Our results indicate that extracellular Ca^2+^ does not simply amplify repair pathways but instead tunes their timing and coordination, promoting rapid and efficient resolution of membrane damage. However, the molecular basis by which Ca^2+^ exerts this control remains unclear. In particular, it is unknown which Ca^2+^-responsive factors sense damage upstream and coordinate TrafE, ESCRT, and autophagy responses to enforce this fast, pulse-like repair regime. Identifying such Ca^2+^-sensitive molecular sensors is therefore essential to understand how extracellular Ca^2+^ orchestrates repair pathway hierarchy and efficiency. To resolve the molecular basis of the Ca^2+^-sensitive phenotypes, we performed “GFP-trap” pulldowns of TrafE (the coordinator), Vps32 (ESCRT-III), and Atg8a (autophagy) in WT cells at steady state and after 10 minutes of LLOMe treatment, thereby identifying partners enriched or depleted upon damage and distinguishing shared from bait-specific interactors. Because exogenous Ca^2+^ enhances repair efficiency, triggering short-lived lesions, pulldowns were performed after LLOMe incubation in standard HL5c to maximize sensitivity to damage-induced interactomes.

Volcano plots of GFP–TrafE, GFP–Vps32, and GFP–Atg8a pull-downs under steady-state and LLOMe-treated conditions illustrate damage-dependent remodeling of each interactome (Fig. 4A). As expected, core ESCRT and autophagy components were selectively recovered with Vps32 and Atg8a, respectively, validating bait specificity and damage responsiveness. Pie charts (Fig. 4B) summarize the number of proteins captured with each bait before and after LLOMe-induced damage. In all three cases, damage increased the total number of interactors (enriched in red, depleted in blue and unchanged upon damage in grey). Proteins found in TrafE, Vps32 and Atg8a pulldowns were functionally classified with the Cytoscape StringApp (Fig. S3A, S3B and S3C, respectively) and displayed as stacked bar plots (Fig. 4C to 4E). Bars report the fraction of proteins in each functional category recovered with the indicated bait. Shades of red indicate the magnitude of enrichment after damage, blue denotes depleted proteins, and grey indicates those not detected (ND). Upon LLOMe treatment, the pulldowns revealed distinct pathway “fingerprints”. TrafE exhibited a pleiotropic interactome engaging membrane-trafficking and mitochondrial functions, consistent with its coordinating role, whereas nuclear and cell-division categories were prominently depleted, indicating disengagement from these functions upon membrane damage (Fig. 4C). Vps32 showed a more specialized profile, with strong enrichment of membrane-remodeling factors, notably V-ATPase subunits, consistent with ESCRT action at acidic endosomes (Fig. 4D). Like TrafE, Atg8a displayed a broad response with marked enrichment of the ubiquitin and proteasome categories and additional recruitment of signaling and cytoskeletal groups (Fig. 4E).

**Fig. 4.**
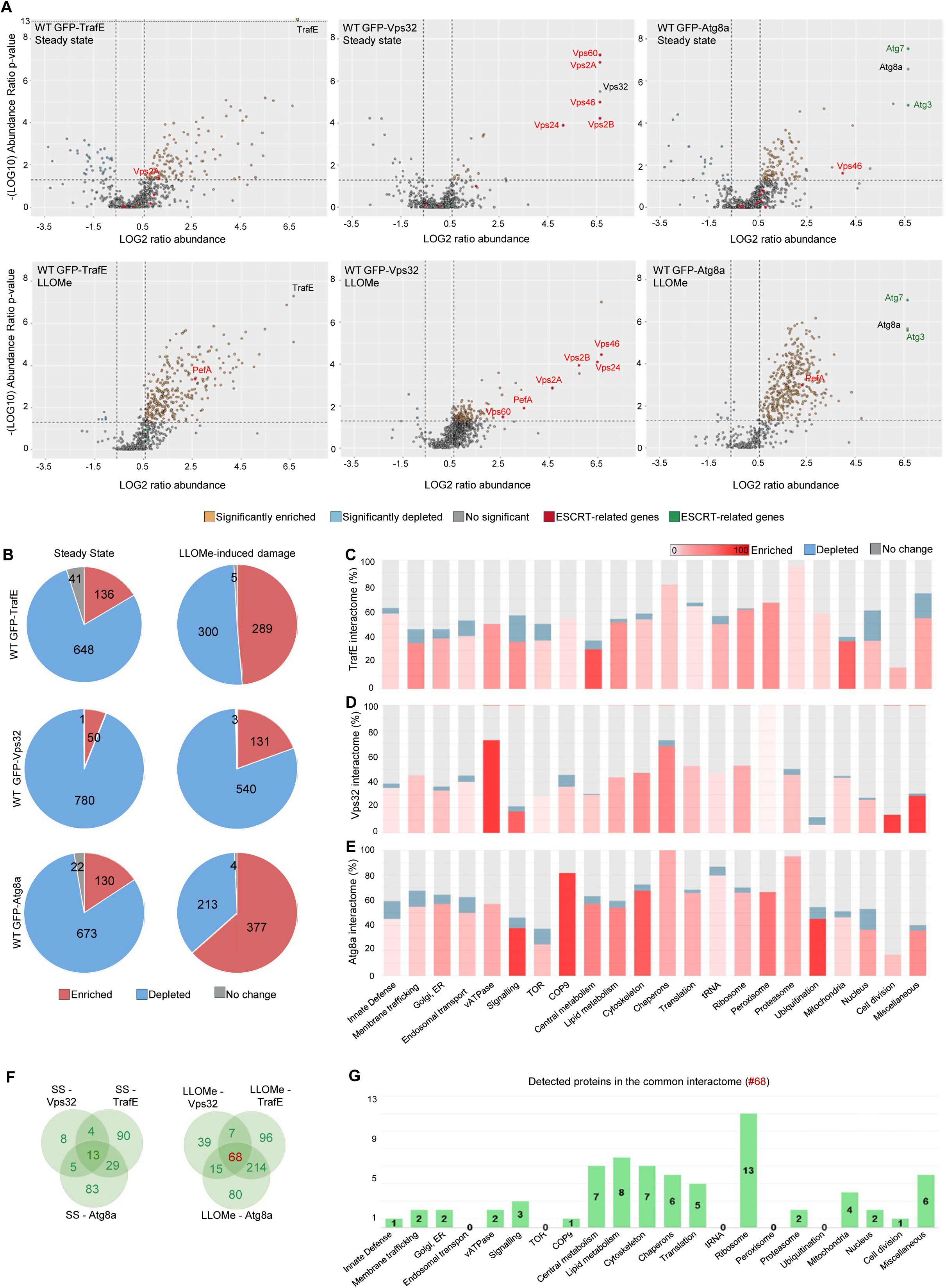
TrafE-, Vps32- and Atg8a–GFP interactomes at steady state and after lysosomal damage. (**A**) Pie charts representing the number of proteins detected in GFP-TrafE, GFP-Vps32 or GFP-Atg8a pulldown experiments, classified as significantly enriched (red), significantly depleted (blue) or not significantly changed (grey) relative to the corresponding GFP control under steady-state (SS) and LLOMe-treated conditions. Numbers within each slice indicate the number of proteins per category. (**B**-**D**) Bar plots showing, for each bait, the proportion of proteins that are enriched, depleted or non-detected within the total interactome, grouped by functional category based on Cytoscape/STRING assignation upon LLOMe treatment. (**E**) Venn diagrams illustrating the overlap between proteins significantly enriched in TrafE, Vps32 and Atg8a interactomes under steady-state conditions (left) and after LLOMe treatment (right). (**F**) Bar graph showing the distribution of common interactors between TrafE, Vps32 and Atg8a in response to LLOMe-induced damage. (**G**) Volcano plots of GFP pulldown datasets for TrafE, Vps32 and Atg8a under steady-state and LLOMe-treated conditions. Each dot represents a protein, plotted as log₂(fold change over GFP control) versus –log₁₀(p-value). Significantly enriched proteins are shown in yellow and depleted proteins in blue according to the applied thresholds; ESCRT-related proteins are highlighted in red and autophagy-related proteins in green.

We next intersected the significantly enriched interactomes using Venn diagrams (Fig. 4F) to show that the shared protein sets expand after LLOMe treatment, indicating that endolysosomal injury drives convergence of TrafE, ESCRT, and autophagy on common repair pathway proteins. The 68 proteins significantly enriched across all three baits were subsequently functionally classified using the Cytoscape StringApp (Fig. S4A) and summarized as stacked bar plots (Fig. 4G). Within this shared interactome, only a single protein mapped to the functional category of innate defense, which could be relevant during (mycobacterial) infection, whereas most differentially recovered proteins were associated with ribosomal complexes, metabolism, cytoskeleton, among others (Fig. 4G).

We therefore prioritized the sole innate-defense protein significantly enriched across all three baits (TrafE, Vps32, Atg8a), the penta-EF-hand Ca^2+^-binding protein PefA, an ALG-2–like protein. PefA was selectively enriched after LLOMe compared to steady state (Fig. S4B), consistent with damage-dependent recruitment. In the light of its known biochemistry, (14, 42), , this enrichment pattern positions PefA as a potential molecular bridge between Ca^2+^ signaling and the coordinated activities of TrafE, ESCRT, and autophagy during membrane repair.

### PefA–dependent Ca^2+^ signaling shapes ESCRT and autophagy responses to lysosomal damage

PEF proteins are classical Ca^2+^ switches. Ca^2+^ loading at specific EF-hands converts ALG-2/PefA into a “target-on” state that enables both partner engagement and membrane association (14). In animals, ALG-2 binds the ESCRT adaptor Alix in a strictly Ca^2+^-dependent manner via EF-1/EF-3, dimerizes through EF-5, and can associate with membranes indirectly through Alix/TSG101 as well as directly with acidic lipids. Moreover, chelating Ca^2+^ abolishes these interactions (43, 44) . Dd encodes two ALG-2-like paralogs with distinct Ca^2+^ sensitivities (PefA ∼30 µM and PefB ∼450 µM), and PefA homodimerization and Alix binding are Ca^2+^-dependent (45). *In vivo*, PefA-GFP rapidly accumulates at wound sites only when extracellular Ca^2+^ is available, while EGTA blocks its recruitment, consistent with Ca^2+^-licensed membrane binding (46). Together, these properties provide a mechanistic rationale for the selective recruitment of PefA following LLOMe-induced Ca^2+^ leakage at damaged compartments, where it promotes transient membrane association. This shift would nucleate Alix-positive ESCRT assembly, thereby linking Ca^2+^ signals to coordinated TrafE, ESCRT, and autophagy engagement at injury sites (31, 36, 47).Having identified PefA as the only innate-defense factor selectively enriched after LLOMe, we next examined whether PefA couples Ca^2+^ signals to coordinate TrafE, ESCRT and autophagy activity at damaged endolysosomes. We visualized and quantified LLOMe-triggered dynamics of TrafE, Vps32 and Atg8a in Dd Δ*pefA* cells and, in parallel, in WT cells treated with BAPTA-AM to chelate intracellular Ca^2+^ (Fig. 5). For completeness, an extended comparison including WT cells cultured with extracellular Ca^2+^ supplementation, Δ*pefA* cells, and BAPTA-treated WT cells is provided in Fig. S5. Representative fields are shown for TrafE, Vps32 and Atg8a under basal conditions and 10–20 min after LLOMe addition, white boxes mark regions enlarged as insets. Quantifications to the right summarize the percentage of responsive cells that exhibit a detectable punctum, and the response magnitude measured as total GFP-reporter dot area per cell.

**Fig. 5.**
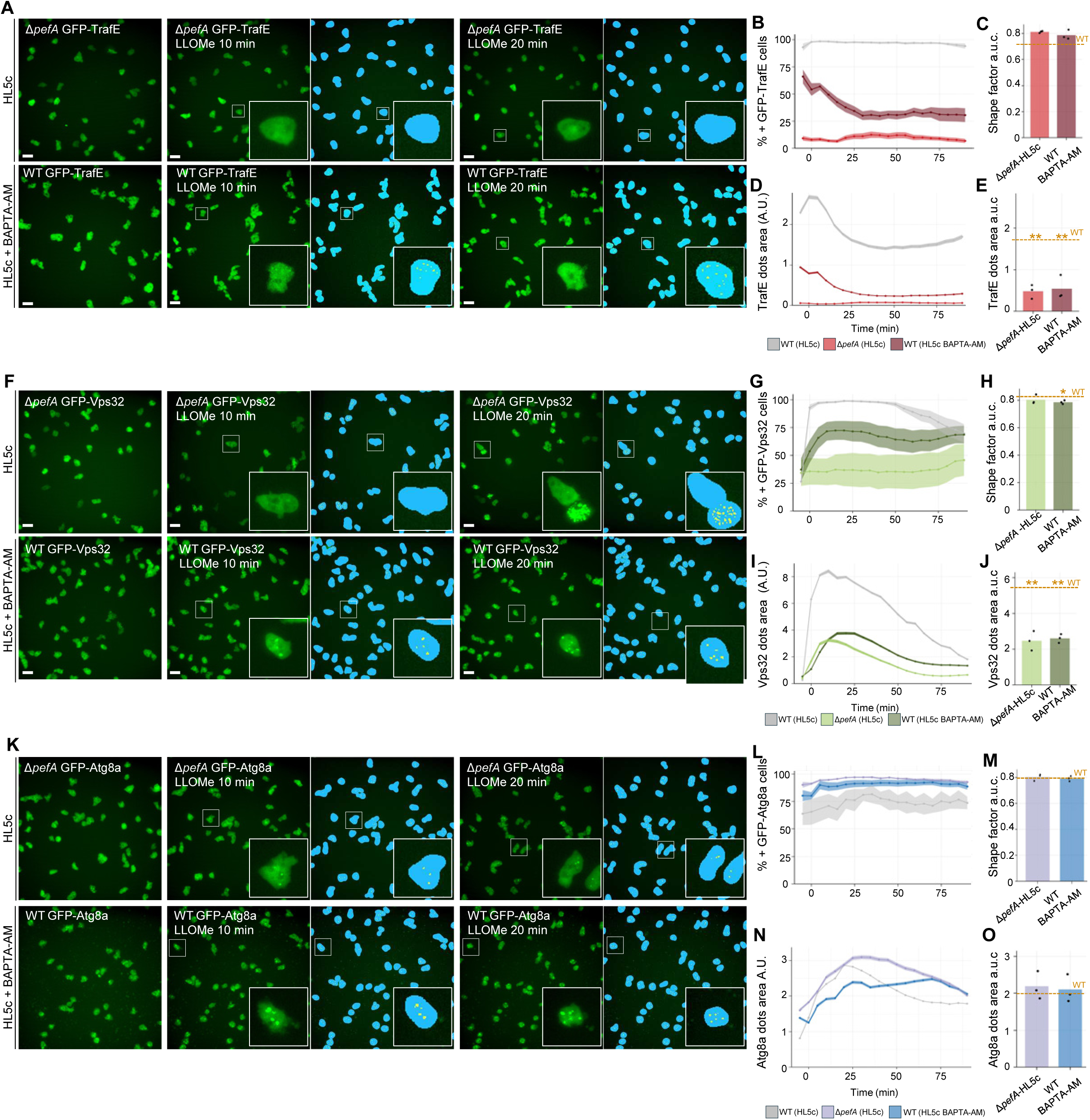
Cell repair is dampened in absence of calcium signaling. Dd WT cultured in HL5c + BAPTA or Δ*pefA* Dd cells cultured in HL5c (top row) expressing GFP-TrafE (**A-E**), GFP-Vps32 (**F-J**), or GFP-Atg8a (**K-O**) were treated with 4.5 mM of LLOMe. (**A**, **F** and **K**) Representative live-cell images before treatment (0 min), and at 10 and 20 min after LLOMe addition. Insets are 5× magnifications of selected cells. Cyan masks indicate the segmented cell area and yellow masks indicate the segmented dot areas used for quantification. Scale bar, 10 µm. Quantitative analyses for each marker include (**B**, **G** and **L**) the percentage of dot-positive cells over time; (**C**, **H** and **M**) the area under the curve (a.u.c.) of cell shape factor; (**D**, **I** and **N**) dots area dynamics over time; and (**E**, **J** and **O**) the corresponding a.u.c. of dots area. WT cells cultured in BAPTA are shown in darker colors, and Δ*pefA* in lighter tones. WT cells cultured in HL5c (from Fig 3) are shown in grey as a reference. Analyses were performed on >17,000 cells per WT GFP-TrafE (HL5c + BAPTA), >7,000 cells per Δ*pefA* GFP-TrafE (HL5c), >10,000 cells per WT GFP-Vps32 (HL5c + BAPTA), >7,000 cells per Δ*pefA* GFP-Vps32 (HL5c), >12,000 cells per WT GFP-Atg8a (HL5c + BAPTA), and >10,000 cells per Δ*pefA* GFP-Atg8a (HL5c), per conditions and replicates (n = 2–3; N ≥ 3). Statistical significance was assessed using the Kruskal–Wallis test, followed by pairwise Wilcoxon rank-sum tests with Benjamini–Hochberg correction for multiple comparisons (# p < 0.05, ## p ≤ 0.005, difference to WT (HL5c) condition).

Loss of PefA markedly impaired TrafE recruitment in response to LLOMe-induced damage (Fig. 5A to E). In Δ*pefA* cells, both representative images (Fig. 5A) and quantification (Fig. 5B) showed a significantly reduced fraction of GFP-TrafE–positive responding cells. Chelation of intracellular Ca^2+^ with BAPTA-AM produced an intermediate phenotype (Fig. 5A and B). Slightly higher shape factor measurements were observed in Δ*pefA*, and BAPTA-treated cells compared to WT cells (Fig. 5C). In addition to fewer responding cells, the amplitude of the TrafE response was reduced. The TrafE-positive dot area was significantly smaller in Δ*pefA* cells and moderately reduced in BAPTA-treated cells (Fig. 5D), resulting in a significantly lower dot area a.u.c. compared to WT (Fig. 5E). These data indicate defective assembly or expansion of TrafE-associated repair structures even when recruitment occurred. This difference is consistent with kinetic constraints, as BAPTA-AM must be de-esterified and diffuse before it can buffer Ca^2+^, whereas the five EF-hands of PefA can respond rapidly to local Ca^2+^ increases within pre-assembled complexes at damage sites, enabling efficient early complex assembly. Thus, in WT cells, some PefA-dependent events may still occur but are attenuated under BAPTA treatment.

ESCRT-III recruitment displayed a similar dependency on PefA and Ca^2+^ signaling (Fig. 5F to J). Representative images illustrate the reduced and less persistent Vps32 puncta formation in the absence of PefA (Fig. 5F). In Δ*pefA* cells, the fraction of GFP–Vps32-positive responders was significantly reduced compared with WT and the BAPTA-treated WT cells again showed an intermediate phenotype (Fig. 5G). Shape factor measurements were comparable across WT, Δ*pefA*, and BAPTA-treated cells (Fig. 5H). Analysis of Vps32 puncta size revealed that Δ*pefA* cells exhibited significantly reduced dot area and dot area a.u.c. compared with WT (Fig. 5I and J), indicating impaired ESCRT assembly/expansion at damaged membranes. Together, these results indicate that PefA and Ca^2+^ promote efficient Ca^2+^-dependent engagement and expansion of ESCRT repair complexes.

Atg8a behaved differently from TrafE and Vps32 (Fig. 5K to O). Both Δ*pefA* and BAPTA-treated cells displayed a higher fraction of GFP-Atg8a positive responders than WT across the time course in steady state and upon LLOMe damage (Fig. 5L), consistent with more frequent autophagy engagement when PefA function or Ca^2+^ signaling are perturbed. The shape factors remained comparable across conditions (Fig. 5M). Despite the higher frequency of Atg8a-positive puncta, Atg8a dot area dynamics differed in WT versus Δ*pefA* and BAPTA-treated cells (Fig. 5N). In WT cells, LLOMe treatment triggered a pronounced increase in Atg8a dot area, whereas in Δ*pefA* and BAPTA-treated cells, dot area increased less prominently (Fig. 5N). Accordingly, the dot area a.u.c. did not differ significantly between conditions (Fig. 5O), indicating that the overall cumulative Atg8a area over the experiment is similar, but in the absence of PefA is redistributed in time and across more numerous, smaller structures.

Together, these data demonstrate that PefA acts as a Ca^2+^-responsive regulator that promotes early assembly and expansion of TrafE- and ESCRT-dependent repair complexes without altering overall cell morphology. In the absence of PefA or upon buffering of cytosolic Ca^2+^ with BAPTA, the primary membrane repair is compromised, resulting in altered autophagy. Under these conditions, Atg8a recruitment is sustained over time and does not resolve as observed in wild-type cells, indicating prolonged autophagy engagement. However, this sustained signal is accompanied by a reduced peak amplitude compared to wild type, suggesting that although autophagy is more persistently engaged in the absence of efficient ESCRT-mediated repair, its activity is less efficient. Thus, impaired Ca^2+^–PefA signaling leads to increased reliance on autophagy coupled to delayed or ineffective turnover, rather than productive completion of the pathway. This establishes PefA as a key molecular link between Ca^2+^ signaling and the coordinated hierarchy of membrane repair pathways.

### Lysosomal damage remodels the PefA interactome toward a core repair network

While previous results identified PefA as a key Ca^2+^-responsive regulator of membrane repair pathway hierarchy, the molecular mechanisms by which PefA exerts this function remain unknown. In particular, it is unclear how PefA interfaces with TrafE, ESCRT, and autophagy components, and whether these interactions are preassembled at steady state or dynamically remodeled upon membrane damage. We therefore reasoned that defining the PefA interaction landscape both under basal conditions and upon endolysosomal damage is critical to understand how Ca^2+^ signals are translated into the coordinated and timely recruitment of distinct repair machineries. To this end, we analyzed the PefA interactome under steady-state conditions and following LLOMe-induced damage by affinity purification of GFP–PefA coupled to mass spectrometry (AP–MS) (Fig. 6). For comparative and integrative analysis, the PefA interactome was processed using the same STRING/Cytoscape workflow and functional module classification previously applied to the TrafE, ESCRT, and autophagy interactomes, thereby enabling direct comparison across repair pathways using a unified analytical framework.

**Fig. 6.**
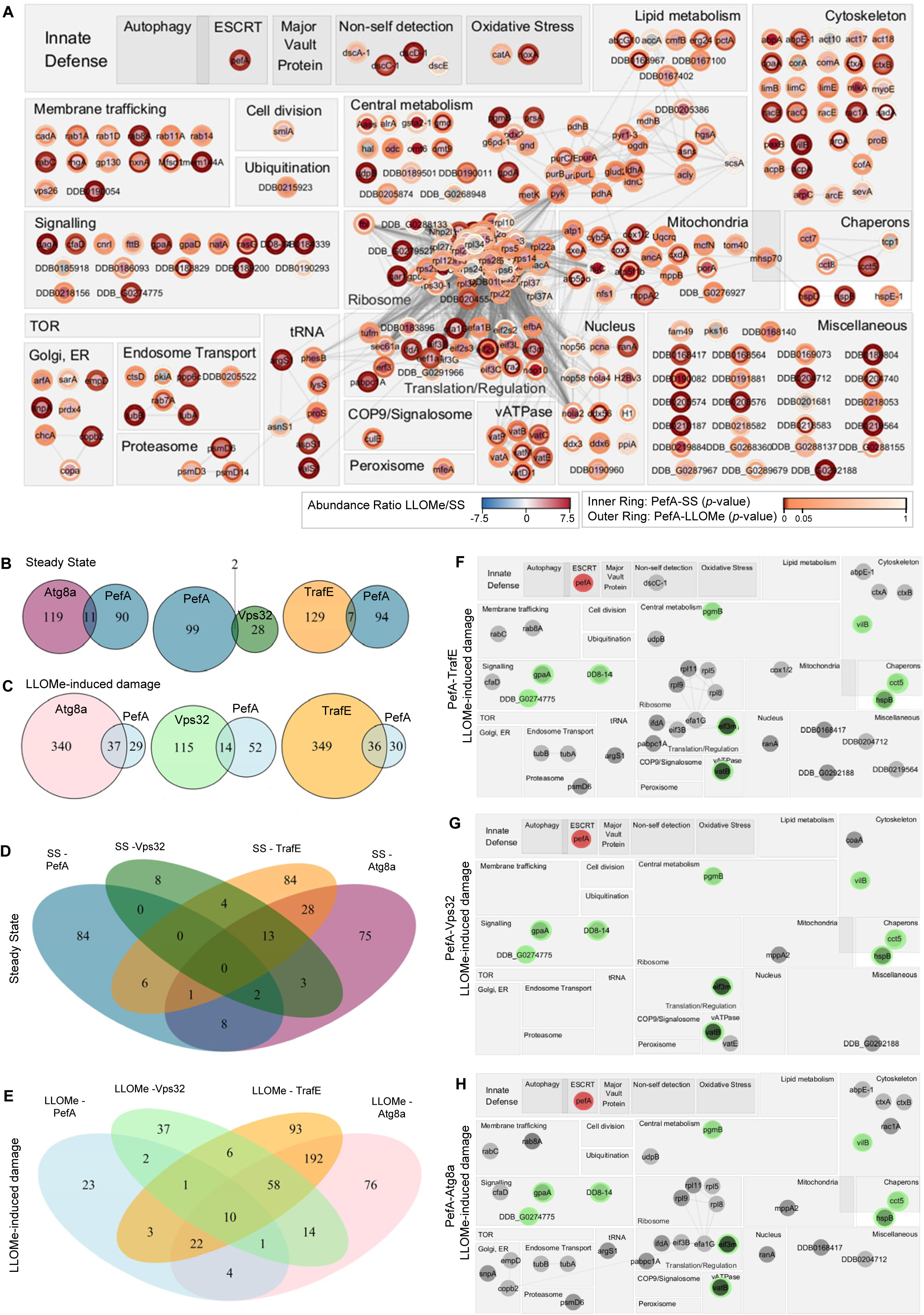
PefA functional organization and overlap with Atg8a, Vps32 and TrafE interactomes at steady state and after lysosomal damage. (**A**) GFP-PefA pulldowns proteins were imported into STRING and visualized in Cytoscape. Each node represents a protein, and proteins are grouped into manually curated functional categories. Node shading indicates the relative change in abundance (LLOMe/steady state), while inner and outer rings denote statistical significance at steady state and after lysosomal damage, respectively. (**B** and **C**) Pairwise comparison of significantly enriched proteins for each bait (PefA, TrafE, Vps32 and Atg8a) at steady state (**B**) and after LLOMe treatment (**C**). Venn diagrams illustrate the size of each interactome, and the number of proteins shared with PefA. (**D** and **E**) Four-way comparison of the Atg8a, Vps32, TrafE and PefA interactomes at steady state (**D**) and after LLOMe treatment(**E**). Venn diagrams depict the global overlap between the four baits and highlight proteins that are uniquely or commonly recruited under each condition. (**F-H**) Functional distribution of proteins shared between PefA and each of the other baits following LLOMe-induced damage. The functional categories defined in (A) are used to classify shared interactors for PefA–TrafE (**F**), PefA–Vps32 (**G**) and PefA–Atg8a (**H**). PefA interactors are shown in red, proteins common to all four baits in green, and the remaining interactome in grey.

The interaction network (Fig. 6A) uses node shading intensity to represent changes in protein abundance upon LLOMe treatment relative to steady state, while the inner and outer rings report statistical significance at steady state and after damage, respectively. Endolysosomal damage selectively strengthened several modules in which the outer ring exceeded the inner ring. The most prominent were ESCRT, autophagy, membrane trafficking (including multiple Rab GTPases), the v-ATPase, cytoskeleton (actin/myosin components and regulators), lipid metabolism, and chaperones. In these categories, many nodes were darkly shaded, indicating damage-associated changes in abundance in addition to increased statistical enrichment. The v-ATPase box, in particular, contained multiple subunits with near-maximal shading and especially prominent outer rings. Similarly, trafficking and cytoskeletal nodes generally showed brighter outer than inner rings, suggesting that, upon damage, PefA strengthens interactions with membrane-remodeling machineries and the actin scaffold. Lipid metabolism enzymes also exhibited intense node shading with bright outer rings probably associated to an elevated lipid synthesis/turnover during repair. By contrast, ubiquitination and the proteasome contained fewer strongly enriched nodes. High baseline (inner-ring) significance with minimal LLOMe-induced change was seen for ribosome, translation, central metabolism, nucleus, tRNA synthetases, and the COP9/signalosome, indicating constitutive/housekeeping associations with PefA. Together, the most intense functional groups potentiated after LLOMe support a model in which PefA acts as a damage-responsive hub linking lysosomal acidification, autophagy/ESCRT activation, and membrane remodeling.

We next compared the PefA interactome with the previously characterized interactomes of TrafE, Vps32, and Atg8a. Across all comparisons, overlap with PefA was minimal under steady-state conditions but increased markedly following LLOMe-induced damage, as revealed by both pairwise analyses and higher-order Venn diagrams (Fig. 6B–E). Complete lists of shared interactors under steady-state and damage conditions, as well as detailed pairwise and higher-order overlap analyses, are provided in Tables S1–S4. Notably, no protein was shared across all four interactomes at steady state, whereas LLOMe treatment uncovered a common, damage-induced core of ten proteins (highlighted in green in Fig. 6F–H). This core comprised factors involved in signaling (GpaA, DDB_G0274775/DDB-14), central metabolism (PgmB), translation/regulation (eIF3m), cytoskeleton organization (VilB), chaperone function (Cct5, Hsp8), and v-ATPase–dependent acidification (VatB), indicating convergence of PefA, TrafE, ESCRT, and autophagy on a shared functional module supporting lysosomal integrity and membrane repair.

Despite this shared core, the composition of the overlapping interactomes revealed pathway-specific biases. The PefA–TrafE overlap showed the broadest intersection, enriched for proteostasis, translation, cytoskeleton, central metabolism, and v-ATPase components, and uniquely included the non-self-detection lectin DscC, pointing to a TrafE-linked branch of the PefA network integrating damage and non-self-sensing (Fig. 6F). In contrast, overlap with ESCRT-III was shifted toward lysosome remodeling and acidification, including an additional v-ATPase subunit (VatE) and lacking ribosomal components (Fig. 6G). The PefA–Atg8a overlap largely mirrored the TrafE-associated profile but without the enhanced acidification signature seen with ESCRT-III, consistent with a preferential link to autophagy and proteostasis rather than organelle remodeling (Fig. 6H).

### PefA links Ca^2+^ sensing to Alix–ESCRT activation

In animal cells, acute plasma membrane injury triggers a rapid Ca^2+^ influx that recruits the Ca^2+^ sensor ALG-2, which in turn engages Alix and ESCRT-III/Vps4 to execute repair of laser-induced plasma membrane wounds in a Ca²-dependent manner (42). Endolysosomal lesions behave similarly. LLOMe-induced damage elicits robust ESCRT responses in which Alix and ESCRT-III subunits accumulate at injured lysosomes to promote repair (16).

Notably, despite the strong parallels with mammalian systems, Alix was not detected in the PefA interactome under either steady-state or LLOMe-induced damage conditions questioning whether PefA physically associates with Alix, or whether their interaction is highly transient and therefore not captured by affinity purification. Given that Ca^2+^-dependent ALG-2–ALIX recruitment during membrane repair is known to occur on rapid timescales in animal cells (42), we reasoned that a similar fast and dynamic interaction might operate in Dd. We thus wondered whether Alix recruitment and functions are nonetheless acutely regulated by Ca^2+^ and PefA during membrane damage. We therefore performed live-cell imaging approaches to directly assess the dynamics of Alix recruitment during plasma membrane and endolysosomal damage (Fig. 7).

**Fig. 7.**
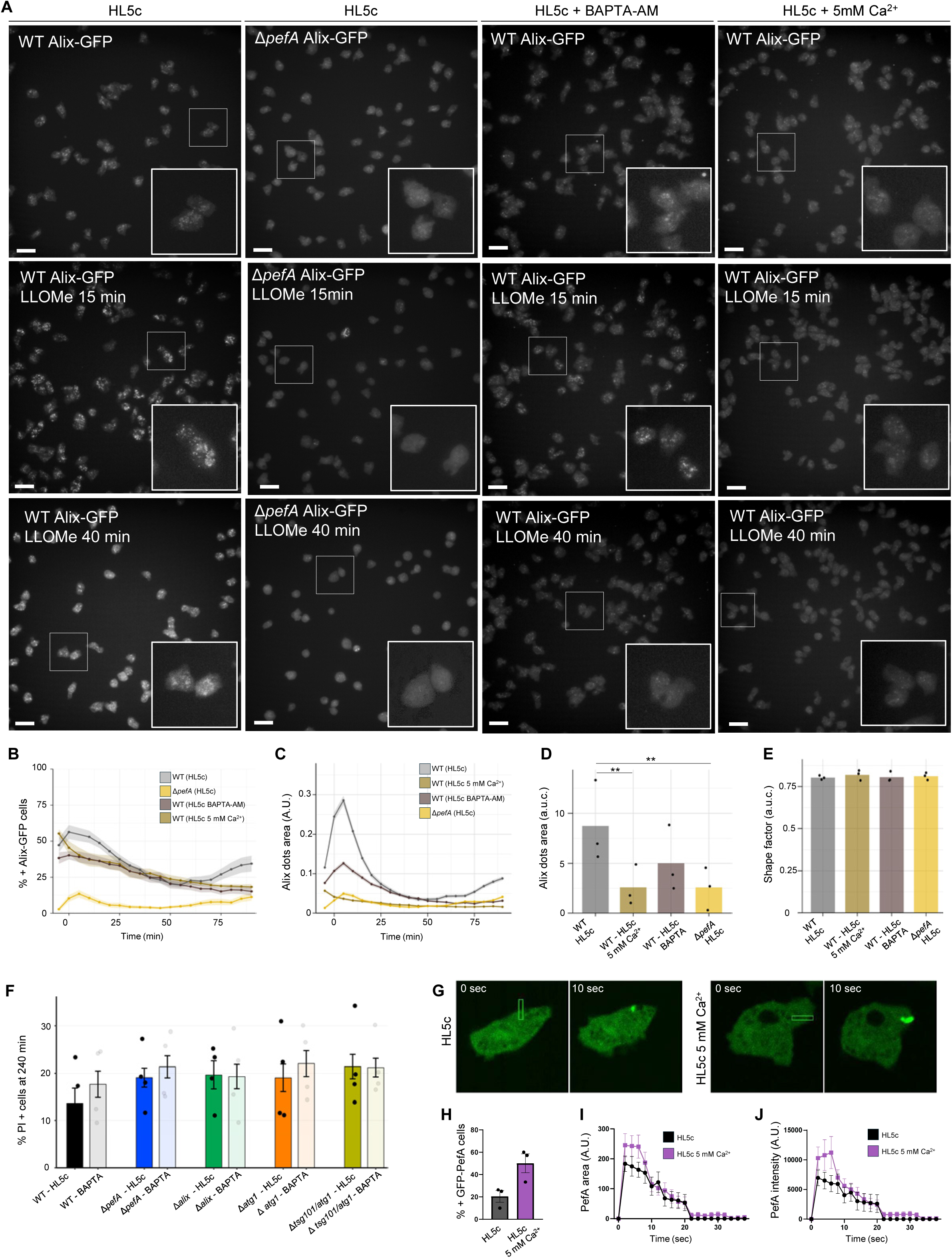
Ca^2+^ availability and PefA regulate membrane repair efficiency and cell survival following lysosomal damage. Dd WT cultured in HL5c (grey), HL5c + 5 mM Ca^2+^ (light brown) or HL5c + BAPTA (brown); Δ*pefA* Dd cells cultured in HL5c, were treated with 4.5 mM of LLOMe. (**A**) Representative live-cell images before treatment (0 min) and at 15 and 40 min after LLOMe addition. Insets are 5× magnifications of selected cells. Scale bar, 10 µm. (**B**) Percentage of dot-positive cells over time. (**C**) Area under the curve (a.u.c.) of cell shape factor. (**D**) Dynamics of dots area over time. (**E**) Corresponding a.u.c. of dots area. (F) Percentage of dead (PI-positive) cells measured 180 min after LLOMe addition in WT, Δ*pefA*, Δ*alix*, Δ*atg1* and Δ*tsg101*/Δ*atg1* cells cultured in HL5c or HL5c + BAPTA (>7,000 cells per WT (HL5c), >13,000 cells per WT (HL5c + 5 mM Ca^2+^), >13,000 cells per WT (HL5c + BAPTA), >13,000 cells per Δ*pefA* (HL5c); mean ± SEM; n = 3; N = 3). Statistical significance was assessed using one-way ANOVA (**p < 0.01, ***p ≤ 0.005). (**G**) Representative two-photon FRAP images of Dd cells expressing GFP-PefA cultured in standard HL5c medium or HL5c supplemented with 5 mM Ca^2+^ following laser-induced plasma membrane damage. (**H**) Quantification of GFP-PefA recruitment at wound sites (**I - J**) Quantitative analyses of GFP-PefA recruitment area (I) and fluorescence intensity (J), mean ±SEM, n=38 (HL5c + 5 mM Ca^2+^) and n=59 (HL5c), N=3.

Representative images show Alix–GFP localization in WT cells, Δ*pefA* cells, WT cells treated with BAPTA-AM, and WT cells supplemented with extracellular Ca^2+^ under basal conditions and following LLOMe treatment (Fig. 7A). Quantification revealed that the fraction of Alix-GFP positive responding cells was higher in WT cells, significantly reduced in Δ*pefA* cells, and intermediate in BAPTA-treated and Ca^2+^-supplemented conditions (Fig. 7B). These differences were mirrored by changes in response magnitude, as both Alix–GFP dot area (Fig. 7C) and dot area a.u.c. (Fig. 7D) were reduced in Δ*pefA* cells or Ca^2+^ supplementation, and partially decreased by Ca^2+^ chelation. Shape factor measurements were comparable across all conditions (Fig. 7E). Together, these findings recapitulate the Vps32 phenotypes and align with the established position of Alix upstream of ESCRT-III. The strong reduction of Alix recruitment in Δ*pefA* cells indicates that PefA acts upstream of Alix to enable efficient ESCRT activation. The diminished phenotype observed upon Ca^2+^ supplementation in comparison to WT further supports a kinetic model in which Ca^2+^–PefA signaling accelerates Alix engagement and resolution, rendering the response transient and partially undetectable at our temporal resolution.

As an integrative readout, we quantified cell death following LLOMe-induced damage across ESCRT and autophagy mutants (Fig. 7F). Calcium chelation with BAPTA in WT cells, as well as loss of PefA, Alix, or Atg1, increased cell death relative to WT cells, while combined loss of ESCRT (*tsg101*) and autophagy (*atg1*) resulted in the highest levels of cell death. Notably, the BAPTA-induced increase in cell death observed in WT cells was not further enhanced, or was only weakly enhanced, in the mutant backgrounds. Although differences among mutant backgrounds primarily manifested as trends rather than statistically significant separations, a schematic model summarizing the genetic contributions to cell death is provided (Fig. S6C-F).

In mammalian cells, plasma membrane injury activates a Ca^2+^-dependent ALG-2–ALIX–ESCRT repair pathway (42). We therefore examined whether Ca^2+^ and PefA regulate ESCRT activation at damaged plasma membranes in Dd. Using laser-induced membrane wounding combined with live imaging, we quantified rapid PefA recruitment. In standard medium, PefA accumulated at wound sites within seconds, and this recruitment was further enhanced by extracellular Ca^2+^ supplementation (Fig. 7G and H). Quantitative analyses showed enhanced PefA recruitment upon Ca^2+^ supplementation with higher peak area and intensity (Fig. 7I and J). Complementary laser-wounding experiments using cytosolic GFP, ALIX–GFP, GFP–Vps32, and GFP–Atg8a showed that ALIX and Vps32 were modestly recruited to wound sites only in the presence of Ca^2+^, with ALIX accumulating more frequently than Vps32 (Fig. S6G and S6H).

Together, these data indicate that extracellular Ca^2+^ enhances PefA recruitment to damaged plasma membranes, consistent with a role for PefA as a Ca^2+^-responsive regulator that promotes rapid, ESCRT-mediated membrane repair.

### The Ca^2+^–PefA axis controls MCV integrity during mycobacterial infection

Having established that extracellular Ca^2+^ and PefA are critical determinants of ESCRT and autophagy engagement and functionality at acutely damaged membranes, we next asked whether this Ca^2+^/PefA interplay extends to the pathogen vacuole during infection (Fig. 8).

**Fig. 8.**
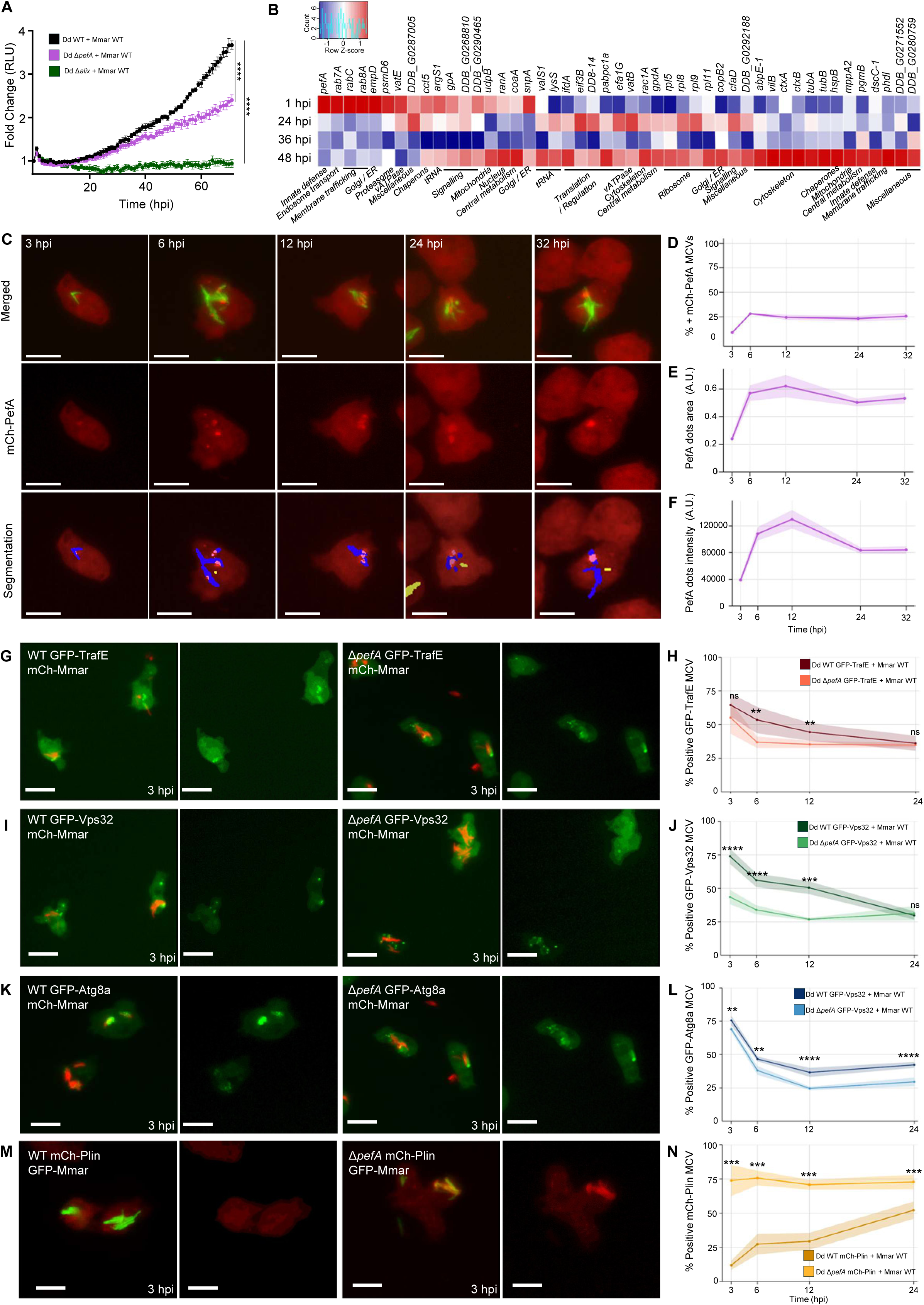
PefA controls mycobacterial replication and MCV-associated host responses in Dd. (**A**) Dd WT, Δ*pefA* and Δ*alix* cells were infected with bioluminescent Mm WT. Bioluminescence was measured for 72 hours (mean fold change ± SEM, n=3, N = 3, two-way ANOVA–Fisher’s LSD test, ****p≤ 0.0001). RLU, relative luminescence units. (**B**) RNA-seq analysis of infected WT cells. Heatmap showing genes significantly enriched in Fig. 6A at 1, 24, 36 and 48 hours post-infection (hpi). (**C**) Representative live-cell high-content (HC) microscopy images of Dd Δ*pefA* expressing mCherry-PefA infected with Mm WT expressing GFP. Scale bars, 10 μm. (**D-F**) Quantification of the proportion of PefA-positive MCVs (**D**), MCV-associated dot area (**E**), and dot intensity (**F**) at 3, 6, 12, 24 and 32 hpi (mean ± SEM, N = 3, n ≥ 650 MCVs per experiment). Dd WT and Δ*pefA* cells expressing GFP-TrafE, GFP-Vps32, GFP-Atg8a or mCherry-Plin were infected with GFP- or mCherry-expressing Mmar and imaged live by HC microscopy at 3, 6, 12 and 24 hpi. (**G, I, K, M**) Representative images of Dd WT and Δ*pefA* cells expressing the indicated markers at 3 hpi. Scale bars, 10 μm. (**H**, **J**, **L**, **N**) Quantification of infected cells positive for the indicated reporters (mean ± SEM, N = 3, n ≥ 750 MCVs per experiment and marker) Statistical comparisons between WT and Δ*pefA* conditions were performed independently at each time point using a two-sided Wilcoxon rank-sum test (**p ≤0.01, ***p ≤0.005, ****p ≤0.0001).

First, Dd cells were infected with lux-expressing Mmar WT and the intracellular growth measured by the luminescence readout. The bacterial burden (quantified in RLU) increased steadily in WT cells, reaching 4-fold by 72 hpi. Growth was significantly reduced in PefA-deficient cells (2–2.5-fold), indicating that loss of PefA might result in premature cytosolic escape and restricted bacteria growth. A similar pattern has been previously observed in *tsg101*-KO cells, where defective MCV repair results in premature escape to the cytosol and partial restriction by xenophagy (29). Strikingly, absence of Alix nearly abolished Mmar growth, demonstrating that autophagy, and ESCRT engagement via Alix are required to maintain the replicative MCV niche and protect against radical xenophagy.

Then, to determine whether the damage-repair network identified by LLOMe treatment is also engaged during mycobacterial infection, we performed RNA-seq analysis of Dd infected with Mmar WT and focused on the 68 host genes corresponding to the common interactome of TrafE, Vps32, and Atg8a identified upon LLOMe-induced damage (Fig. 4E). Remarkably, the vast majority of these genes were significantly modulated during infection (Fig. 8B), indicating that the shared interactome also represents a transcriptionally responsive module during MCV stress. Moreover, these data suggest that the repair pathways uncovered using sterile LLOMe damage are also transcriptionally modulated during pathogen-induced vacuolar injury.

Hierarchical clustering revealed two major temporal expression blocks. An early-response cluster (1 hpi) was enriched for genes involved in innate defense, membrane trafficking, signaling, ER/Golgi function and proteostasis, consistent with rapid sensing and stabilization of damaged compartments. Notably, *pefA* was strongly upregulated at this early time point, supporting its role as an early Ca^2+^-responsive regulator during infection. Differential expression of additional ESCRT, autophagy, and TRAF-family genes during infection is shown in Fig. S7A–C. In contrast, a late-response cluster (24–48 hpi) comprised genes associated with cytoskeletal remodeling, ribosome and translation, mitochondrial function, central metabolism, and membrane trafficking, reflecting sustained remodeling and adaptation to prolonged intracellular infection. Together, these results establish the common TrafE–Vps32–Atg8a interactome as a core host response module required for maintaining vacuolar integrity and demonstrate that LLOMe-induced damage faithfully models key aspects of mycobacteria-triggered MCV stress.

The transcriptional upregulation of *pefA* early during Mmar infection, together with its central position in the shared damage-repair interactome, prompted us to directly examine PefA localization during infection. To this end, we engineered Dd cells to express mCherry–PefA and monitored its dynamics during intracellular GFP-Mmar infection. Representative confocal images show that mCherry–PefA associates with MCV at successive time points (Fig. 8C). Quantitative analyses revealed that the fraction of mCherry-PefA-positive MCVs increased at early time points and then remained constant from 12 to 32 hpi (Fig. 8D). In contrast, the magnitude of PefA recruitment continued to rise, as reflected by increased fluorescence intensity at the MCV and expansion of mCherry-PefA puncta area, which peaked around 12 hpi before declining (Fig. 8E and F). These dynamics indicate progressive accumulation of PefA at MCVs rather than an increase in the number of PefA-positive structures. Together, these results demonstrate that PefA is persistently recruited to the MCV during infection, consistent with a role in sensing and responding to pathogen-induced MCV damage.

To close the loop between sterile lysosomal damage and pathogen-induced vacuolar injury, we tested whether PefA is required to recruit and sustain host repair machineries at the MCV during Mmar infection. Building on the sterile-damage data, we monitored the dynamics of GFP-tagged TrafE, Vps32, and Atg8a in WT and Δ*pefA* cells over the course of infection. Representative images shown at 2 hpi capture early recruitment events (Fig. 8G, I, and K), whereas quantitative analyses extend to 24 hpi, a time window in which PefA transcriptional upregulation and MCV-associated accumulation have already peaked, as indicated by RNA-seq (Fig. 8B) and localization analyses (Fig. 8C to F). In WT cells, TrafE, Vps32, and Atg8a were robustly recruited and accumulated at the MCV early after infection, whereas Δ*pefA* cells exhibited a marked reduction in the fraction of MCV positive for each reporter throughout the time course (Fig. 8H, J, and L). These defects place PefA upstream of ESCRT and autophagy engagement during infection and are consistent with its Ca^2+^-dependent role at damaged endomembranes. Importantly, re-expression of PefA in Δ*pefA* cells fully restored WT-like recruitment of repair reporters and vacuolar integrity, confirming the specificity of the phenotype (Fig. S7D).

To assess the functional consequences of impaired repair, we next quantified cytosolic access of bacteria using mCherry–Plin as a reporter. Plin positivity, which marks bacteria that fully expose their hydrophobic envelope to the cytosol, revealed a striking genotype-dependent difference (Fig. 8M and N). In Δ*pefA* cells, approximately 70–85% of bacteria were Plin-positive across the infection time course, with little temporal variation, indicating precocious and sustained cytosolic escape. By contrast, in WT cells the fraction of Plin-positive mycobacteria remained low at early time points and increased only gradually over time. Thus, loss of PefA leads to early failure of MCV integrity, whereas WT cells largely confine bacteria within a membrane-bound compartment. Together, these results demonstrate that, similar to what was reported for absence of Tsg101, PefA is required for ESCRT- and autophagy-dependent repair at the MCV and for maintaining bacterial confinement. Consistent with the Ca^2+^/PefA model and the Plin readout, intracellular growth of Mmar in Dd depends on timely ESCRT and autophagy-mediated repair of the MCV membrane.

Together, these data indicate that the Ca^2+^/PefA axis identified in sterile membrane damage is also involved in pathogen-induced injury at the MCV. PefA is transcriptionally induced early during infection, accumulates at the MCV, and is required for timely recruitment of ESCRT and autophagy machineries and preservation of vacuolar integrity and to support intracellular bacteria growth. In the absence of PefA, membrane repair is compromised, leading to premature cytosolic escape of bacteria and altered infection outcomes. Collectively, these findings support a role for PefA as a central Ca^2+^-responsive regulator involved in membrane damage sensing and repair during mycobacterial infection. These findings are integrated into the proposed model (Fig. 9), which summarizes how Ca^2+^ availability and PefA-dependent signaling orchestrate repair pathway hierarchy to determine MCV integrity and infection fate during mycobacterial infection. In this model, extracellular Ca^2+^ availability tunes the kinetics of membrane damage responses through PefA, thereby governing the balance between ESCRT-mediated repair, autophagy engagement, and bacteria containment. Rapid Ca^2+^–PefA–dependent ESCRT activation promotes efficient resealing of limited MCV damage and sustains intravacuolar bacterial growth, whereas impaired Ca^2+^ sensing or loss of PefA shifts outcomes towards premature rupture, cytosolic escape, and xenophagic restriction. Thus, Ca^2+^-controlled repair dynamics emerge as a key determinant of both host resilience and mycobacteria fate.

**Fig. 9.**
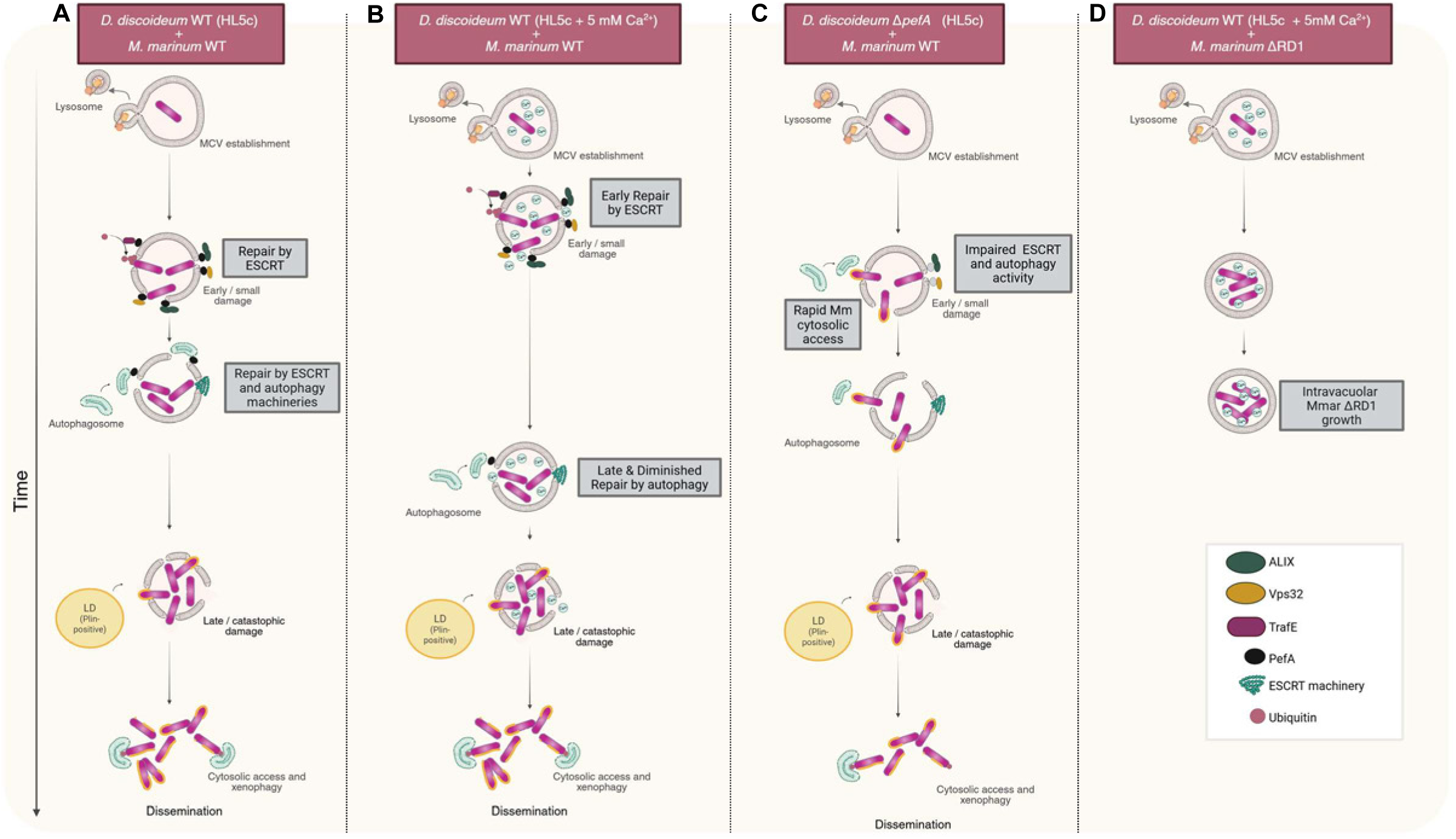
A model for Ca^2+^- and PefA-dependent regulation of membrane repair and mycobacterial fate. Working model illustrating how extracellular Ca^2+^ availability and PefA-dependent signaling modulate the balance between membrane repair, bacterial containment, and intracellular replication during mycobacterial infection. (**A**) Following phagocytosis of Mm, a mycobacteria-containing vacuole (MCV) is formed in Dd. In WT cells cultured in standard medium, early and limited membrane damage to the MCV triggers the recruitment of PefA, ALIX, ESCRT-III (Vps32), and ubiquitin, promoting rapid ESCRT-mediated membrane repair. When damage persists or recurs, autophagy is subsequently engaged, leading to coordinated ESCRT–autophagy repair that preserves MCV integrity, supports intravacuolar bacterial replication, and limits premature cytosolic escape. (**B**) Supplementation with extracellular Ca^2+^ enhances this repair pathway by amplifying early Ca^2+^-dependent signaling, resulting in faster ESCRT recruitment. Consequently, MCV damage is resolved at an earlier stage, autophagy engagement is delayed or reduced, and repair remains predominantly ESCRT driven. This accelerated repair limits catastrophic membrane rupture and confines bacteria within the vacuole for extended periods. (**C**) In contrast, Δ*pefA* cells exhibit impaired early Ca^2+^-dependent damage sensing, leading to inefficient or delayed recruitment of both ESCRT and autophagy machineries. As a result, Mm rapidly gains access to the cytosol, resulting in Plin-positive cytosolic bacteria, engagement of xenophagy, and bacterial restriction rather than productive intravacuolar growth. (**D**) For ESX-1–deficient (ΔRD1) mycobacteria, MCV damage is minimal. Under these low-damage conditions, elevated extracellular Ca^2+^ further stabilizes the MCV and can enhance intravacuolar bacterial replication, consistent with a robust environment that does not require extensive activation of membrane repair pathways. Generated with BioRender.

## Discussion

Cells are continuously exposed to membrane damage caused by mechanical stress, toxins, metabolic by-products, or pathogen-derived activities and must rapidly decide whether to repair, remodel, or eliminate damaged compartments. These decisions occur on short timescales and must preserve ionic homeostasis and cell viability. Here, we identify in Dd Ca^2+^ as a key temporal regulator of membrane repair pathways at damaged lysosomes and MCVs, and define the penta-EF-hand protein PefA as an upstream and central Ca^2+^ sensor that coordinates the hierarchy between ESCRT-mediated repair and autophagy. Using the Dd–Mmar model, we link early Ca^2+^ dynamics to repair pathway engagement and infection outcome through an integrated analysis combining lysosomal physiology, proteomics, genetics, and live infection imaging.

Extracellular Ca^2+^ availability strongly influenced cellular fitness under both sterile and pathogen-induced membrane stress, improving growth, accelerating lysosomal recovery, and reducing cell death (Fig. 1 and Fig. 2). These findings reinforce Ca^2+^ as a global determinant of membrane resilience rather than a simple second messenger, consistent with its conserved role in plasma membrane and endolysosomal repair (8, 12, 37, 48). Importantly, Ca^2+^ did not affect phagocytic uptake or infection prevalence, indicating that its effects manifest downstream of membrane damage rather than at pathogen entry. The differential impact on virulent versus ESX-1-deficient mycobacteria further supports the view that Ca^2+^ signaling becomes decisive specifically when membrane lesions are present.

A major advance of this work is the demonstration that Ca^2+^ regulates both the timing and hierarchical engagement of repair pathways, supporting a kinetic rather than a dose-dependent model. Under Ca^2+^-supplemented conditions, TrafE and ESCRT-III responses to membrane damage were apparently reduced and more transient, coinciding with rapid restoration of lysosomal acidity and cell morphology (Fig. 3). In contrast, Ca^2+^ limitation led to prolonged ESCRT accumulation and increased autophagy engagement (Fig. 5). This behavior is consistent with the concept that Ca^2+^ signals encode information through their amplitude and duration and are interpreted by sensors with distinct affinities and kinetic properties (49, 50). More broadly, Ca^2+^ signaling at damaged membranes likely arises from the coordinated contribution of extracellular Ca^2+^ influx and Ca^2+^ release from internal stores, including ER–endolysosome contact sites. In line with this view, recent work has shown that ER-derived Ca^2+^ release critically shapes lysosomal repair outcomes and ESCRT recruitment, supporting an integrated model of Ca^2+^ microdomain signaling during membrane repair (51). This interpretation is further supported by functional readouts: faster LysoSensor recovery and reduced PI uptake under Ca^2+^ supplementation are indicative of efficient lesion sealing rather than impaired recruitment and function. Accordingly, smaller and shorter-lived ESCRT assemblies reflect successful repair, consistent with studies linking persistent ESCRT accumulation to unresolved damage and progression toward cell death (16, 29, 36).

Autophagy displayed a complementary pattern. Although the fraction of Atg8a-positive cells was largely preserved, Ca^2+^ supplementation reduced and delayed Atg8a accumulation, reflecting more efficient early repair that limits the persistence of membrane damage requiring autophagic engagement, rather than delayed damage accumulation. These findings support a model in which autophagy functions as a secondary repair and containment pathway, mobilized only when ESCRT-mediated sealing is insufficient. This view aligns with recent evidence that the autophagy-related machinery contributes directly to membrane repair rather than serving solely degradative functions (52).

A major advance of our study is the unbiased identification of PefA as a Ca^2+^-responsive interactor shared by TrafE, ESCRT-III, and autophagy (Fig. 4). Among proteins enriched across all three pathways after damage, PefA uniquely belonged to innate defense categories. As an ALG-2–like penta-EF-hand protein with micromolar Ca^2+^ affinity, PefA is well suited to sense transient Ca^2+^ elevations at damaged membranes (14, 45). Loss-of-function analyses showed that Δ*pefA* cells phenocopied Ca^2+^ chelation, displaying impaired TrafE and Vps32 recruitment and compensatory but inefficient autophagy activation. Proteomic mapping further revealed damage-induced remodeling of the PefA interactome toward ESCRT components, v-ATPase subunits, cytoskeletal elements, and lipid metabolism enzymes (Fig. 6), supporting a model in which Ca^2+^-bound PefA acts as a kinetic scaffold organizing transient repair assemblies.

Although stable PefA-Alix interactions were not detected by affinity purification-mass spectrometry, live-cell imaging demonstrated that Alix recruitment to damaged membranes depends on both Ca^2+^ and PefA (Fig. 7), consistent with the conserved ALG-2-ALIX-ESCRT cascade described in mammalian systems and the strictly Ca^2+^-dependent PefA-Alix interaction in Dd (45). The reversible and thus transient nature of Ca^2+^-triggered PEF-Alix interactions likely explains their absence from steady-state proteomics. Rapid PefA recruitment to laser-induced wounds in a calcium-dependent manner further supports its role as a kinetic gatekeeper operating within a narrow temporal window.

A key strength of this work is the demonstration that mechanisms identified under sterile damage are directly relevant during infection. The TrafE–Vps32–Atg8a interactome was robustly induced at the transcripotional level during Mmar infection, with pefA among the earliest upregulated genes. Loss of PefA resulted in premature bacteria access to the cytosol, reduced intracellular replication, and diminished release of bacteria to the extracellular medium, indicating that early vacuolar rupture exposes bacteria to xenophagic restriction (Fig. 8). Importantly, Mmar infection generates a heterogeneous spectrum of intracellular states, ranging from intact or partially damaged MCVs to fully cytosolic bacteria. As ESCRT recruitment is restricted to membrane-bound compartments, population-level measurements during infection necessarily average across these states and therefore underestimate early ESCRT-dependent repair events that occur prior to cytosolic escape. These findings highlight membrane repair as a double-edged process and a potential susceptibility factor in host–pathogen interactions: while it preserves host viability, it can also stabilize pathogen-containing niches. We propose that in wild-type cells, Ca^2+^–PefA-dependent ESCRT engagement maintains the MCV in a state of controlled instability, rapidly sealing small lesions while preserving a permissive replicative environment. Consistently, analogous tuning of vacuolar membrane damage has been reported during ESX-1–dependent mycobacterial infection in mammalian systems, supporting the broader relevance of kinetically regulated repair mechanisms (53, 54). Elevated extracellular Ca^2+^ reinforces this balance, enhancing host survival while inadvertently promoting bacterial persistence, illustrating how Ca^2+^-driven repair pathways can be co-opted by intracellular pathogens to sustain their niche. Taken together, these observations indicate that Ca^2+^–PefA-dependent coordination of ESCRT-mediated repair and autophagy operates across both sterile and mycobacteria-induced membrane damage. These findings raise the possibility that Ca^2+^-dependent coordination of ESCRT and autophagy represents a conserved principle of membrane repair, prompting investigation of whether ALG-2 or other Ca^2+^ sensors execute this function in mammalian cells.

Collectively, our data support a model in which Ca^2+^ dynamics, interpreted through PefA, act as a molecular timer governing the tempo and hierarchy of membrane damage responses (Fig. 9). By accelerating resealing, Ca^2+^ limits prolonged engagement of repair pathways and preserves compartment integrity, whereas failure of this system shifts outcomes toward inefficient autophagy, rupture, and clearance. This framework integrates Ca^2+^ signaling, ESCRT-mediated repair, and autophagy into a dynamic decision-making process shaping both cell fate and infection outcome, with likely relevance to mammalian phagocytes and other intracellular pathogens beyond mycobacteria.

## Materials and Methods

### Dd Strains Culture and Plasmids

Dd strains and plasmids are listed in table S1. Cells were grown axenically at 22°C in HL5c medium (Formedium) supplemented with 100 U/mL of penicillin and 100 μg/mL of streptomycin (Invitrogen). Plasmids were transfected into Dd by electroporation and selected using the appropriate antibiotic. Hygromycin was used at 15 μg/mL for KO cell lines and 50 μg/mL for cell lines with reporters integrated at the safe-haven act5 locus. Blasticidin was used at 5 μg/mL for KO cell lines. To monitor Dd growth, 10^4^ Dd cells expressing cytosolic mCherry were plated in 96-well IBIDI dishes in filtered HL5c were used to quantify Dd growth with Ca^2+^-supplemented medium. Cell imaging was performed with a 60x water immersion objective with the ImageXpress Micro XL HC microscope at 3, 6, 12, 24 and 32 hours after plating (hpi). Analysis per cell was performed to quantify total cell number.

### Dd mutants

Dd Δ*pefA* and Δ*alix* mutant were generated using CRISPR-Cas9 system as previously described ({Sekine, 2018 #2028}). Briefly, the single-guide RNA sequences (table S2) were cloned into the pTM1285 plasmid by Golden-gate assembly using BpiI. Each plasmid was verified by sequencing before 2 µg was electroporated into Dd cells as previously detailed (55). After an over-night recovery, the transfected cells were selected using 10 µg/mL of G418 for 48 hours. The selected transfected cells were then grown few days in absence of G418 before processing to clonal selection on *Klebsiella aerogenes* lawn. Transfected cells were diluted and 1000; 100 and 10 cells were mixed with *K. aerogenes* culture and spread on SM2 agar plates. After 3-4 days at 22°C, plaques of spores appeared on the plates. A dozen of isolated plaques were scraped from the SM2 plates and started in HL5c culture in 24-well plates. To analyze individual clones, a part of the cells coming from each well from the 24-well plates were quickly lysed and the genomic region of interest was amplified by PCR and sequenced. Clones presenting independent mutations were expanded and kept for further analysis.

### Mycobacterial Strains and Culture

Mycobacteria (Mmar WT and Mmar ΔRD1) were grown in Middlebrook 7H9 (Difco) supplemented with 10% OADC (Becton Dickinson), 0.2% glycerol, and 0.05% tyloxapol (Sigma Aldrich) at 32°C with shaking at 150 rpm in the presence of 5 mm glass beads to prevent clumping. Hygromycin was added at a concentration of 100 μg/mL for mCherry/GFP expression, while 25 μg/mL of kanamycin was used for lux expression.

### Induction of Sterile Damage using LLOMe

For LLOMe experiments, 5×10^5^ cells were plated in 96-well IBIDI dishes containing filtered HL5c. To label acidic compartments, 1 μM LysoSensor Green DND-189 (ThermoFisher) was added to the cells 1 hour prior to imaging. For quantification of LysoSensor signal or repair reporters, cells were imaged with a 60x water immersion objective with the ImageXpress Micro XL HC microscope every 5 minutes for 1 hour (LysoSensor) or 1.5 hours (repair reporters). For each condition, 3 wells were imaged with 4 image fields per well acquiring 3 z-sections per field with a 1 µm interval, across the three biological replicates of the experiment. An initial image was acquired for all conditions before addition of LLOMe at the indicated concentration, after which the localization of GFP-TrafE, GFP-Vps32, GFP-Atg8a, GFP-PefA, and Alix-GFP was monitored. For - Ca^2+^ manipulation treatments, cells were pre-incubated for 2 hours in HL5c, HL5c supplemented with 5mM Ca^2+^, or 100 µM BAPTA-AM prior to LLOMe addition. Single-cell analyses were performed to quantify dot formation and area, as well as the cell shape factor. Analysis per image was used to quantify the percentage of dot-positive cells.

### Ca^2+^ measurements

Single-cell live imaging Ca^2+^ assays were performed using Dd constitutively expressing the Ca^2+^ FRET sensor yellow Cameleon (YC) 3.60(38). Briefly, 3 x 10^4^ cells cultured in HL5c or HL5c + 5 mM Ca^2+^ were plated on a CELLview cell culture dishes (Greiner Bio-One North America, Inc). YC 3.60 FRET was measured every 5 sec with CFP and FRET settings where both were excited at 440 and CFP emission measured at 480 ± 40 while FRET channel at 525 ± 40. LLOMe (either at 4.5 mM or 45 mM) was used to induce Ca^2+^ release from endosomes. Analysis was performed using the Trackmate plugin in ImageJ (56).

### Infection Assays

Infections were performed as previously described (57, 58) with a few modifications. Briefly, Mmar WT expressing the lux operon (luxCDABE) under the hsp60 promoter (58, 59)or Mmar WT expressing cytosolic GFP or mCherry was cultured for 24 h prior to the experiment at 32 °C under shaking conditions in 10 – 20 mL 7H9 broth (Becton Dickinson, Difco Middlebrook 7H9) containing 0.2% glycerol (Sigma Aldrich), 10% OADC (Becton Dickinson) and 0.05% tyloxapol (Sigma Aldrich). After Mmar growth, the OD_600_ was measured and adjusted to infect Dd cells plated the day before in at a MOI 10:1. After infection by spinoculation, extracellular bacteria were washed off, and attached infected cells were resuspended in filtered HL5c containing a bacteriostatic dose of 5 μg/mL of streptomycin and 5 U/mL of penicillin to prevent extracellular Mmar growth. Mock-infected cells were treated similarly but without bacteria. To test the mycobacteria lux signal, 3 x 10^4^ Dd cells were plated per well in 96-Well, Nunclon Delta-Treated, Flat-Bottom Microplate (ThermoFisher), using 3 wells per condition in three biological replicates of the experiment. For quantification of the repair reporters, 1-3 x 10^4^ cells were plated per well in 96-well IBIDI dishes, using 3 wells per condition in three biological replicates of the experiment. Plates were sealed with a gas-impermeable membrane (H769.1, Carl Roth) and briefly centrifuged. Luminescence was recorded over 72 h at 25°C with readings taken every hour using an Agilent BioTek H1 plate . For quantification of the repair reporters, cells were imaged using a 60x water immersion objective on an ImageXpress Micro XL HC microscope at the indicated time points, with 3 wells per condition, 9 imaged fields per well, and 3 z-sections per field with a 1 µm interval. Single-cell analysis was performed after segmentation using the MetaExpress software as described (55).

### Proteomic analysis

Dd cells (10^8^ per sample) expressing GFP-TrafE, GFP-Vps32, GFP-Atg8a, GFP-PefA, or cytosolic GFP (control) were left untreated or challenged with 10 mM LLOMe for 10 min to induce endolysosomal damage. Cells were lysed on ice in RIPA buffer (50 mM Tris-HCl pH 7.5, 150 mM NaCl, 0.1% SDS, 2 mM EDTA pH 8.0, 0.5% sodium deoxycholate, 0.5% Triton X-100) supplemented with protease and phosphatase inhibitor cocktails (Roche). Clarified lysates were incubated with 30 µl GFP-Trap® Magnetic Agarose (ChromoTek) for 4 h at 4°C on a wheel, captured on a magnetic rack, and washed 2× in Triton X-100-containing buffer (50 mM Tris-HCl pH 7.5, 150 mM NaCl, 50 mM sucrose, 5 mM EDTA pH 8.0, 0.3% Triton X-100, 1 mM DTT, 5 mM ATP, protease and phosphatase inhibitors) and 3× in the same buffer lacking Triton X-100. Bound proteins were eluted and analysed at the CMU Proteomic Platform (University of Geneva, Switzerland) for LC–MS/MS. Three independent biological replicates (n=3) were processed for each bait and condition (± LLOMe).

### Two-photon FRAP imaging

Two-photon FRAP experiments were performed on a Leica SP8 DIVE microscope (Leica Microsystems) using an HC PL APO CS2 63×/1.40 oil immersion objective and controlled with Leica Application Suite X software. Images were acquired at 512 × 512 pixels with a zoom factor of 2.0 (pixel size ∼68 nm), bidirectional scanning, and a pixel dwell time of ∼6.3 µs. Fluorescence emission was collected using spectral detection. FRAP experiments consisted of 2 pre-bleach frames, 1 bleach iteration, and two post-bleach phases of 10 frames each, with acquisition parameters kept constant. Photobleaching was performed using the multiphoton laser in constant percentage mode, with 40 to 60 laser power during the bleach step.

### RNAseq sample preparation and analysis

Following infection with GFP-expressing bacteria, infected and mock-infected cells were pelleted, resuspended in 500 μl HL5c, passed through 30 μm filters, and sorted by FACS (Beckman Coulter MoFlo Astrios). Gating was performed based on cell size (forward scatter) and granularity (side scatter). Infected (GFP-positive) and non-infected (GFP-negative) sub-populations were defined according to GFP intensity in the FITC channel. Typically, ∼5 x 10^5^ cells of each fraction were collected for RNA isolation. RNA from mock-infected cells or cells infected was extracted at the indicated time points using the Direct-zol RNA MiniPrep kit (Zymo Research) following the manufacturer’s instructions. Quality of RNA libraries, sequencing, and bioinformatic analyses were performed as previously described (60).

### Image analysis and statistical analysis

To determine the percentage of infected Dd cells that were positive for the different reporters, Mmar was segmented based on its fluorescence (mCherry or GFP), and Dd cells were segmented based on the fluorescence channel, depending on expression levels. Reporter recruitment was then assessed by first identifying structures with signal intensity higher than the cytosolic pool and subsequently determining their co-localization with the Mmar mask, defined by pixel overlap with the bacteria. An analogous segmentation approach was used to detect dot formation following LLOMe treatment in Dd cells (in the absence of a bacterial mask). Sample sizes and *p*-values are reported in the corresponding figure legends (n = number of technical replicates; N = number of biological replicates). All statistical analyses were carried out using GraphPad Prism 10 or R pipelines, and data are shown as mean ± SEM. Statistical significance is indicated as follows: * p < 0.05; ** p < 0.01; *** p < 0.001; **** p < 0.0001; “ns” denotes no significant difference.

## Acknowledgments

We gratefully acknowledge Dr. Nicolas Demaurex (University of Geneva) for his suggestions and help on calcium-sensing experiments. We also appreciate the support from the staff at the Proteomics Core Facility (Faculty of Medicine, University of Geneva), special thanks to Dr. Alexandre Hainard (University of Geneva) and Dr. Domitille Schvartz (University of Geneva) for their assistance with protein pull-down experiments and analysis. We also appreciate the support from Dr. Jérôme Bosset at the Photonic Bioimaging Center, University of Geneva. We also thank Dr Lyudmil Raykov, Dr. Nabil Hanna, Dr. Jahn Nitschke for their insights in RNAseq data.

## Author contributions

Conceptualization: SGG, TS

Methodology: SGG, AP, ACS, CM, TS

Data analysis: SGG, AP, TS

Visualization: SGG, TS

Supervision: SGG, TS

Writing—original draft: SGG, TS

Writing—review & editing: SGG, TS

## Data and materials availability

The RNAseq raw data are available in the SRA/GEO Bioproject PRJNA1243452 and the processed data on Zenodo (https://zenodo.org/records/15111828). In addition, the code used for RNAseq analyses is partly available via the UNIGE bioinformatics platform (https://www.unige.ch/medecine/bioinformatics/home), and the code used for DEG analyses is also available on Zenodo (https://zenodo.org/records/15111828). The raw figure data will be available at Zenodo. The vectors created in the present study are available from the corresponding author (TS) upon request. All data and code needed to evaluate and reproduce the results in the paper are present in the paper and/or the Supplementary Materials.

## Supplementary Materials

**Fig S1:**
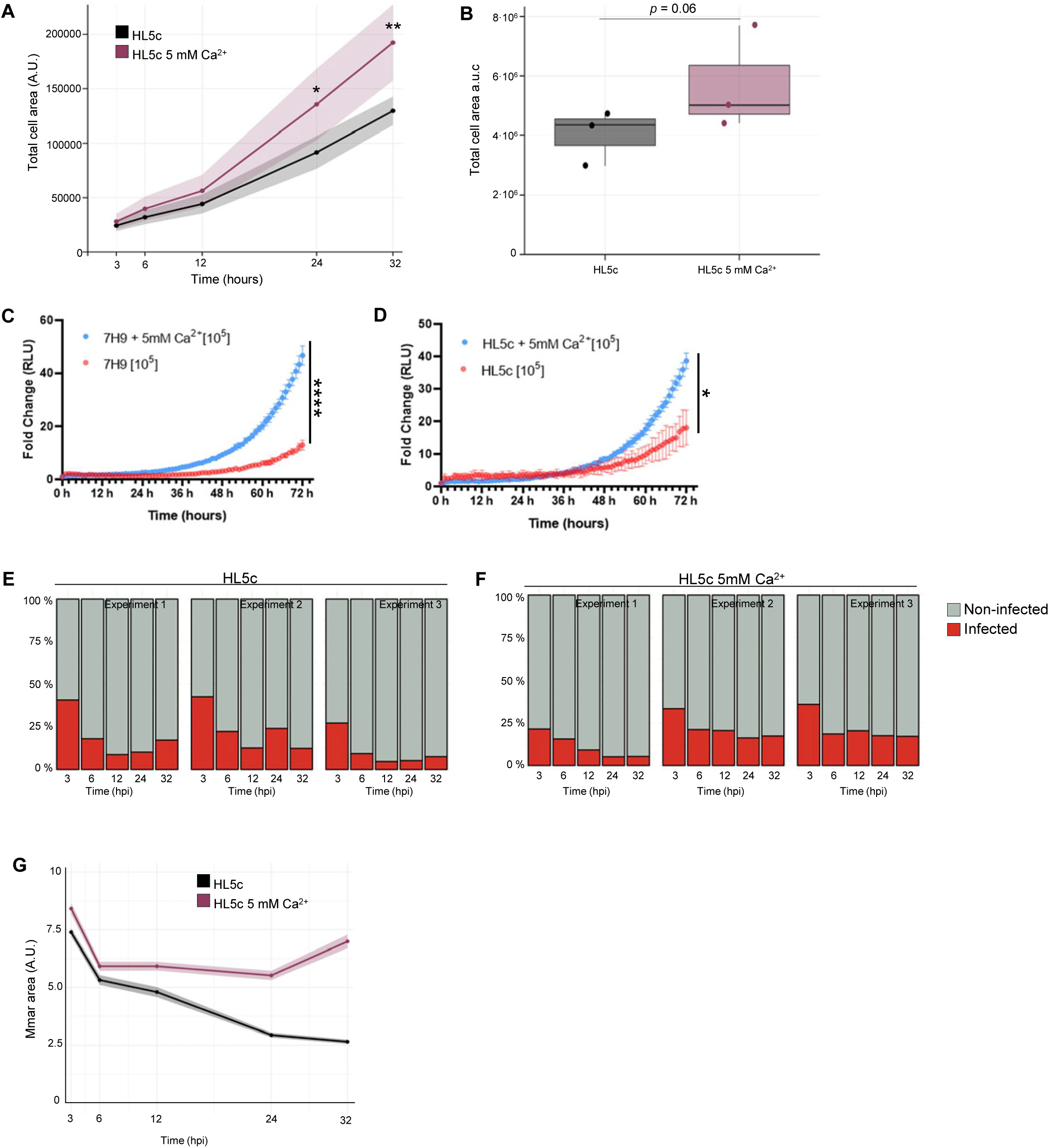
Calcium modulates host cell growth and mycobacterial infection dynamics. **A.** Total Dd cell area over time in HL5c (black) and HL5c + 5 mM Ca^2+^ (magenta); shaded areas indicate SEM. Quantification was performed at single-cell level ( >900 cells per condition: mean ± SEM, n=3; N=3). Statistical comparisons at each time point were performed using a two-sided Welch’s t-test (*p≤0.05, **p≤0.01). **B.** Area under the curve (a.u.c.) of time course shown in (A). Statistical analysis was performed using a two-sided Wilcoxon rank-sum test (Mann–Whitney U) test. *p < 0.05, **p < 0.01. **C.** Growth of 10^5^ cells per well of bioluminescent Mm WT in Middlebrook 7H9 + OADC (red) or 7H9 + OADC supplemented with + 5 mM Ca^2+^ (blue). Bioluminescence was measured for 72 hours (mean fold change ± SEM, n=3, N = 3, two-tailed unpaired t test, (*p≤0.05). RLU, relative luminescence units. **D.** Growth of 10^5^ cells per well of bioluminescent Mm WT in HL5c medium (red) or HL5c supplemented with + 5 mM Ca^2+^ (blue). Bioluminescence was measured for 72 hours (mean fold change ± SEM, n=3, N = 3, two-tailed unpaired t test, ****p≤ 0.0001). **E** and **F.** Proportion of infected cells determined by automated image analysis of Dd cells expressing cytosolic mCherry and infected with GFP-expressing Mmar, quantified over time in HL5c (**E**) or HL5c supplemented with 5 mM Ca^2+^ (**F**), at the indicated times post-infection (>2000 cells per condition; mean ± SEM, n=3; N=3). **G.** Mmar area in Dd infected cells over time in HL5c (black) and HL5c + 5 mM Ca^2+^ (magenta) Mean ± SEM, n=3; N=3.

**Fig. S2:**
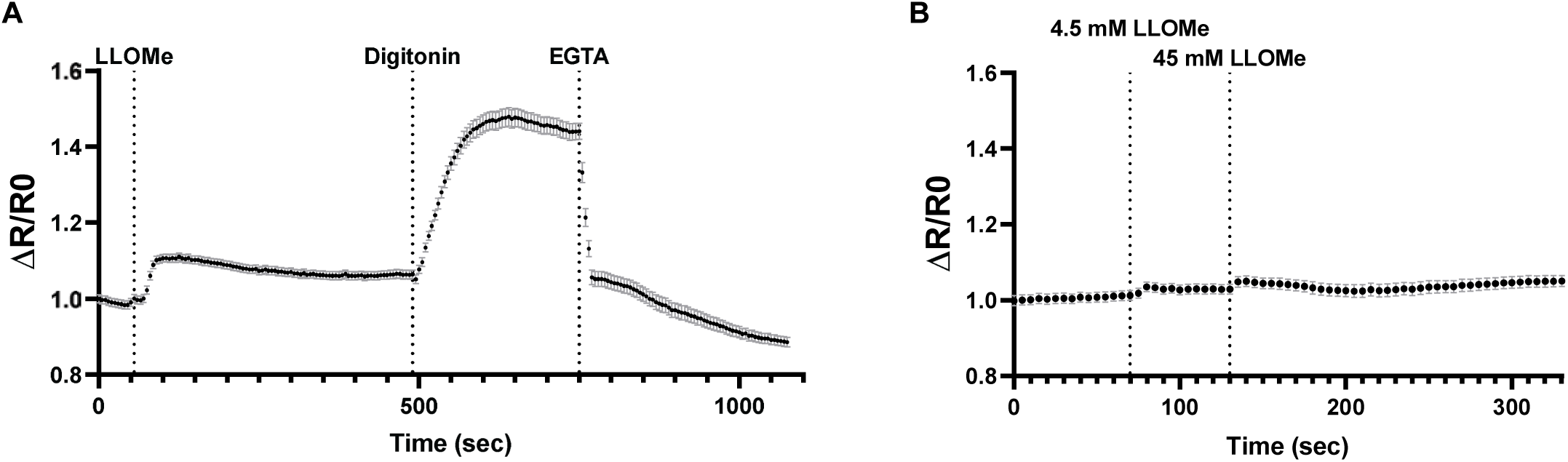
Monitoring intracellular Ca^2+^ levels in Dd using the YC3.60 FRET sensor. **A-B**. One representative experiment showing the normalized FRET ratio of Dd cells expressing the YC3.60 Ca^2+^ FRET sensor cultured in HL5c, imaged every 5 s. (A) Cells were exposed to 4.5 mM LLOMe, followed by 2 µM digitonin and 10 mM EGTA. (mean ± SEM, n=75). (B) Cells were exposed 100 µM BAPTA for 1 hour prior to 4.5 mM LLOMe and 45 mM LLOMe. (mean ± SEM, n=130).

**Fig. S3.**
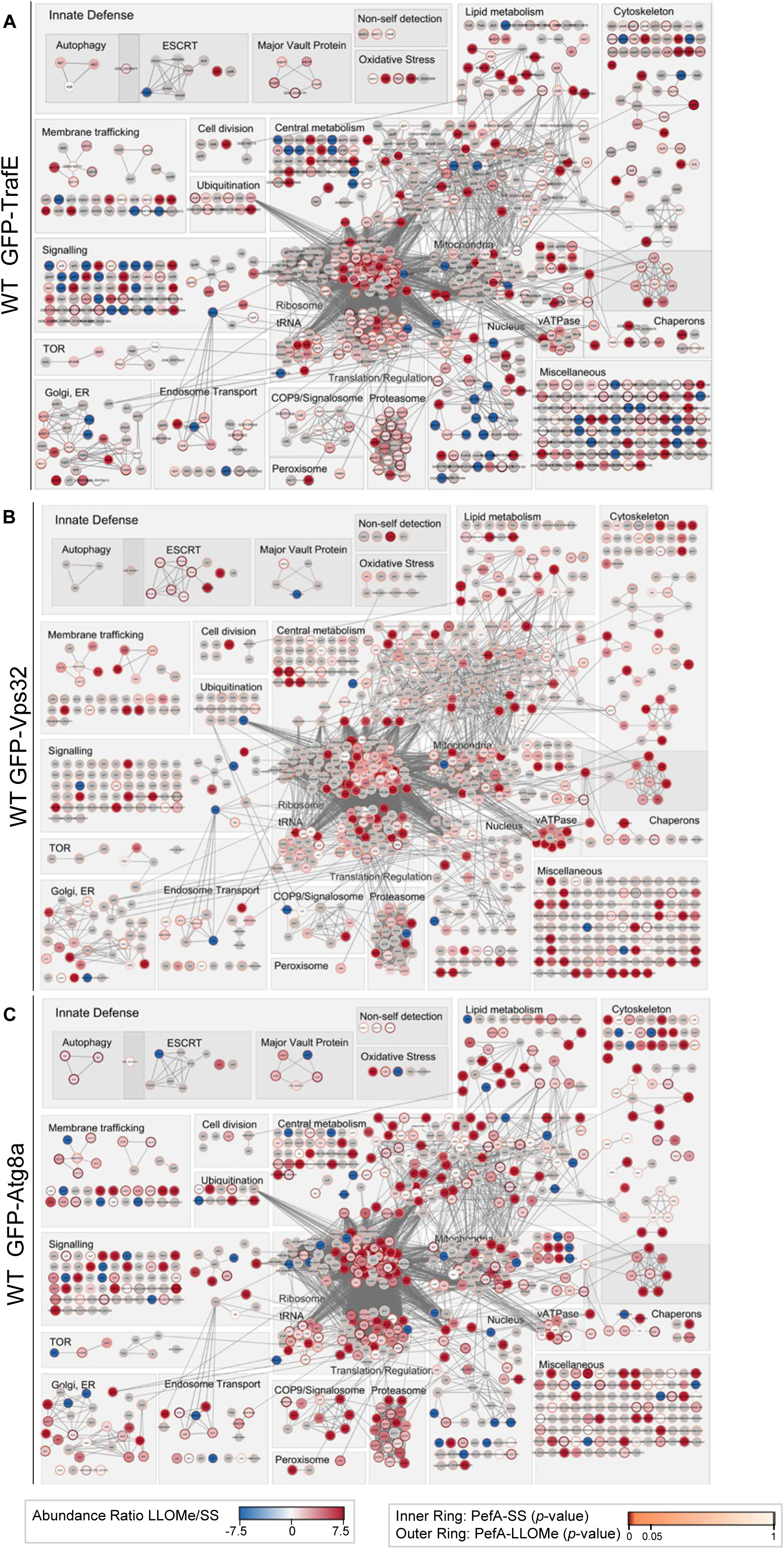
STRING-based network analysis of proteins identified in GFP pull-down assays. **A–C** Proteins identified in GFP–TrafE (A), GFP–Vps32 (B), and GFP–Atg8a (C) pull-down experiments were imported into STRING and visualized using Cytoscape. Each node represents a protein, grouped into manually curated functional categories. Node shading indicates the relative change in abundance (LLOMe versus steady state, SS), while inner and outer rings denote statistical significance at steady state and after lysosomal damage, respectively.

**Fig. S4.**
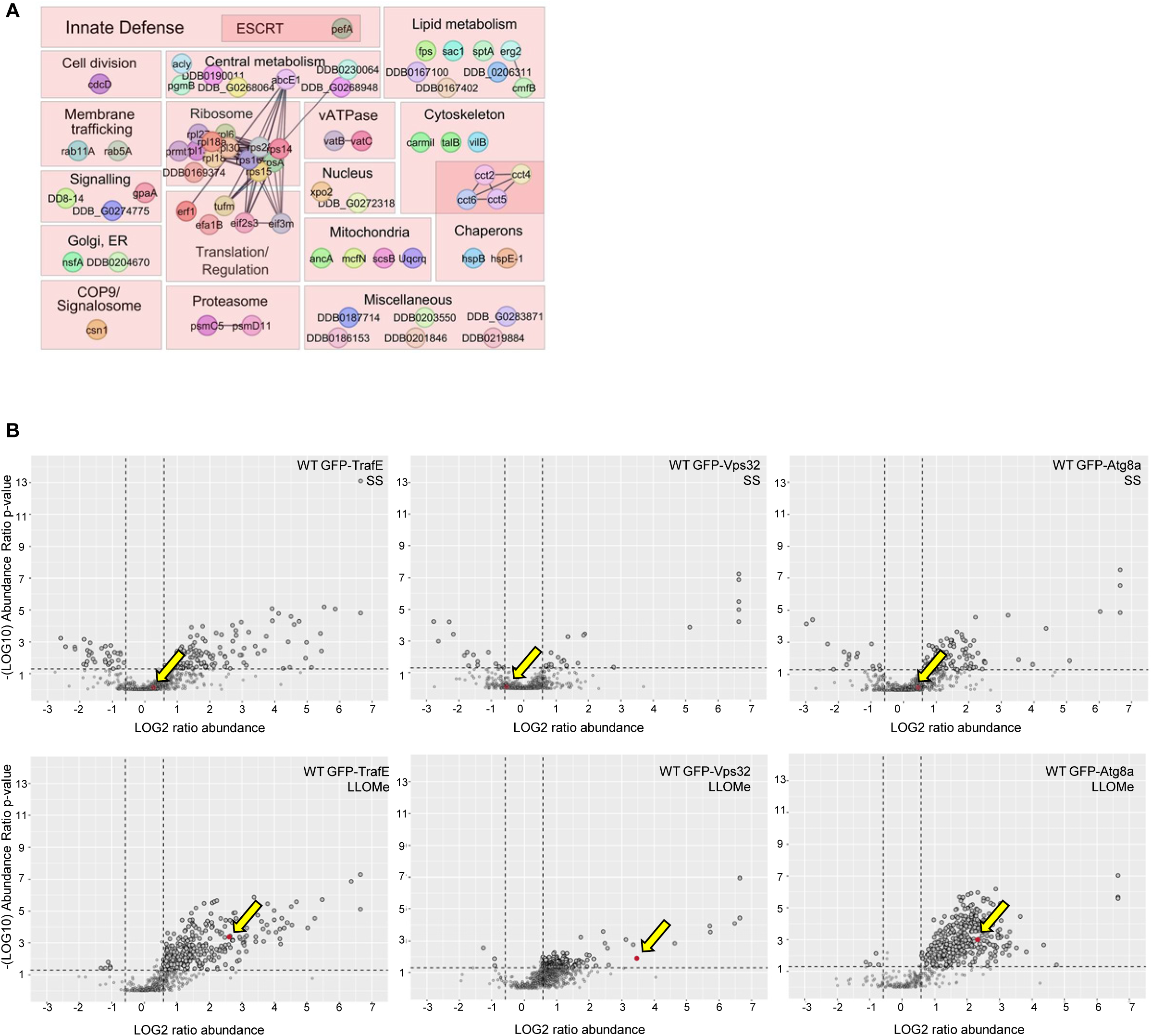
STRING network and volcano plot analysis of TrafE-, Vps32-, and Atg8a-associated proteins. **A.** Distribution of proteins commonly associated with TrafE, Vps32, and Atg8a following LLOMe-induced damage, visualized as a STRING interaction network in Cytoscape. **B**. Volcano plots of GFP pulldown datasets for TrafE, Vps32 and Atg8a under steady-state and LLOMe-treated conditions. Each dot represents a protein, plotted as log₂(fold change over GFP control) versus –log₁₀(p-value). PefA protein is highlighted in red.

**Fig. S5.**
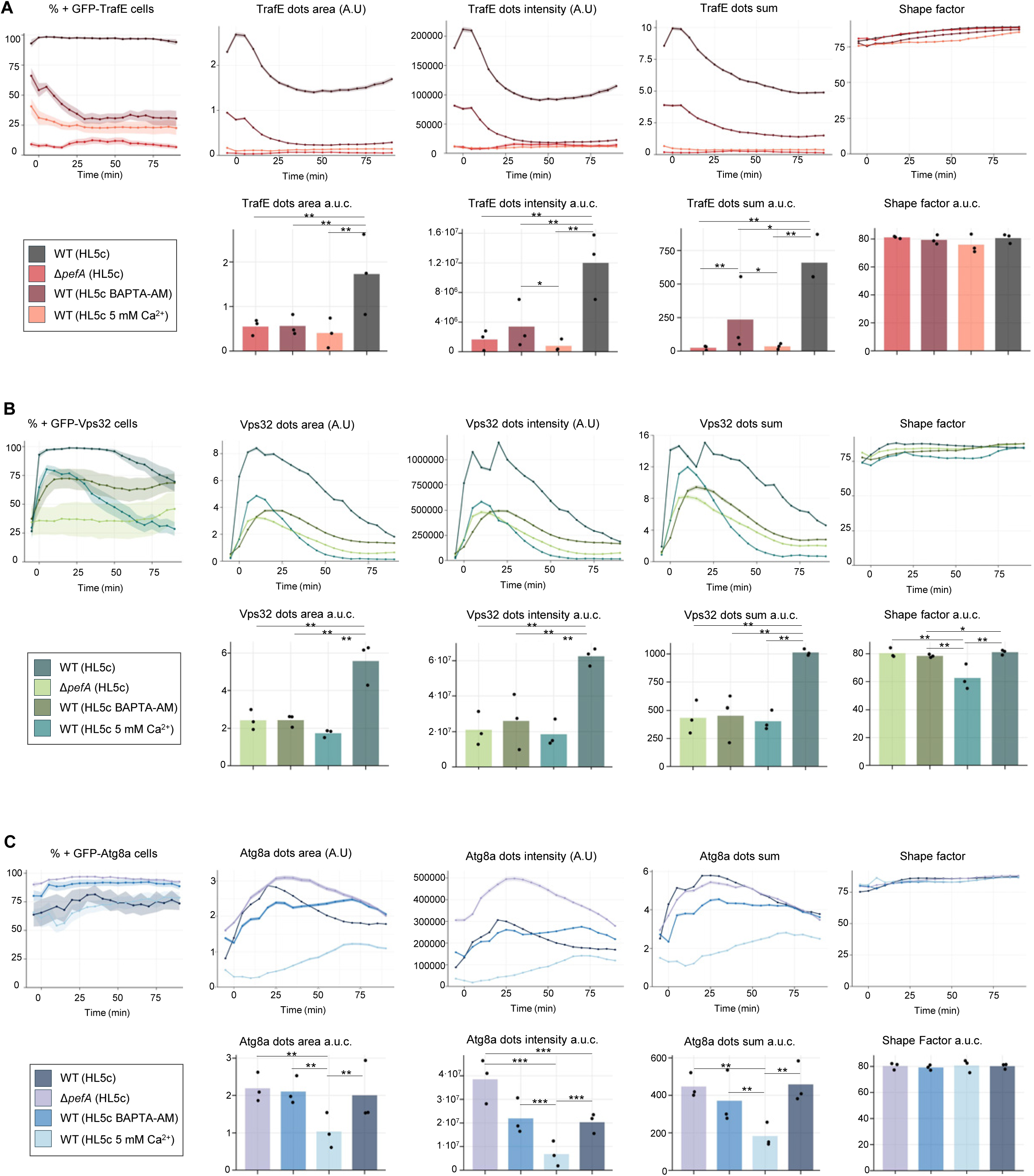
Quantitative analysis of membrane damage repair dynamics under calcium modulation and PefA deletion. **A–C.** Comparative analysis of membrane damage repair dynamics in Dd WT and Δ*pefA* cells under four experimental conditions: WT cultured in HL5c, WT cultured in HL5c supplemented with 5 mM Ca^2+^, WT cultured in HL5c supplemented with BAPTA, and Δ*pefA* cells cultured in HL5c. Cells expressing GFP–TrafE (A), GFP–Vps32 (B), or GFP–Atg8a (C) were treated with 4.5 mM LLOMe to induce lysosomal damage and imaged live by high-content microscopy. For each marker, time-course analyses show the percentage of dot-positive cells, dot area, dot intensity, dot sum, and cell shape factor, as well as the corresponding area under the curve (a.u.c.) for dot area, dot intensity, dot sum, and shape factor. Shaded areas represent mean ± SEM, n**≥**2, N**≥**3. WT cells cultured in HL5c are shown as reference, alongside conditions with elevated extracellular Ca^2+^ (HL5c + 5 mM Ca^2+^), Ca^2+^ chelation (HL5c + BAPTA), or genetic ablation of PefA (Δ*pefA*, HL5c). Statistical significance was assessed using a Kruskal–Wallis test, followed by pairwise two-sided Wilcoxon rank-sum tests with Benjamini–Hochberg correction for multiple comparisons, performed independently for each quantitative parameter (*p ≤0.05, **p ≤0.01, ***p ≤0.005. A.U., arbitrary units.

**Fig. S6.**
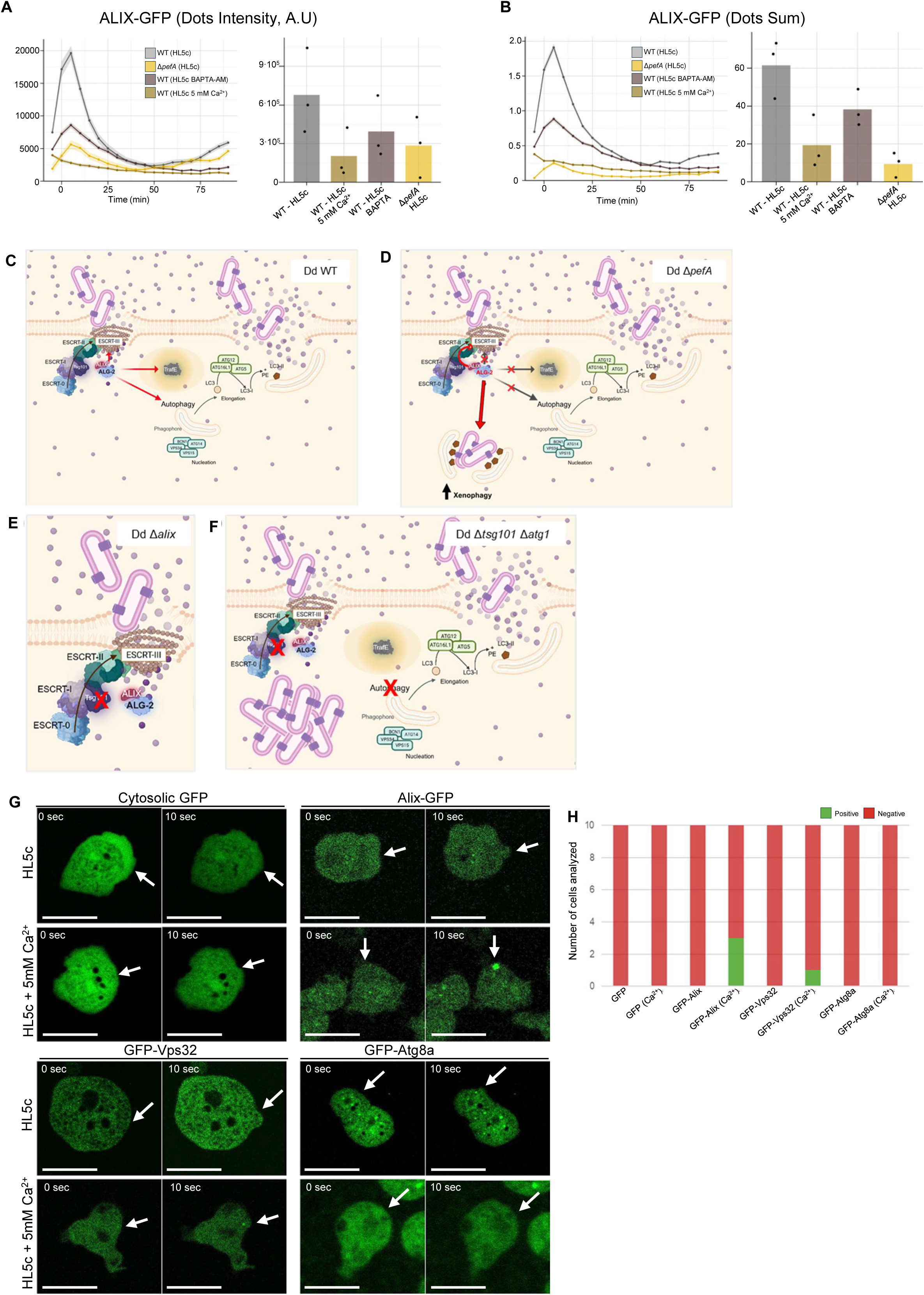
Quantitative analysis and schematic modeling of Ca^2+^- and PefA-dependent membrane repair. **A-B.** Dd WT cultured in HL5c (grey), HL5c + 5 mM Ca^2+^ (light brown) or HL5c + BAPTA (brown); Δ*pefA* Dd cells cultured in HL5c, were treated with 4.5 mM of LLOMe. (A) Dynamics of dots intensity and corresponding area under the curve (a.u.c.). (B) Dynamics of dots sum and corresponding area under the curve (a.u.c.). **C–F**. Schematic representation of how extracellular Ca^2+^ availability and PefA-dependent signaling influence membrane repair, bacterial containment, and intracellular replication in Dd: WT cells (C), Δ*pefA* cells (D), Δ*alix* cells (E), and Δt*sg101/atg1*(F). C. In WT Dd cells, Ca^2+^-dependent activation of ESCRT and autophagy-related pathways promotes efficient membrane repair, limits bacterial escape, and restricts intracellular replication. D. In Δ*pefA* cells, impaired signaling downstream of membrane damage compromises repair efficiency, favoring bacterial escape and enhanced intracellular replication. E. In Δ*alix* cells, defective ESCRT-mediated membrane repair disrupts damage resolution and bacterial containment. F. In Δ*tsg101*/*atg1* cells, combined impairment of ESCRT function and autophagy blocks both membrane repair and xenophagy, leading to uncontrolled bacterial replication. The model integrates genetic perturbations affecting ESCRT and autophagy pathways and supports cell death assays shown in Fig. 7F. **G.** Representative images of Dd cells expressing cytosolic GFP, ALIX-GFP, GFP-Vps32, GFP-Atg8a or cultured in HL5c or HL5c + 5 mM Ca^2+^ subjected to laser-induced plasma membrane damage. Scale bar, 10 µm. **H.** Quantification of positive responses, defined as marker recruitment at the laser-induced injury site (n = 10, N = 1).

**Fig. S7.**
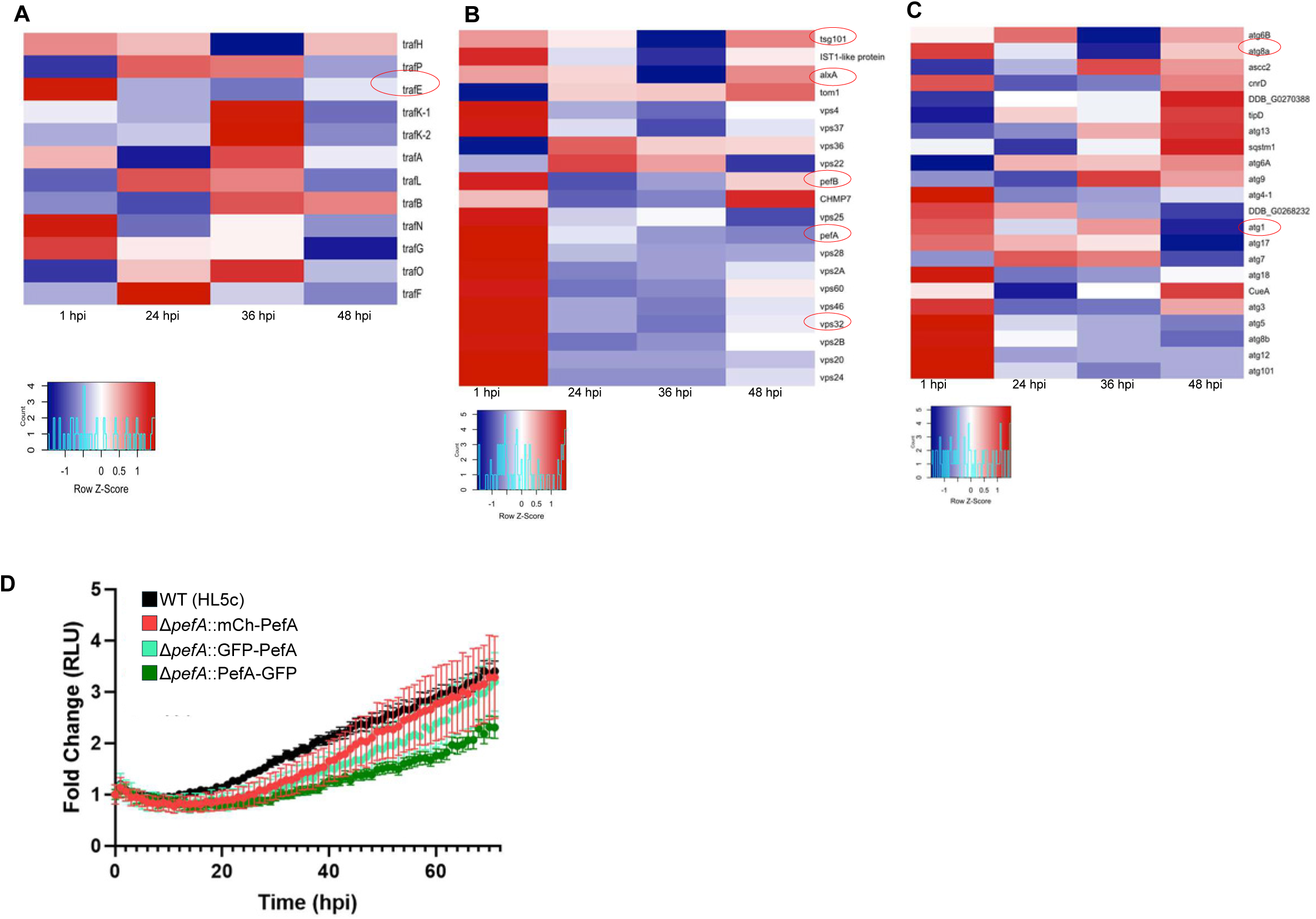
Functional complementation and transcriptomic profiling of host responses during mycobacterial infection. **A.** Dd WT, Δ*pefA* complemented with PefA-GFP, GFP-PefA and mCh-PefA were infected with bioluminescent Mm WT. Bioluminescence was measured for 72 hours (mean fold change ± SEM, n=3, N = 3). RLU, relative luminescence units. **B-D.** Heatmaps of normalized read counts. Filtered and normalized read counts were interrogated for previously defined lists of genes of interest and clustered by conditions. This included a gene set associated to Traf genes (B), the ESCRT machinery (C) and the autophagy pathway (D) Proteins analyzed in this study are highlighted in red.

**Supplementary Table 1.** List of *D. discoideum* and *M. marinum* strains used in this study.

| Strain / Plasmid | Relevant characteristics | Source / Reference |
| --- | --- | --- |
| ***D. discoideum*** |  |  |
| Ax2(Ka) | Wild-type |  |
| Ax2(Ka) GFP-TrafE | pDM1513-TrafE: KI in the Act5 locus | (36) |
| Ax2(Ka) GFP-Vps32 | pDM1513-Vps32: KI in the Act5 locus | (36) |
| Ax2(Ka) GFP-Atg8a | pDM1513-Atg8a: KI in the Act5 locus | (36) |
| Ax2(Ka) ALIX-GFP | pDM1515-ALIX: KI in the Act5 locus | (36) |
| Ax2(Ka) mCherry-Plin | pDM1514-Plin: KI in the Act5 locus | (36) |
| Ax2(Ka) $\Delta pefA$ knock-out strain | | This study |
| Ax2(Ka) $\Delta pefA$ knock-out GFP-TrafE | pDM1513-TrafE: KI in the Act5 locus | This study |
| Ax2(Ka) $\Delta pefA$ knock-out GFP-Vps32 | pDM1513-Vps32: KI in the Act5 locus | This study |
| Ax2(Ka) $\Delta pefA$ knock-out GFP-Atg8a | pDM1513-Atg8a: KI in the Act5 locus | This study |
| Ax2(Ka) $\Delta pefA$ knock-out ALIX-GFP | pDM1515-ALIX: KI in the Act5 locus | This study |
| Ax2(Ka) $\Delta pefA$ knock-out mCherry-Plin | pDM1514-Plin: KI in the Act5 locus | This study |
| ***M. marinum*** |  |  |
| *M. marinum* M strain | Wild-type | L. Ramakrishnan (Washington University) |
| *M. marinum* $\Delta RD1$ | | L. Ramakrishnan (Washington University) |
| **Mycobacteria plasmids** |  |  |
| pCherry10 | mCherry under control of the G13 promoter, Hygr | (Carroll et al., 2010) |
| pMSP12::GFP | GFP under control of the MSP promoter, Kanr | (Cosma et al., 2004) |
| pMV306::lux | bacterial luciferase under control of the hsp60 promoter, Kanr | (Andreu et al., 2010) |

**Supplementary Table 2.**
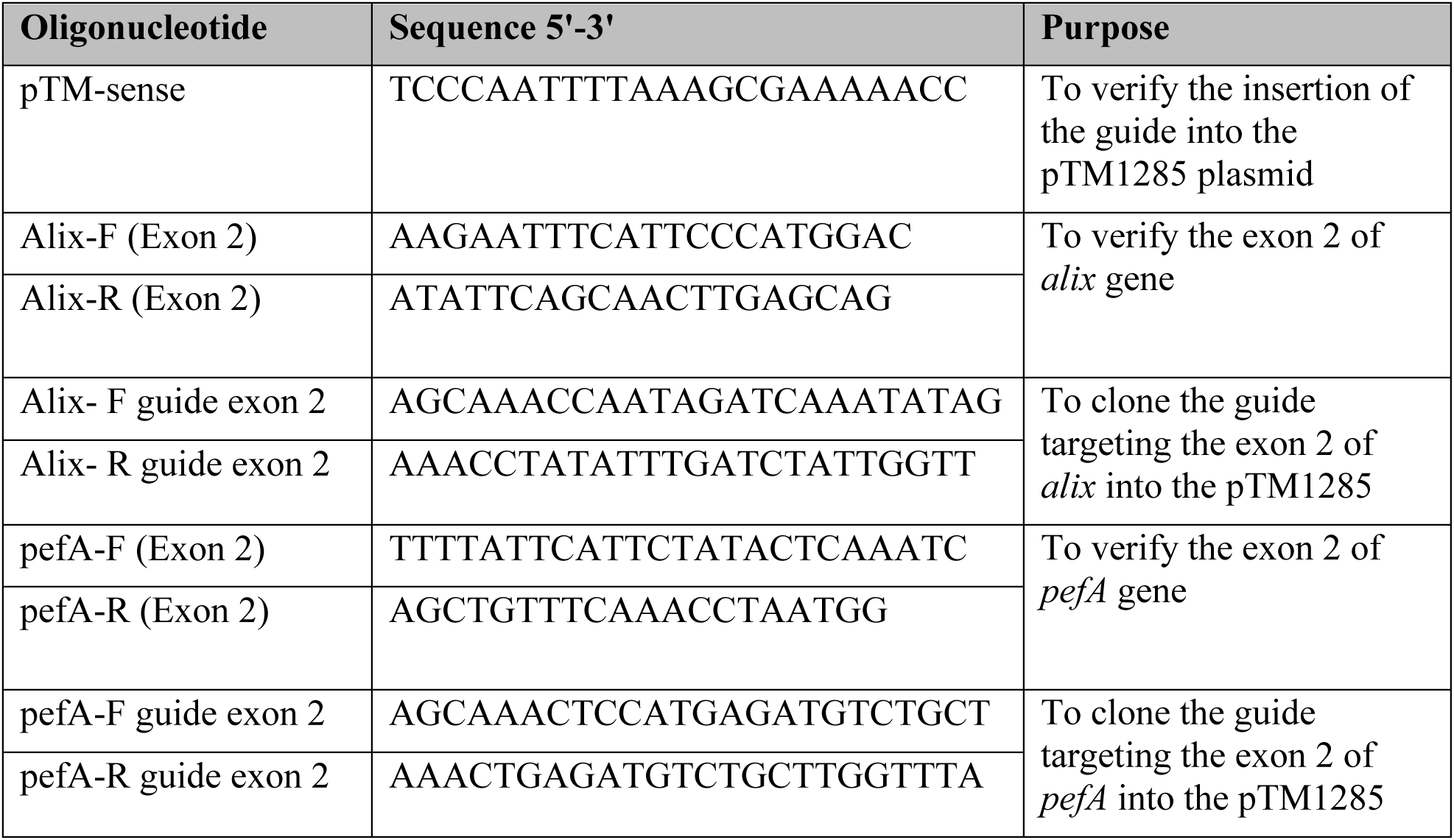
List of oligos used in this study, F: forward, R: reverse.

**Supplementary Table 3.** Proteins significantly enriched in both PefA pull-downs and at least one additional bait (TrafE, Vps32, or Atg8a).

| Prot Id | Protein Name | Condition | Pulldown |  |  |  |
| --- | --- | --- | --- | --- | --- | --- |
|  |  |  | PefA | TrafE | Vps32 | Atg8a |
| Q94469 | Glyceraldehyde-3-phosphate dehydrogenase | SS | x | x |  | x |
| Q54TY4 | Lysine--tRNA ligase | SS | x |  |  | x |
| Q55EN7 | Magnesium-transporting ATPase, P-type 1 | SS | x | x |  |  |
| Q86KU2 | Probable valine--tRNA ligase, cytoplasmic | SS | x |  |  | x |
| P36411 | Ras-related protein Rab-7a | SS | x |  |  | x |
| Q55C75 | 26S proteasome non-ATPase regulatory subunit 6 | LLOMe | x | x |  | x |
| Q58A41 | AAA ATPase domain-containing protein | SS | x | x |  |  |
|  |  | LLOMe | x | x | x | x |
| Q54NP6 | Alpha-soluble NSF attachment protein | LLOMe | x |  |  | x |
| P34121 | Coactosin | LLOMe | x |  | x |  |
| Q54YD8 | Coatomer subunit beta' | LLOMe | x |  |  | x |
| Q54HG2 | Cortexillin-1 | SS | x | x |  |  |
|  |  | LLOMe | x | x |  | x |
| Q550R2 | Cortexillin-2 | LLOMe | x | x |  | x |
| Q54TR1 | Counting factor associated protein D | LLOMe | x | x |  | x |
| O21042 | Cytochrome c oxidase subunit 1+2 | LLOMe | x | x |  |  |
| P02887 | Discoidin-1 subunit B/C | LLOMe | x | x |  |  |
| Q557J6 | Drebrin-like protein | SS | x | x |  |  |
|  |  | LLOMe | x | x |  | x |
| Q769F6 | Emp24/gp25L/p24 family protein | LLOMe | x |  |  | x |
| Q54QW1 | Eukaryotic translation initiation factor 3 subunit B | LLOMe | x | x |  | x |
| Q54KZ8 | Eukaryotic translation initiation factor 3 subunit M | LLOMe | x | x | x | x |
| B0G141 | Glutathione S-transferase domain-containing protein | LLOMe | x | x |  | x |
| P33519 | GTP-binding nuclear protein Ran | SS | x |  |  | x |
|  |  | LLOMe | x | x |  | x |
| P16894 | Guanine nucleotide-binding protein alpha-1 subunit | LLOMe | x | x | x | x |
| P36415 | Heat shock cognate 70 kDa protein 1 | LLOMe | x | x | x | x |
| Q54XX3 | Large ribosomal subunit protein uL18 | SS | x |  |  | x |
|  |  | LLOMe | x | x |  | x |
| P13023 | Large ribosomal subunit protein uL2 | SS | x |  |  | x |
|  |  | LLOMe | x | x |  | x |
| P16168 | Large ribosomal subunit protein uL5 | SS | x |  | x | x |
|  |  | LLOMe | x | x |  | x |
| Q54XI5 | Large ribosomal subunit protein uL6 | LLOMe | x | x |  | x |
| Q54F93 | Mitochondrial-processing peptidase subunit alpha-2 | SS | x | x |  |  |
|  |  | LLOMe | x |  | x | x |
| Q54F93 | Mitochondrial-processing peptidase subunit alpha-2 | LLOMe | x |  | x | x |
| **Q95YL5** | **Penta-EF hand domain-containing protein 1** | **LLOMe** | **x** | **x** | **x** | **x** |
| Q86JG3 | PH domain-containing protein | LLOMe | x | x |  | x |
| Q86IV4 | PH domain-containing protein DDB_G0274775 | LLOMe | x | x | x | x |
| Q54BM2 | Polyadenylate-binding protein 1-A | LLOMe | x | x |  | x |
| Q558Z0 | Probable arginine--tRNA ligase, cytoplasmic | SS | x | x |  |  |
|  |  | LLOMe | x | x |  | x |
| Q54UQ2 | Probable phosphoglucomutase-2 | LLOMe | x | x | x | x |
| P36418 | Protovillin | LLOMe | x | x | x | x |
| P20790 | Ras-related protein Rab-8A | SS | x |  | x | x |
|  |  | LLOMe | x | x |  | x |
| P34143 | Ras-related protein RabC | LLOMe | x | x |  | x |
| P34144 | Rho-related protein rac1A | SS | x |  |  | x |
|  |  | LLOMe | x |  |  | x |
| P90529 | RNA helicase (Fragment) | SS | x |  |  | x |
|  |  | LLOMe | x | x |  | x |
| Q54TD3 | T-complex protein 1 subunit epsilon | LLOMe | x | x | x | x |
| P32255 | Tubulin alpha chain | LLOMe | x | x |  | x |
| P32256 | Tubulin beta chain | LLOMe | x | x |  | x |
| Q54G31 | Uncharacterized protein | LLOMe | x | x |  |  |
| Q54UX5 | Uncharacterized protein | LLOMe | x | x |  | x |
| Q54UM3 | Uridine phosphorylase | LLOMe | x | x |  | x |
| Q54DL7 | von Willebrand factor A domain-containing protein | LLOMe | x | x | x |  |
| Q76NU1 | V-type proton ATPase subunit B | LLOMe | x | x | x | x |
| O00780 | V-type proton ATPase subunit E | LLOMe | x |  | x |  |

**Supplementary Table 4.** Proteins significantly enriched exclusively in PefA and PefA pull-downs and additional baits (TrafE, Vps32, or Atg8a).

|  | Accession | Description |
| --- | --- | --- |
| **Exclusive PefA-LLOMe** |  |  |
| 1 | Q54WC4 | Uncharacterized protein |
| 2 | Q54DC6 | AIG1-type G domain-containing protein |
| 3 | P35401 | Protein CRAC |
| 4 | Q55FF2 | Arrestin C-terminal-like domain-containing protein |
| 5 | Q55FV8 | Transmembrane protein 144 homolog A |
| 6 | Q86KU2 | Probable valine--tRNA ligase, cytoplasmic |
| 7 | Q55EN7 | Magnesium-transporting ATPase, P-type 1 |
| 8 | Q54F95 | Band 7 domain-containing protein |
| 9 | Q54TZ4 | Galactose oxidase |
| 10 | P02888 | Discoidin-1 subunit D |
| 11 | Q54XE6 | Probable H/ACA ribonucleoprotein complex subunit 1 |
| 12 | Q54WN8 | Uncharacterized protein DDB_G0279527 |
| 13 | P34148 | Rho-related protein racB |
| 14 | Q86KX9 | J domain-containing protein |
| 15 | P13021 | F-actin-capping protein subunit beta |
| 16 | Q94469 | Glyceraldehyde-3-phosphate dehydrogenase |
| 17 | Q75JQ1 | Aspartate--tRNA ligase, cytoplasmic 1 |
| 18 | Q9U9A3 | Serine/threonine-protein phosphatase 6 catalytic subunit |
| 19 | P0CT31 | Elongation factor 1-alpha |
| 20 | Q54PA9 | Ribose-phosphate pyrophosphokinase A |
| 21 | Q9XYS3 | Superoxide-generating NADPH oxidase heavy chain subunit A |
| 22 | Q54TR3 | 39S ribosomal protein L28, mitochondrial |
| 23 | Q54Y98 | Transformer-2 protein homolog |
| **Common between PefA - Vps32 LLOMe** |  |  |
| 1 | P34121 | Coactosin |
| 2 | O00780 | V-type proton ATPase subunit E |
| **Common between PefA - TrafE LLOMe** |  |  |
| 1 | Q54G31 | Uncharacterized protein |
| 2 | P02887 | Discoidin-1 subunit B/C |
| 3 | O21042 | Cytochrome c oxidase subunit 1+2 |

| Common between PefA - Atg8a LLOMe |  |  |
| --- | --- | --- |
| 1 | Q54NP6 | Alpha-soluble NSF attachment protein |
| 2 | Q54YD8 | Coatomer subunit beta' |
| 3 | P34144 | Rho-related protein rac1A |
| 4 | Q769F6 | Emp24/gp25L/p24 family protein |
| Common between PefA - Vps32 - TrafE LLOMe |  |  |
| 1 | Q54DL7 | von Willebrand factor A domain-containing protein DDB_G0292188 |
| Common between PefA - Vps32 - Atg8a LLOMe |  |  |
| 1 | Q54F93 | Mitochondrial-processing peptidase subunit alpha-2 |
| Common between PefA - TrafE - Atg8a LLOMe |  |  |
| 1 | Q54QW1 | Eukaryotic translation initiation factor 3 subunit B |
| 2 | P32256 | Tubulin beta chain |
| 3 | Q558Z0 | Probable arginine--tRNA ligase, cytoplasmic |
| 4 | Q54BM2 | Polyadenylate-binding protein 1-A |
| 5 | P16168 | Large ribosomal subunit protein uL5 |
| 6 | Q55C75 | 26S proteasome non-ATPase regulatory subunit 6 |
| 7 | Q54UM3 | Uridine phosphorylase |
| 8 | B0G141 | Glutathione S-transferase domain-containing protein |
| 9 | Q54XX3 | Large ribosomal subunit protein uL18 |
| 10 | Q54UX5 | Uncharacterized protein |
| 11 | P20790 | Ras-related protein Rab-8A |
| 12 | P34143 | Ras-related protein RabC |
| 13 | P13023 | Large ribosomal subunit protein uL2 |
| 14 | P90529 | RNA helicase (Fragment) |
| 15 | P32255 | Tubulin alpha chain |
| 16 | Q550R2 | Cortexillin-2 |
| 17 | P33519 | GTP-binding nuclear protein Ran |
| 18 | Q86JG3 | PH domain-containing protein |
| 19 | Q54TR1 | Counting factor associated protein D |
| 20 | Q54XI5 | Large ribosomal subunit protein uL6 |
| 21 | Q557J6 | Drebrin-like protein |
| 22 | Q54HG2 | Cortexillin-1 |
| Common between PefA - TrafE - Atg8a - Vps32 LLOMe |  |  |
| 1 | Q58A41 | AAA ATPase domain-containing protein |
| **2** | P36418 | Protovillin |
| **3** | Q54TD3 | T-complex protein 1 subunit epsilon |
| **4** | Q54UQ2 | Probable phosphoglucomutase-2 |
| **5** | Q86IV4 | PH domain-containing protein DDB_G0274775 |
| **6** | Q95YL5 | Penta-EF hand domain-containing protein 1 |
| **7** | Q54KZ8 | Eukaryotic translation initiation factor 3 subunit M |
| **8** | P16894 | Guanine nucleotide-binding protein alpha-1 subunit |
| **9** | P36415 | Heat shock cognate 70 kDa protein 1 |
| **10** | Q76NU1 | V-type proton ATPase subunit B |

**Supplementary Table 5.**
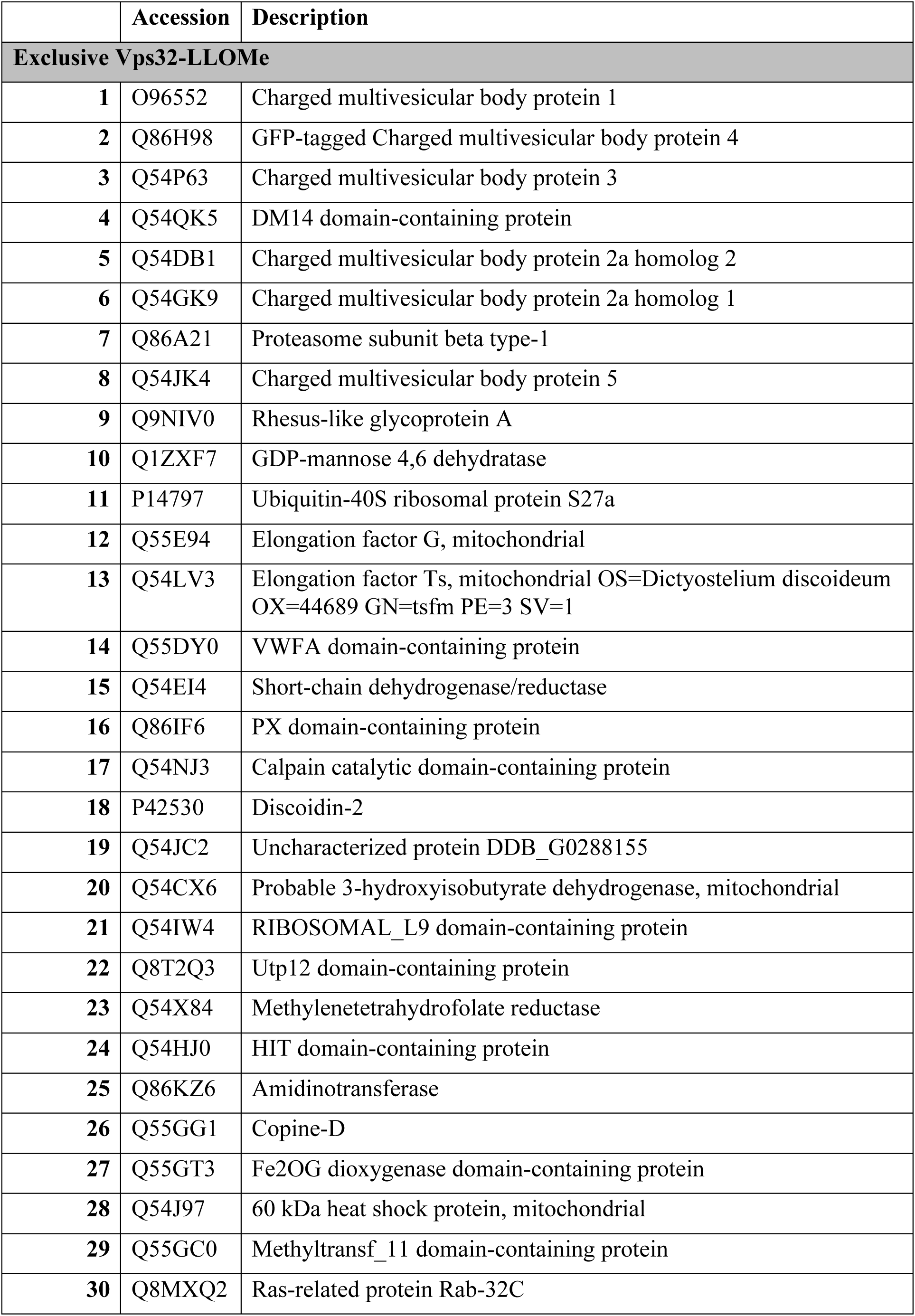
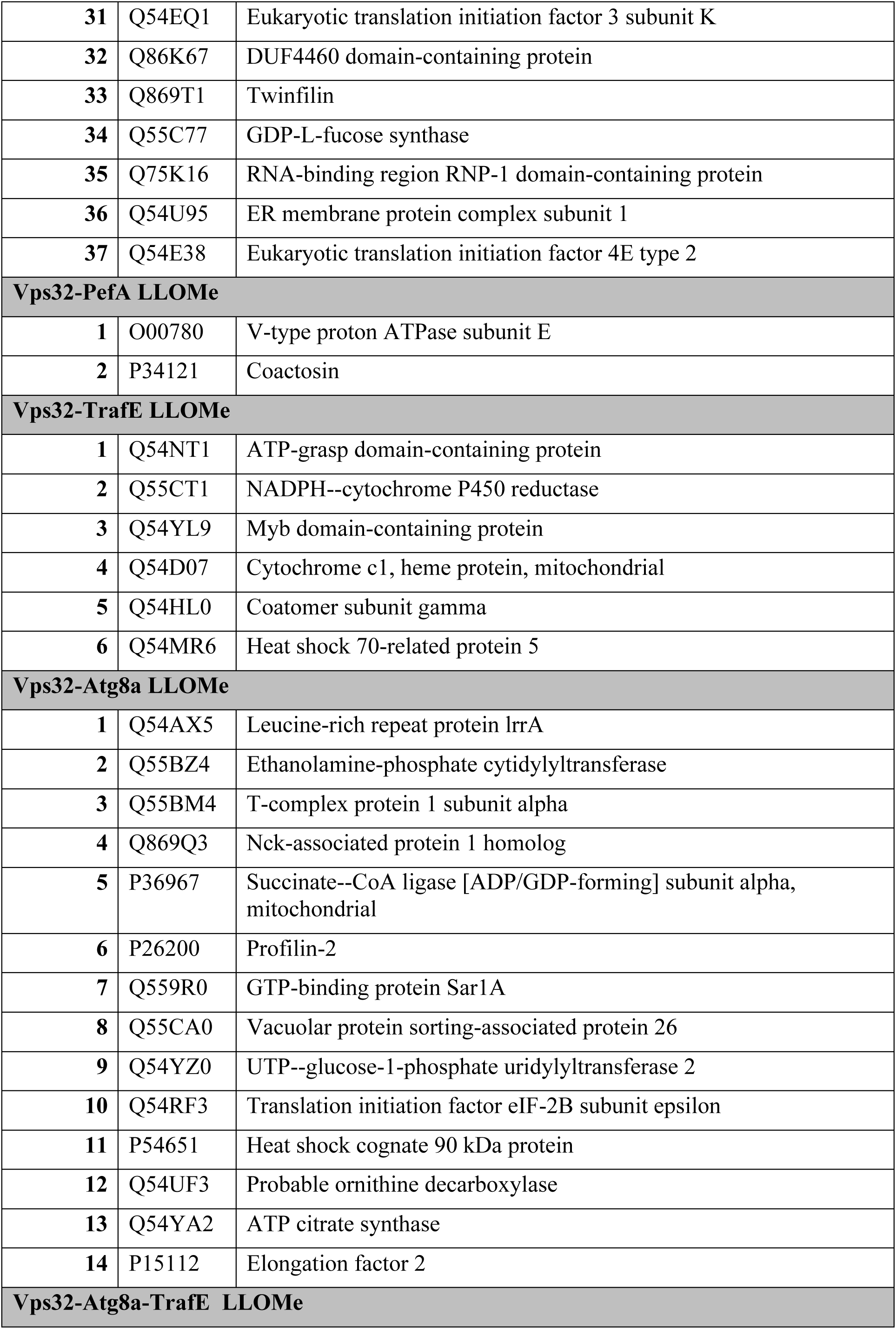
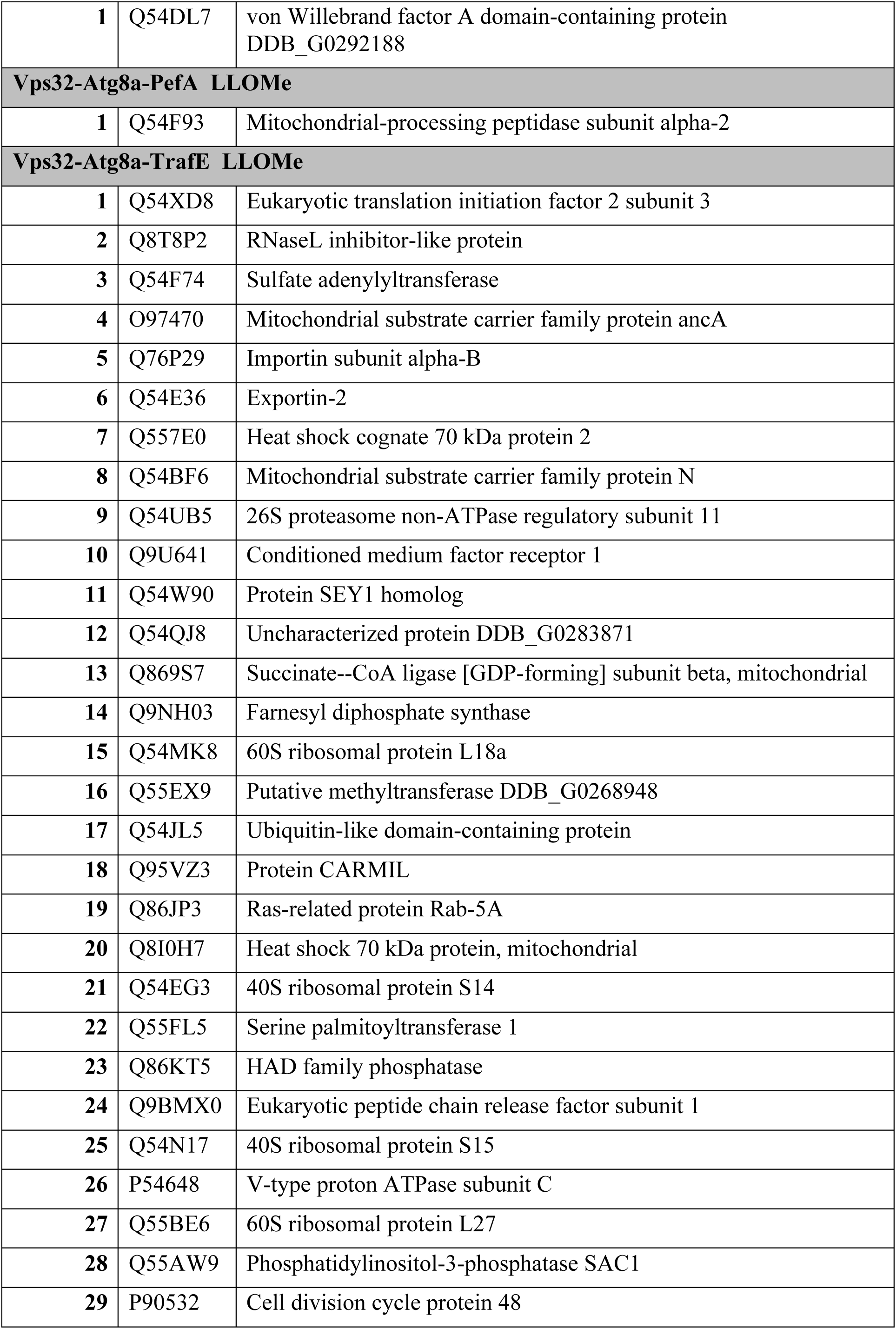
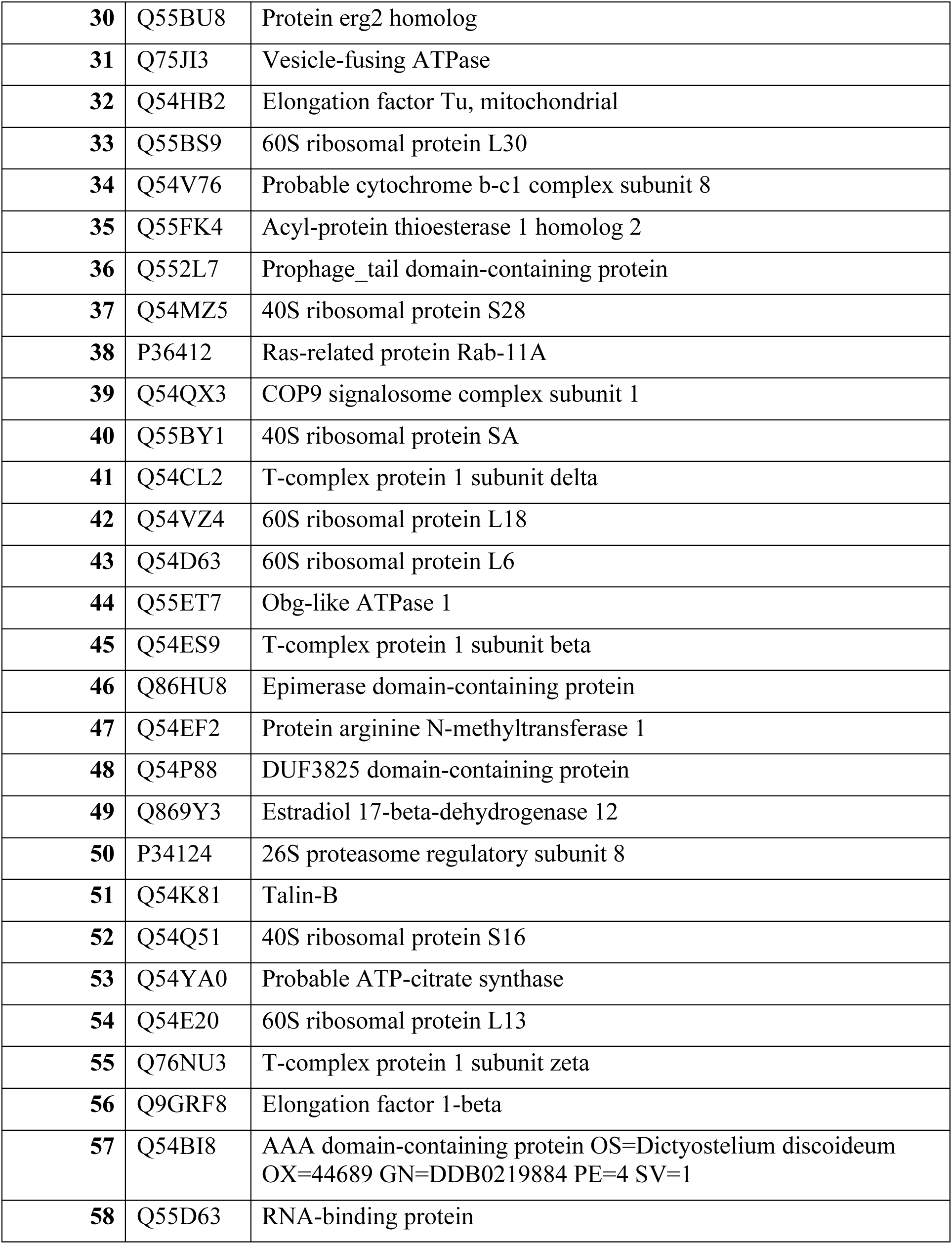
Proteins significantly enriched exclusively in Vps32 and Vps32 pull-downs and additional baits.

**Supplementary Table 6.**
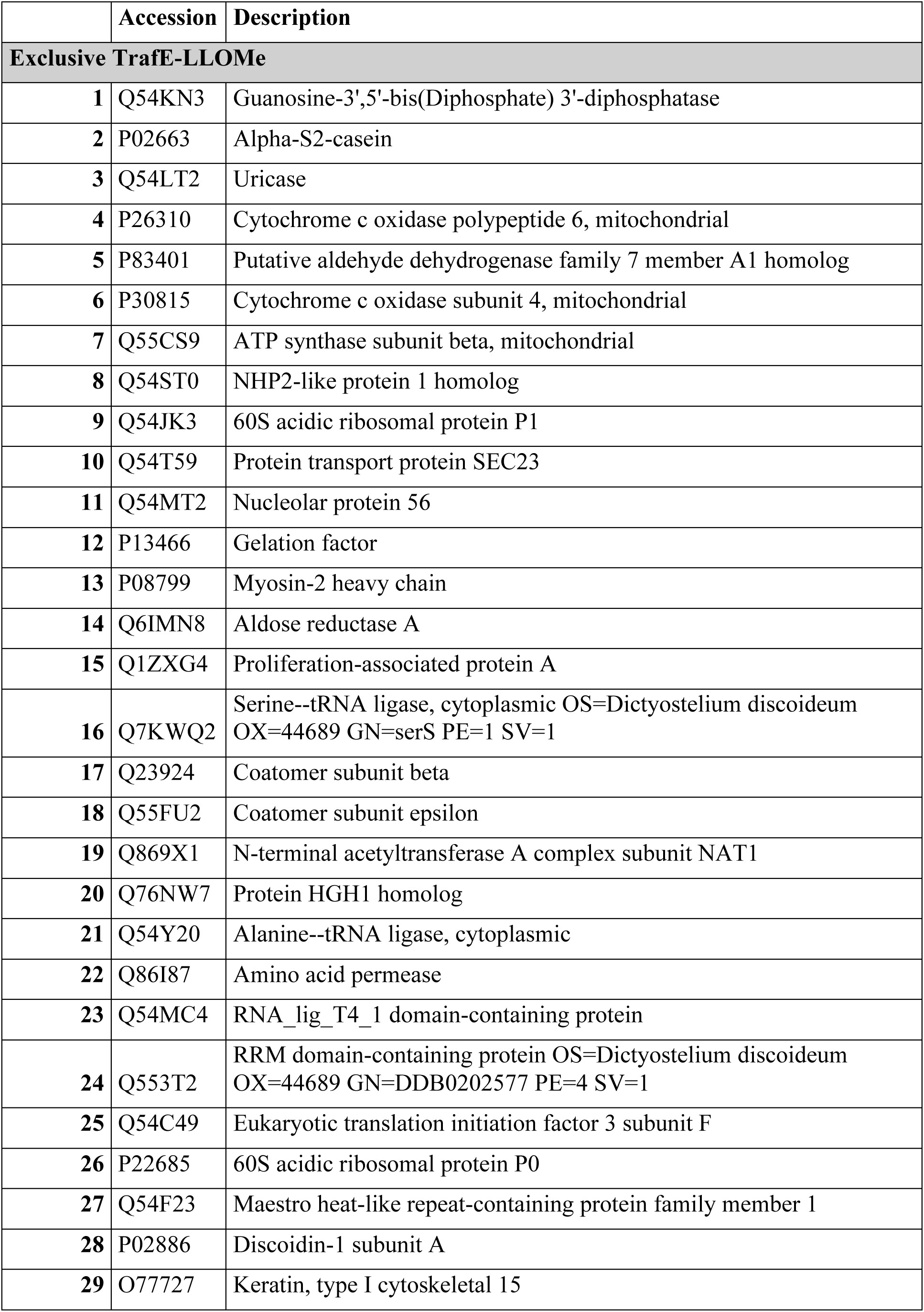
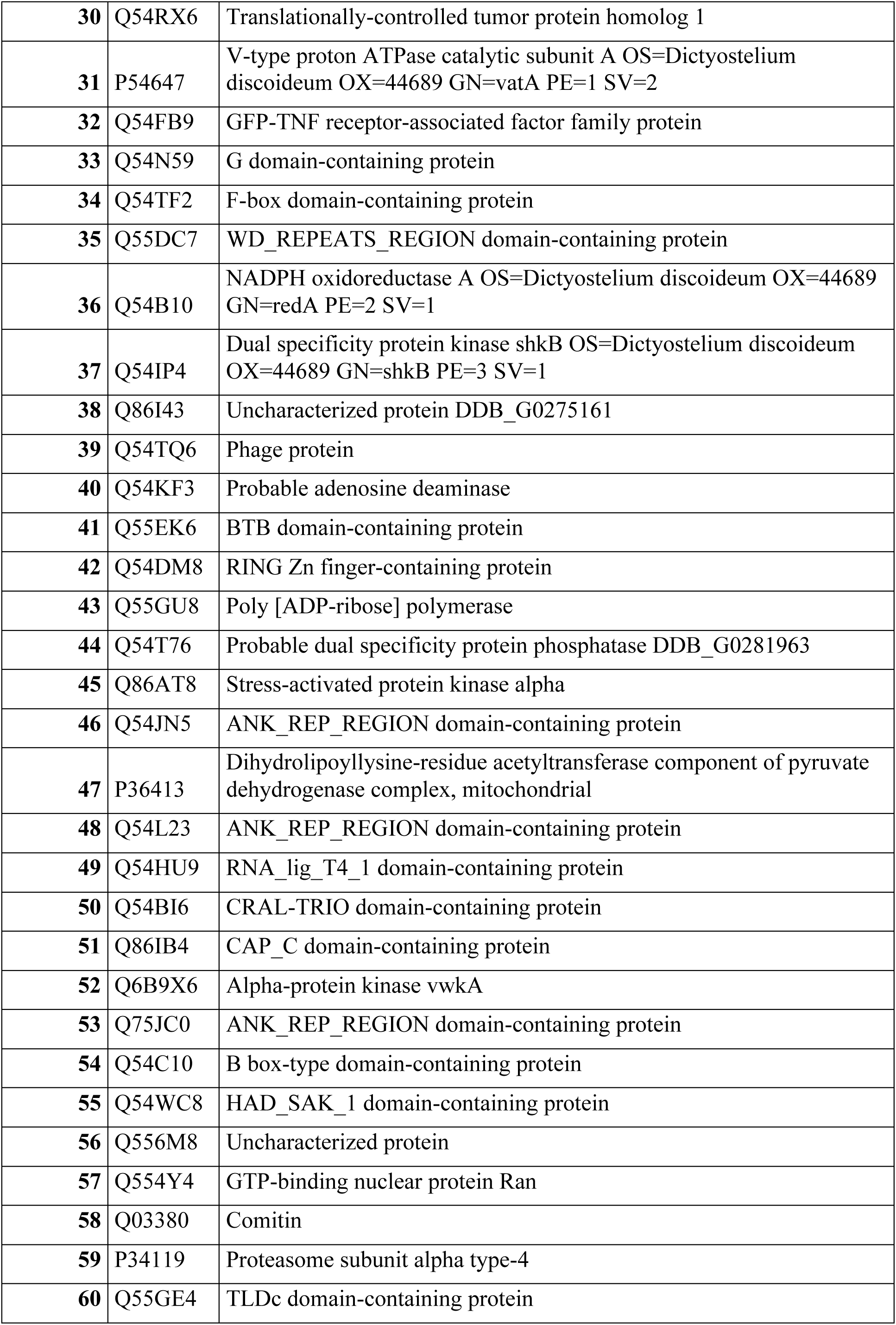
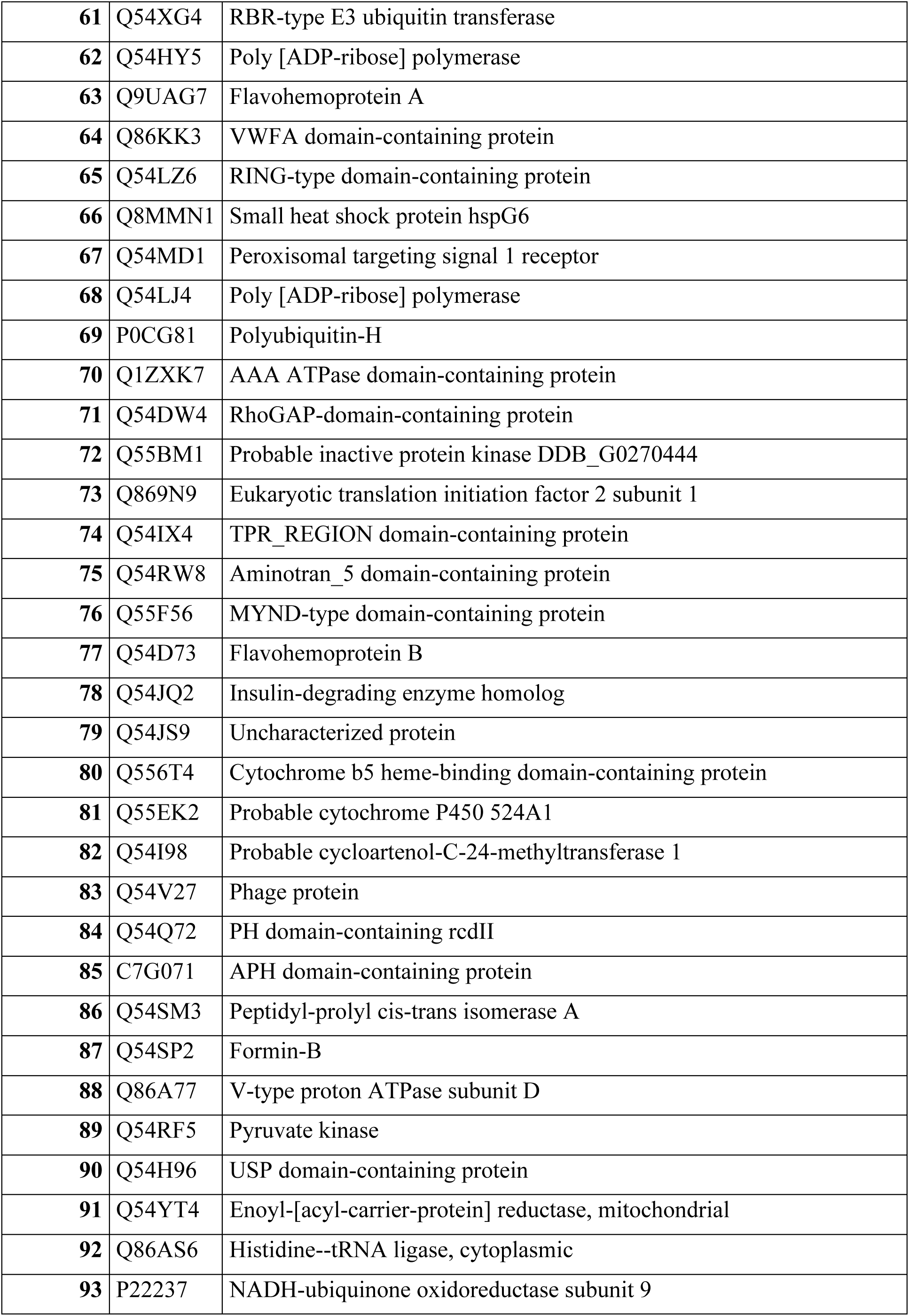
Proteins significantly enriched exclusively in TrafE and TarfE pull-downs and additional baits.

**Supplementary Table 6.** Proteins significantly enriched exclusively in Atg8a and Atg8a pull-downs and additional baits.

|  | Accession | Description |
| --- | --- | --- |
| **Exclusive TrafE-LLoMe** |  |  |
| 1 | Q54TH8 | T-complex protein 1 subunit gamma |
| 2 | Q54XF7 | Cullin-5 |
| 3 | Q86L05 | 60S ribosomal protein L10a |
| 4 | B0G0Y7 | ATP synthase-coupling factor 6, mitochondrial |
| 5 | Q54PX6 | PPM-type phosphatase domain-containing protein |
| 6 | Q75JR3 | Isocitrate dehydrogenase [NADP] cytoplasmic |
| 7 | Q9XPJ9 | ATP synthase subunit alpha, mitochondrial |
| 8 | Q86K89 | ATP synthase subunit d, mitochondrial |
| 9 | Q54F07 | S-adenosylmethionine synthase |
| 10 | Q54PH8 | 40S ribosomal protein S13 |
| 11 | P05095 | Alpha-actinin A |
| 12 | Q8MP23 | Transmembrane protein |
| 13 | Q54QR9 | Protein phosphatase 2A scaffold subunit |
| 14 | Q55CJ9 | Aldehyde dehydrogenase |
| 15 | Q75JY8 | Actin binding protein |
| 16 | Q54NU2 | Ras-related protein Rab-1D |
| 17 | Q54W38 | Glyco_hydro_63 domain-containing protein |
| 18 | Q54R45 | Translocon-associated protein subunit alpha |
| 19 | Q54TY4 | Lysine--tRNA ligase |
| 20 | A9CLV8 | ATPase subunit 4 |
| 21 | Q8T2H0 | CYFIP-related Rac1 interactor homolog |
| 22 | P54872 | Hydroxymethylglutaryl-CoA synthase A |
| 23 | Q8T2R4 | Bet_v_1 domain-containing protein |
| 24 | Q54FL2 | Ras-related protein RabG2 |
| 25 | Q54Q31 | Prohibitin-2 |
| 26 | Q86IC8 | Probable caffeoyl-CoA O-methyltransferase 2 |
| 27 | Q54X04 | Probable cysteine desulfurase, mitochondrial |
| 28 | Q54C27 | Dolichyl-diphosphooligosaccharide--protein glycosyltransferase subunit 1 |
| 29 | O00909 | ADP-ribosylation factor 1 |
| 30 | P26199 | Profilin-1 |
| **31** | Q54JD2 | Putative uncharacterized protein DDB_G0288137 |
| **32** | P24639 | Annexin A7 |
| **33** | Q01501 | Mitochondrial outer membrane protein porin |
| **34** | Q86IY4 | Thiolase |
| **35** | P32073 | GMP synthase [glutamine-hydrolyzing] |
| **36** | Q869Q8 | Probable serine carboxypeptidase CPVL |
| **37** | Q55GQ6 | Uracil phosphoribosyltransferase |
| **38** | Q54C24 | Vacuolar protein sorting-associated protein 35 |
| **39** | Q23862 | Rho-related protein racE |
| **40** | Q54LK1 | glutamine--fructose-6-phosphate transaminase (isomerizing) |
| **41** | P15064 | Ras-like protein rasG |
| **42** | Q55D79 | ECH_2 domain-containing protein |
| **43** | Q54B43 | Carm_PH domain-containing protein |
| **44** | Q54Q07 | proline--tRNA ligase |
| **45** | Q55C16 | Ubiquitin-like modifier-activating enzyme 1 |
| **46** | Q54I27 | ATP-dependent Clp protease ATP-binding subunit ClpB |
| **47** | Q54NM9 | Dolichyl-diphosphooligosaccharide--protein glycosyltransferase subunit STT3 |
| **48** | Q54PF3 | COP9 signalosome complex subunit 5 |
| **49** | P34140 | Ras-related protein Rab-1B |
| **50** | P18613 | Ras-related protein rapA |
| **51** | Q9NKX1 | Endoplasmin homolog |
| **52** | Q86L04 | TNF receptor-associated protein 1 homolog, mitochondrial |
| **53** | Q54RB9 | Calcium-binding mitochondrial carrier protein |
| **54** | P34109 | Myosin ID heavy chain |
| **55** | Q54VI3 | Glutamate dehydrogenase 2 |
| **56** | Q8T849 | CBS domain-containing protein |
| **57** | Q55F30 | proton-translocating NAD(P)(+) transhydrogenase |
| **58** | Q75JR2 | Isocitrate dehydrogenase [NADP], mitochondrial |
| **59** | P54679 | Probable plasma membrane ATPase |
| **60** | Q555Q9 | 1,4-alpha-glucan-branching enzyme |
| **61** | Q86KD1 | Cullin-associated NEDD8-dissociated protein 1 |
| **62** | Q55AD9 | ADP-ribosylation factor-like protein 5 |
| **63** | Q6RYT0 | Fructose-1,6-bisphosphatase |
| **64** | Q9NKW1 | Peroxisomal multifunctional enzyme A |
| **65** | Q54IF0 | Probable 39S ribosomal protein L21, mitochondrial |
| **66** | B0G159 | Mitochondrial substrate carrier family protein C |
| **67** | Q55CK8 | Uncharacterized protein |
| **68** | Q54QQ0 | Inosine-5'-monophosphate dehydrogenase |
| **69** | Q54VI4 | Putative glutathione S-transferase alpha-3 |
| **70** | Q54GG6 | Transmembrane protein |
| **71** | Q00766 | Glycogen phosphorylase 1 |
| **72** | Q54HL6 | COP9 signalosome complex subunit 2 |
| **73** | P08800 | UTP--glucose-1-phosphate uridylyltransferase 1 |
| **74** | Q54CA5 | S5 DRBM domain-containing protein |
| **75** | Q54RA2 | Delta-1-pyrroline-5-carboxylate dehydrogenase, mitochondrial |
| **76** | Q86CR8 | GFP-Autophagy-related protein 8 |

## References

1. Wang F, Gomez-Sintes R, Boya P. Lysosomal membrane permeabilization and cell death. Traffic. 2018;19(12):918–31.

2. Zhen Y, Radulovic M, Vietri M, Stenmark H. Sealing holes in cellular membranes. EMBO J. 2021;40(7):e106922.

3. Meyer H, Kravic B. The Endo-Lysosomal Damage Response. Annu Rev Biochem. 2024;93(1):367–87.

4. Bohannon KP, Hanson PI. ESCRT puts its thumb on the nanoscale: Fixing tiny holes in endolysosomes. Curr Opin Cell Biol. 2020;65:122–30.

5. Jukic N, Perrino AP, Humbert F, Roux A, Scheuring S. Snf7 spirals sense and alter membrane curvature. Nat Commun. 2022;13(1):2174.

6. Olmos Y. The ESCRT Machinery: Remodeling, Repairing, and Sealing Membranes. Membranes (Basel). 2022;12(6).

7. Williams JK, Ngo JM, Lehman IM, Schekman R. Annexin A6 mediates calcium-dependent exosome secretion during plasma membrane repair. Elife. 2023;12.

8. Gerke V, Gavins FNE, Geisow M, Grewal T, Jaiswal JK, Nylandsted J, Rescher U. Annexins-a family of proteins with distinctive tastes for cell signaling and membrane dynamics. Nat Commun. 2024;15(1):1574.

9. Giannini C, Ponzone L, Barroero N, Hirsch E. The interplay between phosphoinositides and ESCRT proteins. Adv Biol Regul. 2025:101126.

10. Xun J, Tan JX. Lysosomal Repair in Health and Disease. J Cell Physiol. 2025;240(5):e70044.

11. Gong YN, Guy C, Olauson H, Becker JU, Yang M, Fitzgerald P, et al. ESCRT-III Acts Downstream of MLKL to Regulate Necroptotic Cell Death and Its Consequences. Cell. 2017;169(2):286–300 e16.

12. Radulovic M, Schink KO, Wenzel EM, Nahse V, Bongiovanni A, Lafont F, Stenmark H. ESCRT-mediated lysosome repair precedes lysophagy and promotes cell survival. EMBO J. 2018;37(21).

13. Chatellard-Causse C, Blot B, Cristina N, Torch S, Missotten M, Sadoul R. Alix (ALG-2-interacting protein X), a protein involved in apoptosis, binds to endophilins and induces cytoplasmic vacuolization. J Biol Chem. 2002;277(32):29108–15.

14. Maki M, Takahara T, Shibata H. Multifaceted Roles of ALG-2 in Ca(2+)-Regulated Membrane Trafficking. Int J Mol Sci. 2016;17(9).

15. Shibata H. Adaptor functions of the Ca(2+)-binding protein ALG-2 in protein transport from the endoplasmic reticulum. Biosci Biotechnol Biochem. 2019;83(1):20–32.

16. Skowyra ML, Schlesinger PH, Naismith TV, Hanson PI. Triggered recruitment of ESCRT machinery promotes endolysosomal repair. Science. 2018;360(6384).

17. Suzuki H, Kawasaki M, Inuzuka T, Okumura M, Kakiuchi T, Shibata H, et al. Structural basis for Ca2+-dependent formation of ALG-2/Alix peptide complex: Ca2+/EF3-driven arginine switch mechanism. Structure. 2008;16(10):1562–73.

18. Vietri M, Radulovic M, Stenmark H. The many functions of ESCRTs. Nat Rev Mol Cell Biol. 2020;21(1):25–42.

19. Cambier CJ, Falkow S, Ramakrishnan L. Host evasion and exploitation schemes of Mycobacterium tuberculosis. Cell. 2014;159(7):1497–509.

20. Snyder NA, Silva GM. Deubiquitinating enzymes (DUBs): Regulation, homeostasis, and oxidative stress response. J Biol Chem. 2021;297(3):101077.

21. Sharma V, Verma S, Seranova E, Sarkar S, Kumar D. Selective Autophagy and Xenophagy in Infection and Disease. Front Cell Dev Biol. 2018;6:147.

22. Boyle KB, Randow F. The role of ’eat-me’ signals and autophagy cargo receptors in innate immunity. Curr Opin Microbiol. 2013;16(3):339–48.

23. Jamwal SV, Mehrotra P, Singh A, Siddiqui Z, Basu A, Rao KV. Mycobacterial escape from macrophage phagosomes to the cytoplasm represents an alternate adaptation mechanism. Sci Rep. 2016;6:23089.

24. Thurston TL, Wandel MP, von Muhlinen N, Foeglein A, Randow F. Galectin 8 targets damaged vesicles for autophagy to defend cells against bacterial invasion. Nature. 2012;482(7385):414–8.

25. Guallar-Garrido S, Soldati T. Exploring host-pathogen interactions in the Dictyostelium discoideum-Mycobacterium marinum infection model of tuberculosis. Dis Model Mech. 2024;17(7).

26. Xander C, Rajagopalan S, Jacobs WR, Jr., Braunstein M. The SapM phosphatase can arrest phagosome maturation in an ESX-1 independent manner in Mycobacterium tuberculosis and BCG. Infect Immun. 2024;92(7):e0021724.

27. Wong D, Bach H, Sun J, Hmama Z, Av-Gay Y. Mycobacterium tuberculosis protein tyrosine phosphatase (PtpA) excludes host vacuolar-H+-ATPase to inhibit phagosome acidification. Proc Natl Acad Sci U S A. 2011;108(48):19371–6.

28. Simeone R, Bobard A, Lippmann J, Bitter W, Majlessi L, Brosch R, Enninga J. Phagosomal rupture by Mycobacterium tuberculosis results in toxicity and host cell death. PLoS Pathog. 2012;8(2):e1002507.

29. Lopez-Jimenez AT, Cardenal-Munoz E, Leuba F, Gerstenmaier L, Barisch C, Hagedorn M, et al. The ESCRT and autophagy machineries cooperate to repair ESX-1-dependent damage at the Mycobacterium-containing vacuole but have opposite impact on containing the infection. PLoS Pathog. 2018;14(12):e1007501.

30. Meyer H, Bug M, Bremer S. Emerging functions of the VCP/p97 AAA-ATPase in the ubiquitin system. Nat Cell Biol. 2012;14(2):117–23.

31. Chen D, Fearns A, Gutierrez MG. Mycobacterium tuberculosis phagosome Ca(2+) leakage triggers multimembrane ATG8/LC3 lipidation to restrict damage in human macrophages. Sci Adv. 2025;11(13):eadt3311.

32. Crespo-Yanez X, Oddy J, Lamrabet O, Jauslin T, Marchetti A, Cosson P. Sequential action of antibacterial effectors in Dictyostelium discoideum phagosomes. Mol Microbiol. 2023;119(1):74–85.

33. Dunn JD, Bosmani C, Barisch C, Raykov L, Lefrancois LH, Cardenal-Munoz E, et al. Eat Prey, Live: Dictyostelium discoideum As a Model for Cell-Autonomous Defenses. Front Immunol. 2017;8:1906.

34. Cardenal-Munoz E, Barisch C, Lefrancois LH, Lopez-Jimenez AT, Soldati T. When Dicty Met Myco, a (Not So) Romantic Story about One Amoeba and Its Intracellular Pathogen. Front Cell Infect Microbiol. 2017;7:529.

35. Nitschke J, Hanna N, Soldati T. Dictyostelium discoideum-Mycobacterium marinum infection model: a powerful high-throughput screening platform for anti-infective compounds. Front Microbiol. 2025;16:1612354.

36. Raykov L, Mottet M, Nitschke J, Soldati T. A TRAF-like E3 ubiquitin ligase TrafE coordinates ESCRT and autophagy in endolysosomal damage response and cell-autonomous immunity to Mycobacterium marinum. Elife. 2023;12.

37. Andrews NW, Corrotte M. Plasma membrane repair. Curr Biol. 2018;28(8):R392–R7.

38. Horikawa K, Yamada Y, Matsuda T, Kobayashi K, Hashimoto M, Matsu-ura T, et al. Spontaneous network activity visualized by ultrasensitive Ca(2+) indicators, yellow Cameleon-Nano. Nat Methods. 2010;7(9):729–32.

39. Hashimura H, Morimoto YV, Hirayama Y, Ueda M. Calcium responses to external mechanical stimuli in the multicellular stage of Dictyostelium discoideum. Sci Rep. 2022;12(1):12428.

40. Perret A, Bosmani C, Leuba F, Eblighatian K, Gueho A, Raykov L, et al. Membrane microdomains are crucial for Mycobacterium marinum EsxA-dependent membrane damage, escape to the cytosol, and infection. Sci Adv. 2026;12(1):eady0812.

41. Garrity AG, Wang W, Collier CM, Levey SA, Gao Q, Xu H. The endoplasmic reticulum, not the pH gradient, drives calcium refilling of lysosomes. Elife. 2016;5.

42. Scheffer LL, Sreetama SC, Sharma N, Medikayala S, Brown KJ, Defour A, Jaiswal JK. Mechanism of Ca(2)(+)-triggered ESCRT assembly and regulation of cell membrane repair. Nat Commun. 2014;5:5646.

43. Shukla S, Chen W, Rao S, Yang S, Ou C, Larsen KP, et al. Mechanism and cellular function of direct membrane binding by the ESCRT and ERES-associated Ca(2+)-sensor ALG-2. Proc Natl Acad Sci U S A. 2024;121(9):e2318046121.

44. Subramanian L, Crabb JW, Cox J, Durussel I, Walker TM, van Ginkel PR, et al. Ca2+ binding to EF hands 1 and 3 is essential for the interaction of apoptosis-linked gene-2 with Alix/AIP1 in ocular melanoma. Biochemistry. 2004;43(35):11175–86.

45. Aubry L, Mattei S, Blot B, Sadoul R, Satre M, Klein G. Biochemical characterization of two analogues of the apoptosis-linked gene 2 protein in Dictyostelium discoideum and interaction with a physiological partner in mammals, murine Alix. J Biol Chem. 2002;277(24):21947–54.

46. Talukder MSU, Pervin MS, Tanvir MIO, Fujimoto K, Tanaka M, Itoh G, Yumura S. Ca(2+)-Calmodulin Dependent Wound Repair in Dictyostelium Cell Membrane. Cells. 2020;9(4).

47. Kitaura Y, Matsumoto S, Satoh H, Hitomi K, Maki M. Peflin and ALG-2, members of the penta-EF-hand protein family, form a heterodimer that dissociates in a Ca2+-dependent manner. J Biol Chem. 2001;276(17):14053–8.

48. Cooper ST, McNeil PL. Membrane Repair: Mechanisms and Pathophysiology. Physiol Rev. 2015;95(4):1205–40.

49. Clapham DE. Calcium signaling. Cell. 2007;131(6):1047–58.

50. Dubois C, Prevarskaya N, Vanden Abeele F. The calcium-signaling toolkit: Updates needed. Biochim Biophys Acta. 2016;1863(6 Pt B):1337–43.

51. Shukla S, Larsen KP, Ou C, Rose K, Hurley JH. In vitro reconstitution of calcium-dependent recruitment of the human ESCRT machinery in lysosomal membrane repair. Proc Natl Acad Sci U SA. 2022;119(35):e2205590119.

52. Corkery DP, Castro-Gonzalez S, Knyazeva A, Herzog LK, Wu YW. An ATG12-ATG5-TECPR1 E3-like complex regulates unconventional LC3 lipidation at damaged lysosomes. EMBO Rep. 2023;24(9):e56841.

53. Augenstreich J, Arbues A, Simeone R, Haanappel E, Wegener A, Sayes F, et al. ESX-1 and phthiocerol dimycocerosates of Mycobacterium tuberculosis act in concert to cause phagosomal rupture and host cell apoptosis. Cell Microbiol. 2017;19(7).

54. Conrad WH, Osman MM, Shanahan JK, Chu F, Takaki KK, Cameron J, et al. Mycobacterial ESX-1 secretion system mediates host cell lysis through bacterium contact-dependent gross membrane disruptions. Proc Natl Acad Sci U S A. 2017;114(6):1371–6.

55. Perret A, Michard C, Moreau D, Soldati T. Tiny organisms, big advantages: Exploring membrane damage responses using Dictyostelium discoideum in high throughput single cell analyses. In: S. Linder CW, editor. Live Biological Imaging Across Scales: Wiley 2026.

56. Tinevez JY, Perry N, Schindelin J, Hoopes GM, Reynolds GD, Laplantine E, et al. TrackMate: An open and extensible platform for single-particle tracking. Methods. 2017;115:80–90.

57. Hagedorn M, Soldati T. Flotillin and RacH modulate the intracellular immunity of Dictyostelium to Mycobacterium marinum infection. Cell Microbiol. 2007;9(11):2716–33.

58. Arafah S, Kicka S, Trofimov V, Hagedorn M, Andreu N, Wiles S, et al. Setting up and monitoring an infection of Dictyostelium discoideum with mycobacteria. Methods Mol Biol. 2013;983:403–17.

59. Andreu N, Zelmer A, Fletcher T, Elkington PT, Ward TH, Ripoll J, et al. Optimisation of bioluminescent reporters for use with mycobacteria. PLoS One. 2010;5(5):e10777.

60. Hanna N, Burdet F, Melotti A, Bosmani C, Kicka S, Hilbi H, et al. Time-resolved RNA-seq profiling of the infection of Dictyostelium discoideum by Mycobacterium marinum reveals an integrated host response to damage and stress. bioRxiv 2019.

